# Ventral striatal astrocytes contribute to reinforcement learning

**DOI:** 10.1101/2025.10.20.683205

**Authors:** Julia Pai, Fatih Sogukpinar, Kei Ogasawara, Garrett J Smith, Francesca R Fiocchi, Yanchao Dai, Yifan Wu, Michael J Frank, ShiNung Ching, Federica Lucantonio, Thomas Papouin, Marco Pignatelli, Naoki Hiratani, Ilya E Monosov

## Abstract

Astrocytes influence synaptic plasticity and neuronal function through astrocytic calcium dynamics (ACD). However, astrocytes’ contribution to cognitive operations like reinforcement learning (RL) remains unclear. To examine this, we trained mice on a RL dependent probabilistic decision-making task. We attenuated ACD across distinct striatal regions, and found ACD attenuation specifically in ventral striatum (VS) increased decision noisiness and impaired reward-guided choice performance. This effect was largely due to a reduction in “win-stay” behavior. Using in-vivo calcium imaging, we found that VS ACD correlated with reward prediction errors (RPEs). Furthermore, these trial-by-trial ACD fluctuations predicted trial-by-trial choice variability. In-silico lesions of a biologically constrained circuit model suggest that astrocytes could regulate behavioral variability by sharing RPE signals across populations of striatal neurons. Together, these results suggest that VS astrocytes contribute to cortico-striatal functions to mediate decision noisiness.

## Introduction

Reinforcement learning (RL) describes the processes by which an agent or organism learns, through trial and error, to predict and take actions that maximize cumulative reward. RL is thought to be dependent on the striatum, with striatal subregions thought to play distinct roles in learning values of actions or policies (Burton et al., 2015; Cox & Witten, 2019; Ito & Doya, 2011; H. Kim et al., 2009). Critically, these striatal circuits can provide key links between theoretical descriptions of learning and their biological implementation. For example, dopaminergic signaling within the striatum has been shown to track key learning signals predicted by formal RL frameworks (Bayer & Glimcher, 2005; H. R. Kim et al., 2020; Schultz, 1998), providing a clear link between neural activity and theoretical descriptions of learning processes.

There is mounting evidence that astrocytes are active participants within these same circuits. They respond to neuromodulatory signals, including dopamine (Corkrum et al., 2020; Lazaridis et al., 2024), and contribute to synaptic function and ongoing neural activity through multiple functional knobs, including neurotransmitter reuptake and recycling, control of ionic concentrations, metabolic regulation, and transmitter release (Araque et al., 1999a; Corkrum et al., 2020; De Pittà et al., 2016; Guttenplan et al., 2025; Haydon & Carmignoto, 2006; Henneberger et al., 2010; Oliveira et al., 2015; Petzold & Murthy, 2011). Despite the well-established roles that astrocytes have in physiological control of neuronal functions, whether and how astrocytes shape RL behavior is poorly defined.

Astrocytes have several key functional properties that could allow them to uniquely support or contribute to RL processes. Astrocytic modulation of neuronal activity and synaptic functions is thought to be dependent on astrocytic calcium dynamics (ACD) and can occur on a spectrum of spatiotemporal scales: relatively fast and locally (sub-second, synapse-level) and relatively slowly and more globally (hours-days, population-level) (Adamsky et al., 2018; Benoit et al., 2025; Di Castro et al., 2011; Fujii et al., 2017; Panatier et al., 2011; Scemes & Giuame, 2006; Stobart et al., 2018). ACD are thought to reflect ongoing circuit activity, including synaptic activity, neuronal firing and neuromodulator release, such dopamine and norepinephrine (Allen, 2014; Bai et al., 2024; Bekar et al., 2008; Charles et al., 1991; Corkrum et al., 2020; Cornell-Bell et al., 1990; Di Castro et al., 2011; Khakh & McCarthy, 2015; Lefton et al., 2025; Ma et al., 2016; Mu et al., 2019; Panatier et al., 2011; Schipke et al., 2008; Sofroniew & Vinters, 2010). In addition, astrocyte networks possess unique properties that distinguish them from neurons. For example, astrocytes are thought to tile the brain in non-overlapping domains (Bushong et al., 2002; Halassa et al., 2007; Ogata & Kosaka, 2002), with each astrocyte closely interacting with hundreds of thousands of synapses (Oberheim et al., 2009), and they are extensively gap-junction coupled into functional networks (Cooper et al., 2025; Dermietzel et al., 1991).

These abilities might confer astrocytes the capacity to regulate neuronal computations over a wide spatial and temporal scale in response to local computations. That is, astrocytes could uniquely contribute to neuronal circuits by helping to address a key function required for cognitive operations: adaptively regulating variability within local circuits to produce stable representations to guide behavior. This possibility that astrocytes participate in averaging and sharing information across nearby neurons (also known as input sharing) has strong analogs in artificial intelligence, where such properties are crucial for multiple forms of learning algorithms (Fukushima, 1980; Kipf & Welling, 2017; Krizhevsky et al., 2012; LeCun et al., 1989, p. 19) and neuromorphic architectures (Becker et al., 2022; Gong et al., 2024; Irizarry-Valle & Parker, 2015; Kozachkov et al., 2023; Nazari et al., 2015; G. Tang et al., 2019); in such instantiations, input sharing is not an active non-linear component of algorithmic implementation per se, but directly contributes to it.

To start to assess whether and how astrocytes contribute to RL, we attenuated ACD across different subregions of striatum while animals performed a RL-dependent probabilistic reversal task. Importantly, we used a computationally tractable task that allowed us to model the effects of ACD attenuation within a normative learning framework. We found that attenuation of ACD in dorsomedial striatum (DMS) and dorsolateral striatum (DLS) had relatively little impact on key reinforcement learning parameters, such as on learning rate and inverse temperature (decision noisiness). In contrast, VS ACD attenuation decreased inverse temperature (increasing decision noisiness) and impaired reward-guided choice, specifically by reducing win-stay behavior. Using fiber photometry to monitor ACD during RL revealed that VS ACD reflected model inferred reward prediction errors (RPEs), for many seconds after the outcome, until the start of the next trial. We built a biologically constrained circuit model of neuron-astrocyte interactions to assess how astrocyte encoding of RPE learning signals could support RL-guided decision-making. In both in vivo photometry recordings and circuit model, trial-by-trial ACD fluctuations predicted trial-by-trial choice variability, particularly in win-stay behavior. In-silico experiments suggested that choice performance and win-stay behavior could be mediated by astrocytes averaging and sharing dopaminergic RPE inputs across medium spiny neuron (MSN) populations. These findings highlight a role of ACD in specific RL computations and generate novel hypotheses for how astrocytes contribute to value-based decision-making and cognition.

## Results

### Attenuating astrocyte calcium dynamics (ACD) in ventral striatum impairs probabilistic decision-making in a reinforcement learning task

To begin to assess the role of striatal ACD in reinforcement learning, we first trained mice to perform a non-stationary probabilistic reversal learning task (‘bandit’ task) commonly used to study reinforcement learning (RL) algorithms in animals and artificial systems (Bertsekas & Tsitsiklis, 1996; Costa et al., 2015; Grossman et al., 2022; Ito & Doya, 2011; Sutton & Barto, 2018). At the start of each trial, freely moving mice received a visual start cue that informed them that they could make a choice by performing a left or right nose poke into an in-home cage behavioral device (FED3, (Matikainen-Ankney et al., 2021)). These two ports were associated with two varying reward probabilities: either the left one was 80% reward, and the right one was 20% reward, or the left one was 20% reward, and the right one was 80% reward (Figure 1A). The trials were blocked so that the contingency (higher reward probability in left arm or right arm) was stable for 20-30 rewarded trials and then switched, un-cued to the mouse.

**Figure 1.**
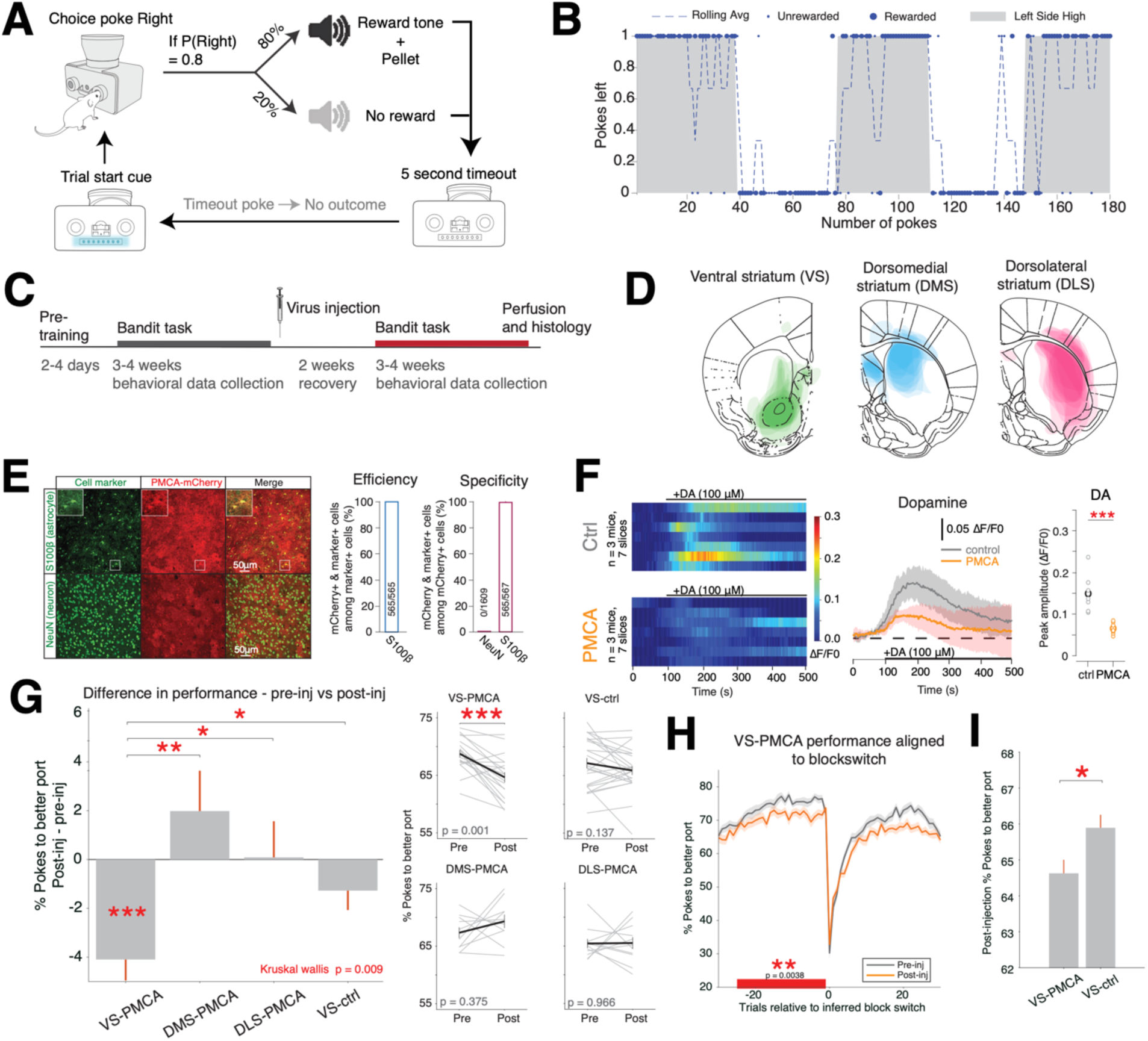
Attenuating astrocyte calcium dynamics (ACD) across striatum subregions leads to performance deficits preferentially in VS. A. Trial diagram of the probabilistic decision-making task. Animals are trained to poke for rewards via an in-home cage operant behavior device, which lights up when trials are available. Animals initiate choices by making a nose poke into the left or right port. Ports have either 80% or 20% probabilities of being rewarded. If rewarded, a reward tone is presented, and a pellet is delivered. If un-rewarded, a different no-reward tone is presented. After each choice, the device enters a 5 second time out, before another trial becomes available. B. Behavior from one example session. Dots on the top and bottom indicate choices the animal made to the left and right and the dashed line shows the rolling average of left choices. C. Experimental timeline. D. Average PMCA virus expression localization in VS, DMS, and DLS cohorts. E. Immunohistochemistry (IHC) showing the colocalization of PMCA expression (mCherry) with an astrocyte marker (S100β) in the VS. By contrast, there was no expression of PMCA in neurons, identified with the neuronal marker NeuN. Quantification of efficiency and specificity of PMCA expression in astrocytes is shown on the right. Numbers in the bar graphs indicate the number of cells quantified as indicated on the axis title. F. Left: heatmaps of VS astrocyte calcium responses to bath application of dopamine (DA) in control (top heatmap) or PMCA (bottom heatmap) slices. DA application occurs at 100s. Each row represents one slice’s responses to one application; black lines on top indicate the time scale of DA application. Middle: average traces of control or PMCA ACD responses to DA application. Shaded area is 95% confidence interval. Right: comparison of the peak responses to DA (maximum response between 100-300 s) between control and PMCA slices. G. Change in decision-making performance (quantified as % of choices to the port with better reward probability) pre- vs post-injection for VS-PMCA (n = 19), DMS-PMCA (n = 10), DLS-PMCA (n = 11), and VS-ctrl (n = 24) cohorts. Left: Average change in performance for each cohort across animals; error bars are S.E.M. across animals. Comparisons across groups are Wilcoxon rank sum. Right subplots for each cohort: Gray lines are individual animals’ performance; black lines are average across animals. Error bars are S.E.M. across animals. Within-animal comparisons for each cohort are Wilcoxon signed-rank. H. Decision-making performance for VS-PMCA pre-injection sessions (black) and post-injection sessions (orange) aligned to the time of un-cued block switches, as a Bayesian ideal observer would infer them (Bartolo & Averbeck, 2020). Aligning choice performance to ideal observer-inferred block switches corrects for the fact that true switch points can be obscured by stochastic reward delivery. For instance, if the right port becomes the 80% reward side but the left port continues to deliver rewards by chance, it may take several trials before the change is detectable. This analysis accounts for such misleading trial sequences and allows for more accurate comparison of behavioral dynamics across blocks. Choice performance was significantly different between pre- and post-injection behavior assessed in a 25-trial window prior to inferred block switches (gray bar, Wilcoxon sign rank test). Error bars are S.E.M. across animals (n = 19). I. Decision-making performance for post-injection VS-PMCA (n = 389 sessions) and post-injection VS-ctrl mice (n = 485 sessions). Post-injection VS-PMCA mice show significantly worse choice performance compared to post-injection VS-ctrl mice (p = 0.0236, Wilcoxon rank sum test). *= p<0.05, ** = p<0.01, ***=p<0.001.

Although structurally simple, the non-stationarity and stochasticity of reward schedules impose continuous demands on subjects’ value updating and reward prediction error processing. This constrained design enabled precise, trial-by-trial quantification of latent RL variables with modeling frameworks (i.e., RPEs, learning rates, and decision parameters) (Bertsekas & Tsitsiklis, 1996; Cohen, 2008; Costa et al., 2016; Dalton et al., 2014; Daw et al., 2006; Frank et al., 2004; Grossman et al., 2022; Humphries et al., 2012; Ineichen et al., 2012; Metha et al., 2020; Montague et al., 1996; Sutton & Barto, 2018), while enabling high throughput training of many mice within days (Supplemental Figure 1A). We could then assess how these factors changed after ACD attenuation across striatal subregions.

Mice chose the higher probability port most of the time (mean = 67.4%, SD = 7.6%, Supplemental Figure 1C), on par with previously reported performance of mice in comparably structured bandit tasks (Hattori et al., 2019; Iigaya et al., 2018). Behavior from one example session is shown in Figure 1B. After mice were trained on the task, we attenuated ACD across different subregions of striatum to probe its role in supporting RL and probabilistic decision-making. Rodent striatum is often subdivided into ventral striatum (VS; here we use it to refer to nucleus accumbens), dorsomedial striatum (DMS), and dorsolateral striatum (DLS). Striatal subregions receive input from different cortical, thalamic, and dopaminergic regions (with some overlap between subregions; Burton et al., 2015; Haber, 2003; Humphries et al., 2012; Parent & Hazrati, 1995), and are known to contribute differentially to motivated behavior, decision-making, and RL (Burton et al., 2015; Cox & Witten, 2019; Ito & Doya, 2011; H. Kim et al., 2009; H. Tang et al., 2022). To our knowledge, few studies have systematically assessed subregion-specific contributions to performance in a probabilistic reversal learning task like ours (but see (Ito & Doya, 2015; Parker et al., 2016a)), let alone the role of astrocytes across subregions. We therefore began to screen how ACD contributes to RL by attenuating ACD in VS, DMS, and DLS in task-trained mice and quantifying their behavior before and after ACD attenuation.

We injected separate cohorts of mice bilaterally in VS (n = 19), DMS (n = 10), and DLS (n = 11) with AAV5-gfaABC1D-PMCA-mCherry (Figure 1C, 1D) (Yu et al., 2018). We also injected a cohort of mice with control virus (AAV5-gfaABC1D-tdTomato, or AAV5-GFAP104-mCherry) in VS (n=24). We allowed 2 weeks after virus injection for the viral construct to adequately express (Figure 1C, (Reimsnider et al., 2007)). Viral injections resulted in astrocyte-specific expression of plasma membrane Ca2+ ATPase (PMCA), which constitutively extrudes cytosolic Ca2+ and functionally attenuates astrocytic calcium transients (Yu et al., 2018). As expected, colocalization with the astrocyte marker S100β but lack of colocalization with NeuN indicated that PMCA was expressed selectively in astrocytes and not in neurons (Figure 1E).

We used *ex vivo* calcium imaging to validate that PMCA expression effectively attenuated ACD in ventral striatum in response to bath application of dopamine, a key neuromodulator for RL in the striatum that is also known to drive striatal astrocyte calcium (Corkrum et al., 2020; Haber, 2003)) (Figure 1F). Cell-wide, peak Ca^2+^ response amplitudes to DA were significantly attenuated in PMCA slices compared to control (Figure 1F). As expected, and in line with prior studies in the striatum (Nimitvilai-Roberts et al., 2021; Yu et al., 2018), the effect of PMCA on ACD was not specific to dopamine but effectively reduced astrocyte overall responsiveness, including to norepinephrine (Supplemental Figures 2B-G).

To begin to assess whether and how ACD contributes to RL, we compared decision performance (the percent of choices mice made to the better port) before versus after virus injection. Various prior manipulations across different subregions of striatum have been shown to have heterogeneous effects on decision-making behavior (Beeler et al., 2010; Dalton et al., 2014; Kwak & Jung, 2019). We found that PMCA-induced ACD attenuation in VS, but not in DMS, DLS, or control-injected VS mice, led to a significant decrease in choice performance in the probabilistic reversal task (Figure 1G). This effect can be further visualized when trial-by-trial choice performance is aligned to the time of un-cued block switches (Figure 1H, Supplemental Figure 3). Furthermore, the choice performance of post-injection VS-PMCA mice was significantly worse than post-injection performance of VS-ctrl mice (Figure 1I).

Next, we assessed whether mice’s win-stay or lose-switch strategies changed due to ACD attenuation. Win-stay/lose-switch strategies are simple, outcome-dependent decision-making heuristics in which subjects tend to repeat a choice after a rewarded outcome (“win-stay”) and switch after a non-rewarded outcome (“lose-switch”). Win-stay/lose-switch strategies have been widely studied to understand the mechanisms of value updating and action selection in reinforcement learning paradigms, both in animals and humans (Daw et al., 2006; Lee et al., 2012; Robbins, 1952). Because post-reward and post-non-reward strategy can arise from different algorithmic and circuit-level implementations, this analysis could provide deeper understanding of the relationship of ACD and RL performance.

We found that mice in the VS-PMCA cohort, but not other cohorts, exhibited a significant decrease in win-stay choices post-injection compared to pre-injection (Figure 2A). However, there was no significant difference in lose-switch behavior in any of the cohorts (Figure 2B).

**Figure 2.**
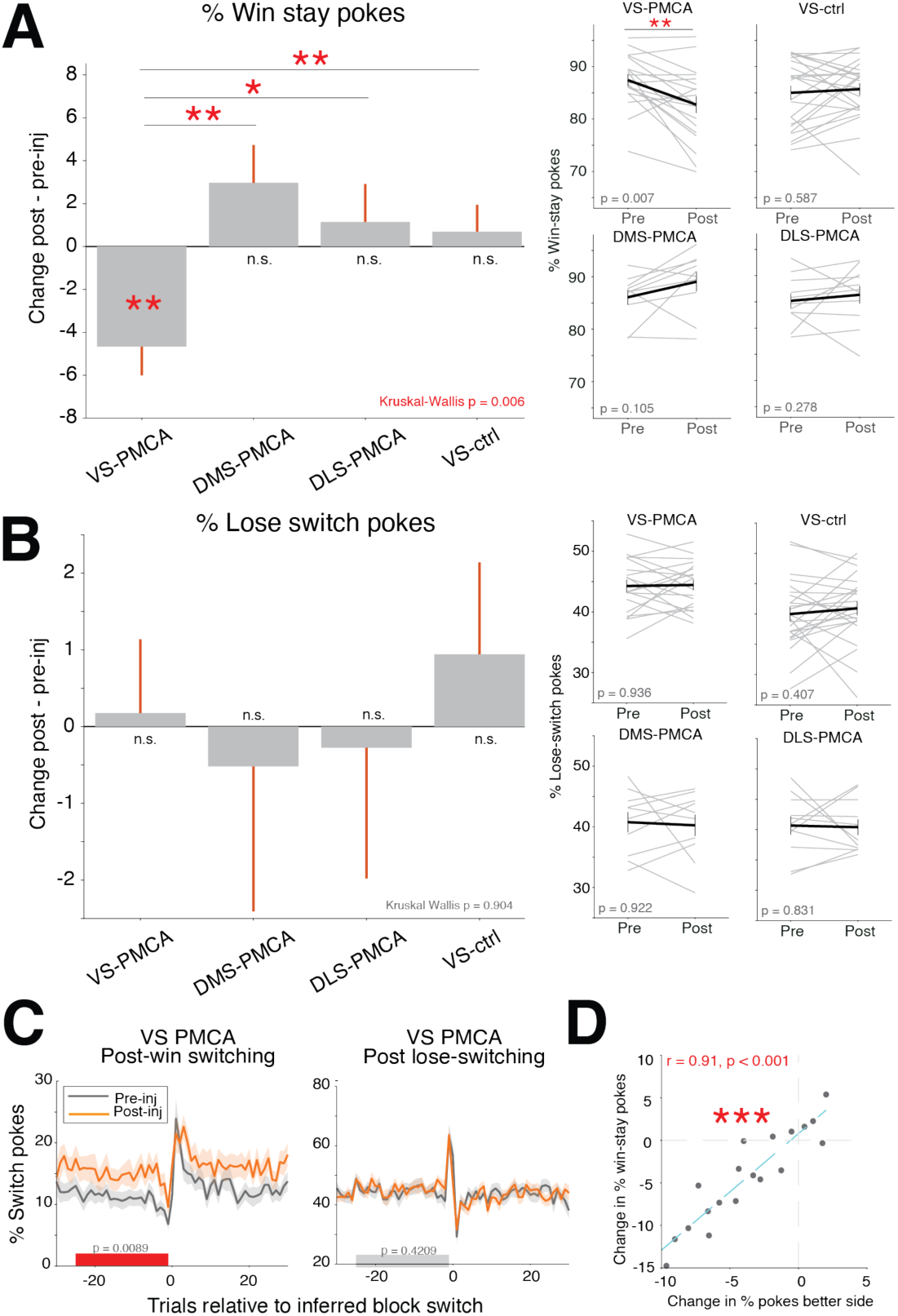
Performance deficits in mice with attenuated VS ACD are correlated with decreases in win-stay behavior. A. Change in win-stay pokes (quantified as % of choices after rewarded trials where the mouse switches relative to the previous choice) pre- vs post-injection for VS-PMCA, DMS-PMCA, DLS-PMCA, and VS-ctrl cohorts. Left: Average change in win-stay choices for each cohort across animals; error bars are S.E.M. across animals. Comparisons across groups are Wilcoxon rank sum. Right: Gray lines are individual animals’ performance; black lines are average across animals. Error bars are S.E.M. across animals. Within-animal comparisons for each cohort are Wilcoxon signed-rank. B. Change in lose-stay pokes (quantified as % of choices after unrewarded trials where the mouse repeats the same action relative to the previous poke) pre- vs post-injection across cohorts; conventions follow A. C. Average post-win switching (left) and post-lose switching (right) behavior aligned to inferred block switches (see Methods) for VS-PMCA mice. Post-win, but not post-lose, switching was significantly higher for PMCA mice in a 25-trial window prior to inferred block switches (gray bar, Wilcoxon sign rank test). Error bars are S.E.M. over animals. D. Pre- vs post-injection change in performance correlated against change in win-stay pokes for the VS-PMCA cohort; each dot indicates one mouse. There is a strong and significant correlation between change in performance and change in win-stay on an animal-by-animal basis (Pearson’s correlation).

Consistent with the win-stay effect, mice in the VS-PMCA cohort exhibited significantly more switching behavior following rewarded outcomes and not after unrewarded outcomes (Figure 2C). This effect was not observed in DMS-PMCA, DLS-PMCA mice, or VS-control mice (Supplemental Figure 3C-D). Furthermore, the change in VS-PMCA mice correct choice rate was strongly and significantly correlated with the change in their win-stay behavior on a mouse-by-mouse basis (Figure 2D).

Our data thus far suggests that attenuating ACD in VS impairs performance by altering decision making strategies relative to prior rewards. This suggests that attenuating ACD in VS influences RL-related computation in the VS circuit. However, VS is known to be highly involved in motivation, appetitive behavior, and reward-seeking (Haber & Knutson, 2010; Kelley, 2004). In order to control for the possibility that these decision-making deficits in the VS-PMCA cohort were due to motivational, hedonic, or motor effects brought on by PMCA ACD attenuation, or surgery-related damage to VS, we performed additional analyses and behavioral tests on VS-PMCA and VS-ctrl mice.

VS-PMCA-induced changes in decision-making were not due to general changes in motivation to initiate trials, motor or impulsivity measures (Supplemental Figures 1, 6, 7). Similarly to other cohorts, VS-PMCA mice initiated more trials after injection, indicating that they did not have reduced motivation to perform the task (Supplemental Figure 1D, E). VS-PMCA mice did not show significant differences in behavior from VS-ctrl mice in a progressive ratio task, a commonly used assay related to reward consumption and willingness to expend effort for reward (Johnson et al., 2022; Richardson & Roberts, 1996; Stewart, 1975) (Supplemental Figure 6). VS-PMCA mice largely did not show differences from VS-ctrl mice in a battery of behavioral tests commonly used to assay sensorimotor, cognitive, motivational, or emotion-regulation related phenotypes (Supplemental Figure 7). In summary, decision-making deficits in VS-PMCA mice were not directly attributable to sensorimotor functions, to memory performance changes, to general motivation, or to the surgery itself.

### ACD attenuation in VS increases decision noisiness in reinforcement learning

To better understand how decision-making algorithms are altered in mice with attenuated ACD in VS, we aimed to quantitatively describe processes underlying pre-and post-injection behavior using a model of decision-making behavior. We compared several models, including multiple variations of win-stay lose-shift and Q-learning models (Supplemental Figure 8, Table 3 (Bertsekas & Tsitsiklis, 1996; Sutton & Barto, 2018)). A version of Q-learning model incorporating action kernels provided the most parsimonious fit to mice’s behavior, far outperforming probabilistic win-stay lose-shift models in model comparison. This action kernel has been previously incorporated into RL models of rodent behavior to capture decision hysteresis (Findling et al., 2025; Ito & Doya, 2009; Metha et al., 2020; Parker et al., 2016b).

This RL agent (Figure 3A) maintains a trial-by-trial decision variable (*D_t_*) which consists of: *Q^left^*and *Q^right^* values (also called action values, which track the reward-associated value of left and right), *K^left^* and *K^right^* (action kernels, which track previous action history), and *b* (a side bias term). The weighting terms *β_q_* (also known as inverse temperature) and *β_k_* determine how sensitive choices are to value differences in Q- and K-values, respectively. To make a choice, these values are passed through a decision function (or a *decision process*) that transforms them into action probabilities. After each choice and outcome, the model computes a reward prediction error (RPE), which is the difference between the received reward and the expected value of the chosen action. This RPE is used to update the Q-value of the chosen action, scaled by the learning rate parameter *α_q_*. K-values of both chosen and unchosen options are also updated, independent of the outcome, scaled by the action kernel learning rate parameter *α_k_*. An example session with model-predicted choice probabilities is shown in Figure 3B.

**Figure 3.**
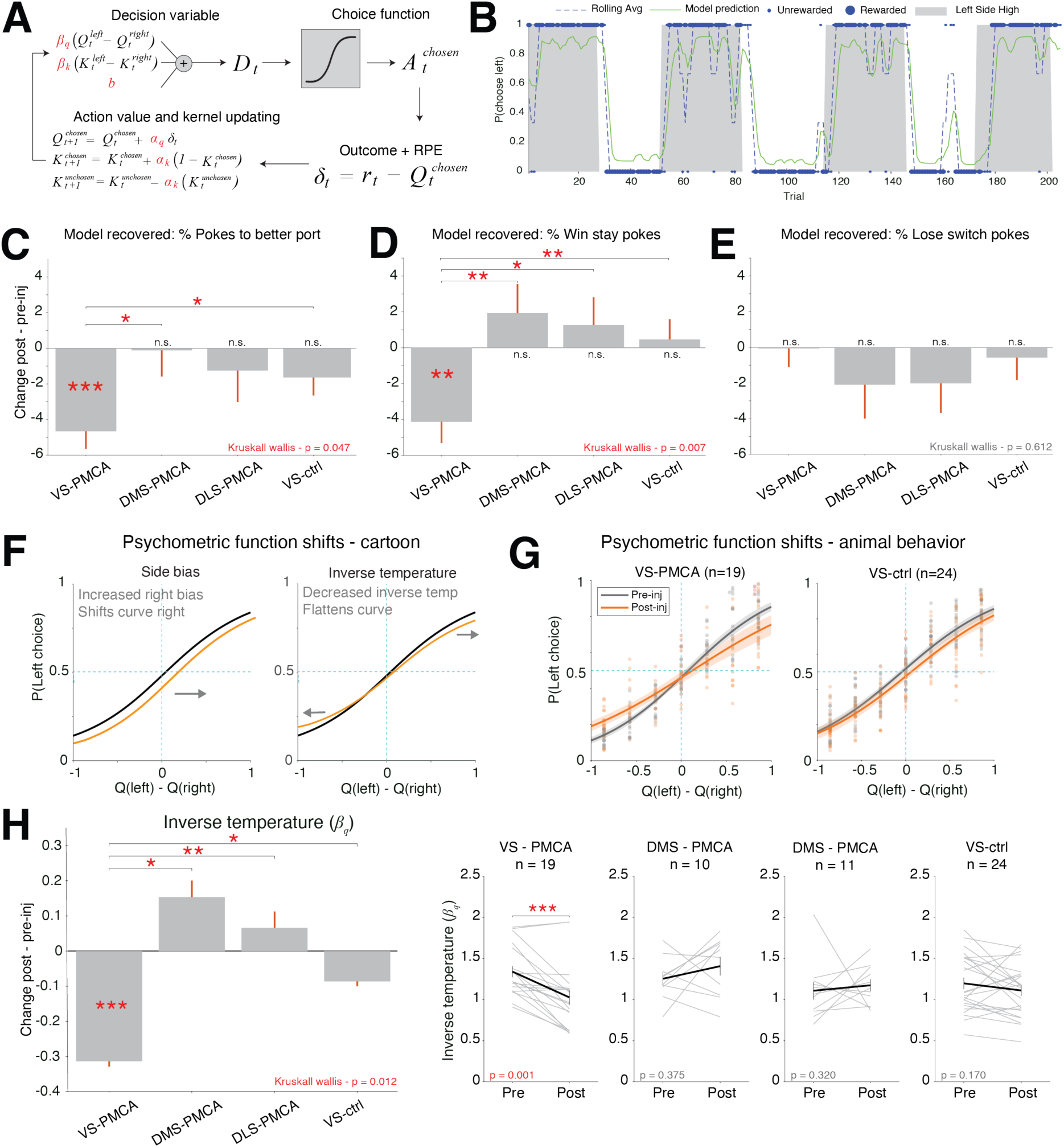
RL modelling of behavioral noise following VS ACD attenuation. A. Q-learning reinforcement learning (RL) model. Action values for left and right are stored in *Q^left^* and *Q^right^* for each trial; how deterministically Q-value differences affect choice is weighted by *β_q_* (inverse temperature). *K^left^* and *K^right^* store previous action history values (action kernels); how deterministically K-value differences affect choice is weighted by *β_k_*. *b* is a side bias term. These values are summed and passed into a sigmoidal choice function, which probabilistically outputs a left or right choice. A reward prediction error is calculated based on the choice decision and the current value of the action taken. The RPE is used to update the Q-value of the chosen action; the rate of value updating is weighted by the learning rate *α_q_*. Independently of RPE, both the K-value of the chosen side is strengthened and the K-value of the unchosen side is weakened; the rate of this updating is weighted by the action kernel learning rate *α_k_.* Variables in red indicate free parameters fitted for each animal. B. Behavioral data from one example session (conventions following Figure 1B) and RL model fit. The dashed blue line indicates the rolling average of the mouse’s real left decisions, and the green line indicates the model prediction of probability of left choice. R-squared for this session = 0.559. C – E. Model validations for the Q-learning model. To make sure the fitted Q-learning model is able to capture the key behavioral aspects in the data, we simulated behavioral data using the model parameters obtained from model fits to mice’s behavior pre- and post-injection. The model captured the same key behavioral effects seen in real mice’s data. VS-PMCA model simulated data, but not other cohorts, showed a significant decrease in choice performance and win-stay rate, but no change in lose-stay rate after injection. Statistical tests are sign-rank within cohort and rank sum across cohorts; error bars are S.E.M. across animals. F. Cartoon psychometric functions and how they are affected by shifts in inverse temperature, side bias, and lapse rate. Black indicates ‘baseline’, orange indicates function after shift in specific parameters. X-axis is difference in Q-value between left and right options; Y-axis is agent’s probability of choosing the left option. Note that a change in slope of the psychometric curve corresponds to a change in the *β_q_* parameter, where a shallower curve (lower slope) corresponds to a more random choice (lower *β_q_*), and a steeper curve (higher slope) corresponds to a more deterministic choice (higher *β_q_*). On the other hand, a shift of the psychometric curve left or right away from the indifference point (x = 0.0, y = 0.5) results from the side bias term, with an increase in the side bias term shifting the curve towards the right. G. Psychometric curves for VS-PMCA and VS-ctrl mice. Dots are individual animals’ left choice probabilities across different inferred Q-value differences; lines are average fitted sigmoid curves for pre-injection (black) and post-injection (orange) sessions across animals. Error bars = S.E.M. across animals. H. Change in model fit inverse temperature pre- vs post-injection for the different cohorts. Inverse temperature was significantly decreased for VS-PMCA but not other cohorts, and this effect was significantly different between VS-PMCA and other cohorts. Left: average change in inverse temperature for each cohort across animals; error bars are S.E.M. across animals. Comparisons across groups are Wilcoxon rank sum. Right: Gray lines are individual animals’ inverse temperature parameter fits; black lines are average across animals. Error bars are S.E.M. across animals. Within-animal comparisons for each cohort are Wilcoxon signed-rank. Across group comparison is Wilcoxon rank sum (same test as in right subplot between VS-PMCA and VS-ctrl).

We first assessed whether the fitted Q-learning model captures the key statistics of animal behavior. Data simulated with model fitted parameters of the different cohorts recreated the key behavioral effects that we saw in real animals’ data: namely, VS-PMCA, but not other cohorts, showed significantly decreased choice performance and win-stay choices, with no change in lose-switch choices (Figures 3C-E).

We next assessed psychometric curves for mice’s behavior before and after virus injection. The shape of these curves, which here visualize agents’ choice patterns against the underlying inferred action values, can be used to infer the decision process used to transform internal estimates of value into actual choices (Ballesta et al., 2020; Britten et al., 1996; Gold & Ding, 2013). Figure 3F shows cartoon diagrams of how changes in the decision process terms *β_q_* (inverse temperature) and side bias could change the shape of the psychometric function relative to inferred Q-value differences. (Note that *α_q_* affects the rate of Q-value updating but does not explicitly shift choice functions relative to stored Q-values). When we plotted VS-PMCA and VS-ctrl mice’s choice probabilities against inferred action values, we saw a post-injection shift in the average psychometric function that resembled a decrease in inverse temperature for VS-PMCA mice. The psychometric functions of VS-ctrl (Figure 3G) and DMS-PMCA and DLS-PMCA mice (Supplemental Figure 9) did not show such a shift.

Indeed, detailed analyses of the model parameters showed that inverse temperature (*β_q_*) significantly decreased for VS-PMCA but not other cohorts after virus injection (Figure 3H). A decreased inverse temperature indicates less weighting of learned value estimates in the decision process, leading to more stochastic (or ‘noisy’/’random’) behavior. VS-PMCA mice, but not other cohorts, also showed a significant decrease in action kernel learning rate (*α_k_*), consistent with reduced choice persistency (Supplemental Figure 10). We further validated our model fits with parameter recovery analysis to quantify the uncertainty in our parameter estimation: we simulated data using fitted parameters, refit the model to these simulated data, and then bootstrapped the pre- vs post- parameter comparison (Supplemental Figures 11). This procedure reproduced and validated the significant decrease in VS-PMCA *β_q_*.

In contrast to the selective change in inverse temperature in VS-PMCA mice, both VS-PMCA and VS-ctrl mice showed increases in learning rate (*α_q_*) (Supplemental Figure 10). To further verify that changes in inverse temperature, and not learning rate or other parameters were responsible for the decrease in choice performance win-stay behavior in VS-PMCA mice, we simulated changes in choice performance, win-stay, and lose-shift rates across the model parameter space. We empirically verified that within the parameter regime that fits animals’ behavior, changes in inverse temperature best explain the choice performance, win-stay, and lose-switch patterns that align with what we see in VS-PMCA mice’s behavior, rather than changes in learning rate or action kernel learning rate (Supplemental Figure 13). These effects were replicated in a version of our Q-learning model with separate learning rates for rewarded and non-rewarded outcomes. Both VS-PMCA and VS-ctrl mice showed increases in learning rates in that model, but only VS-PMCA mice showed a significant decrease in inverse temperature (Supplemental Figure 14). Thus, the increase in inverse temperature for VS-PMCA was not simply due to inadequate capture by a single learning rate term.

In sum, behavioral changes in choice performance and win-stay behavior induced by VS ACD attenuation are best captured by increased decision noisiness in a Q-learning model. We note that choice stochasticity inferred from models itself could be due to noisier learning or directly due to noisier choices (Nassar et al., 2010). To gain insight into how VS astrocytes support choice performance and win-stay stability, we next assessed what signals VS astrocytes carry *in vivo* while mice are engaging in the task.

#### Ventral striatum ACD is correlated with RPEs

We first aimed to assess how VS ACD relate to model inferred RPEs, because win-stay strategy is strongly related to post-reward positive RPEs (i.e., agents are more likely to repeat previous actions after unexpected reward outcomes). To this end, we trained naïve mice (n = 4) on a head-fixed version of the probabilistic decision-making task with the same reward statistics as the version used in Figures 1–3, except here mice were presented two lick spouts (left and right), and they indicated their choices by licking the left spout (left choice) or the right spout (right choice; Figure 4A).

**Figure 4.**
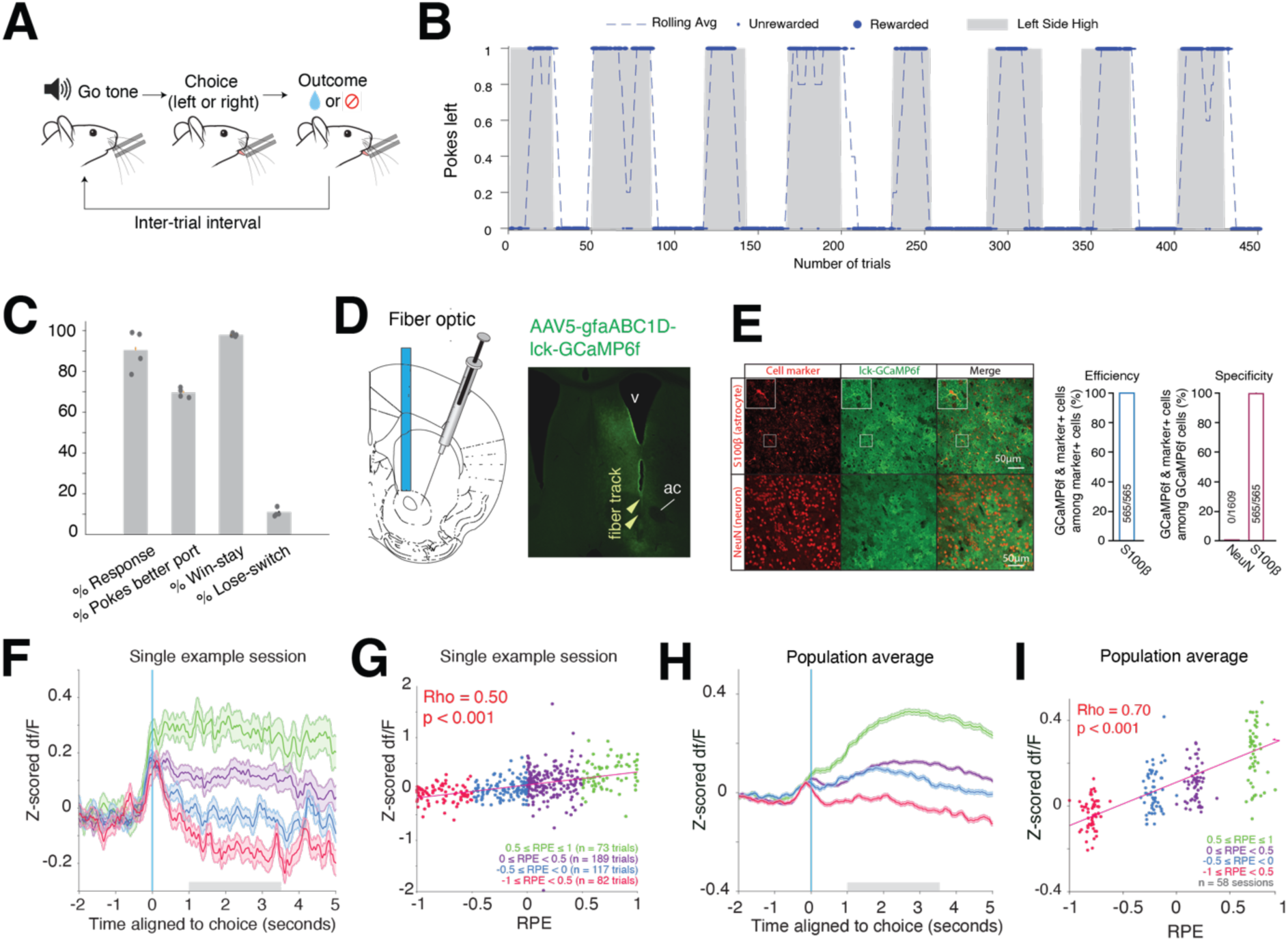
Astrocyte calcium dynamics in VS track reward prediction errors. A. Trial diagram for the head-fixed probabilistic decision-making task. On each trial, a go-tone indicated that the animal could make a choice by licking a left or right lick spout; this choice window persisted for 1.5 second. Reward probabilities were either 80% or 20%. Once the animal made a choice lick, the outcome was immediately delivered (reward or no reward). B. Behavioral data from one example session (conventions as in Figure 1B). Dots on the top and bottom indicate choices the animal made to the left and right and the dashed line shows a rolling average of left choices. The mouse made choices that closely follow the true underlying reward probabilities of left and right pokes. C. Behavioral metrics for mice in the head-fixed decision-making task. Dots indicate averages for each animal (n=4) and bars indicate average across animals. D. Schematic of the surgical procedure. AAV5-gfaABC1D-lck-GCaMP6f was injected into the ventral striatum, and a fiber optic was implanted 200um above the injection site. E. IHC of Ick-GCaMP6f expression in VS. S100β was used as an astrocyte marker and NeuN was used as a neuron marker. Quantification of efficiency and specificity of PMCA expression in astrocytes is shown on the right. F. Ventral striatum astrocyte calcium responses for one example sessions, split out by model inferred RPE. Individual traces show baseline subtracted fluorescence, averaged over trials with different RPEs (color key in G). Baseline was calculated per trial as the mean fluorescence in the 4-second window before the go cue started. VS astrocyte calcium activity scales with RPE, for example displaying greatest response for most positive RPE and weakest response for the most negative RPE. Error bars are S.E.M. over trials. Gray bar indicates time window used to calculate trial averages for G. G. Correlation between model-estimated RPE and post-decision astrocyte calcium activity for the example session in F. Each dot represents RPE and fluorescence from one trial. Y-axis is the mean value of baseline-subtracted fluorescence taken from a window 1-3.5 seconds after the outcome. There was a significant positive correlation between RPE and astrocyte calcium activity. Colors denote RPE levels (inset). Spearman’s correlation. H. Ventral striatum population astrocyte calcium responses (from all sessions and all animals), split out by model-inferred RPE. For shorter ITIs and strong RPE responses, calcium transients could remain elevated into the baseline of following trials. Error bars are S.E.M. over sessions. Gray bar indicates time window used to calculate trial averages for I. I. Correlation between model-estimated RPE and population post-decision astrocyte calcium activity. Each RPE bin shows within-session average fluorescence for trial outcomes with model-inferred RPE values within the specified bin. Conventions follow G.

Similarly to the free-moving task (Figure 1), trials started with an explicit go-cue consisting of an auditory tone and signal LED light, and reward contingencies were 80% and 20%. Reward contingencies switched between left and right spouts after 20-35 trials and inter-trial intervals were 5 to 9 seconds long. Outcomes were delivered immediately after a choice was made (Grossman et al., 2022).

An example session is shown in Figure 4B. In this session, the mouse’s choices closely aligned to the true underlying reward probability. Overall, mice exhibited high response rates and averaged correct choice rates of 69.5% +/- 0.8% S.E.M (22,249 trials included across 58 sessions, Figure 4C) on par with their performance in the free moving task (pre-injection correct choice rate across all groups = 67.5%). Their choice performance and switching behavior aligned to block switches is shown in Supplemental Figure 15A. Following block switches, mice increased their switching behavior, and their performance quickly recovered as they corrected their choices to the newly better port.

To confirm that mice’s choices reflected recent reward and choice history, we fit a generalized linear model (GLM) to their trial-by-trial decisions (Supplemental Figure 15B). The GLM captured key behavioral patterns observed in the task: mice tended to repeat previous choices, and this tendency was strengthened when the prior choice had been rewarded. These effects decayed over the last three trials. This model achieved high accuracy, correctly predicting the animal’s choice on 94% of trials, consistent with the hypothesis that recent reward–choice history accounted for a large amount of variance in behavior.

Having confirmed that mice can accurately perform the head-fixed version of our task, we next investigated if in-vivo VS ACD exhibits key signals predicted by our model and biology, and if these signals correlate with win-switch choices on a trial-by-trial basis. We injected AAV5-gfaABC1D-Ick-GCaMP6f into the VS to express the membrane-bound calcium indicator GCaMP6f in astrocytes and implanted an optic fiber to image bulk ACD over the injection site (Figures 4D, 4E). All 4 mice included in the study were successfully transduced (Figure 4E) and thereafter learned the task (Methods).

We first assessed the relationship of RPEs and ACD. ACD responses to trial outcomes aligned to Q-learning model inferred RPEs for an example session are shown in Figure 4F. VS astrocytes showed outcome responses that significantly correlated with the RPE level associated with the outcome (Figure 4G). ACD was highest when the RPE was highest (strongly positive RPEs in green), followed by relatively more expected rewards (positive RPEs in purple), expected non-rewards (negative RPEs in blue), and unexpected non-rewards (strongly negative RPEs in red). ACD correlation with RPE was strong and significant at the population level as well, across all mice and all sessions (Figures 4H, 4I).

These RPEs were reflected in ACD for many seconds (Supplementary Figure 16), for a relatively longer time than what is typically observed in single neurons’ firing activity (H. Kim et al., 2009; Oyama et al., 2010) or neurons’ calcium dynamics in response to RPEs (Zachry et al., 2024). Thus, VS astrocyte calcium levels encode outcome RPE signals that persist for many seconds, sometimes into the start of next trial.

The data thus far show that VS ACD are correlated with RPEs, a crucial value updating signal in RL, and that attenuating ACD disrupts decision-making performance by reducing win-stay decisions. Together, these data suggest that VS astrocytes contribute to the utilization of RPE signals in VS-guided post-reward decision-making. We next sought to draw on the biological circuit to generate hypotheses of potential mechanisms that could underlie this behavioral effect.

### Circuit modeling of neuron-astrocyte computations

To date, there are few (but emerging; see (Gerasimov et al., 2021; Meron Asher & Goshen, 2025; Serra et al., 2025)) techniques for manipulating astrocytes while simultaneously assessing large scale neural dynamics. And manipulation of ACD on timescales relevant for trial-by-trial RL remains a challenge. Therefore, to go further in assessing the algorithms and rules through which astrocytes could contribute to RL and generate testable hypotheses regarding their function, we constructed a computational model of VS with two key features: (1) it incorporates several mechanisms through which ACD could impact medium spiny neuron (MSN) function constrained by our (Figures 1–4; Supplemental Figures 16-17) and others’ previous data (Benoit et al., 2025; Bushong et al., 2002; Chai et al., 2017; Charles et al., 1991; Fujii et al., 2017; Mahon et al., 2006; Ogata & Kosaka, 2002; Stobart et al., 2018), and (2) it is capable of performing the bandit task, allowing us to probe distinct theories of astrocyte-neuron function and their effect on behavior (e.g., by comparing models’ behavior and the behavior of PMCA mice). In the model, we can then disrupt astrocytes in a relatively specific manner and observe the details of resulting task-behavior, resulting in precise hypotheses of how astrocyte-neuron interactions may support RL.

We constructed a minimal model of the cortico-basal ganglia circuit thought to underlie RL behavior, incorporating important aspects of RL-related basal ganglia architecture (Figure 5A; (Glimcher, 2011; Roesch et al., 2009; Samejima et al., 2005; Schultz, 1998; Sutton & Barto, 2018)). We incorporated astrocytes into this model with biologically-plausible functions and then assessed the role of different model astrocyte ablations on decision-making behavior in our task.

**Figure 5.**
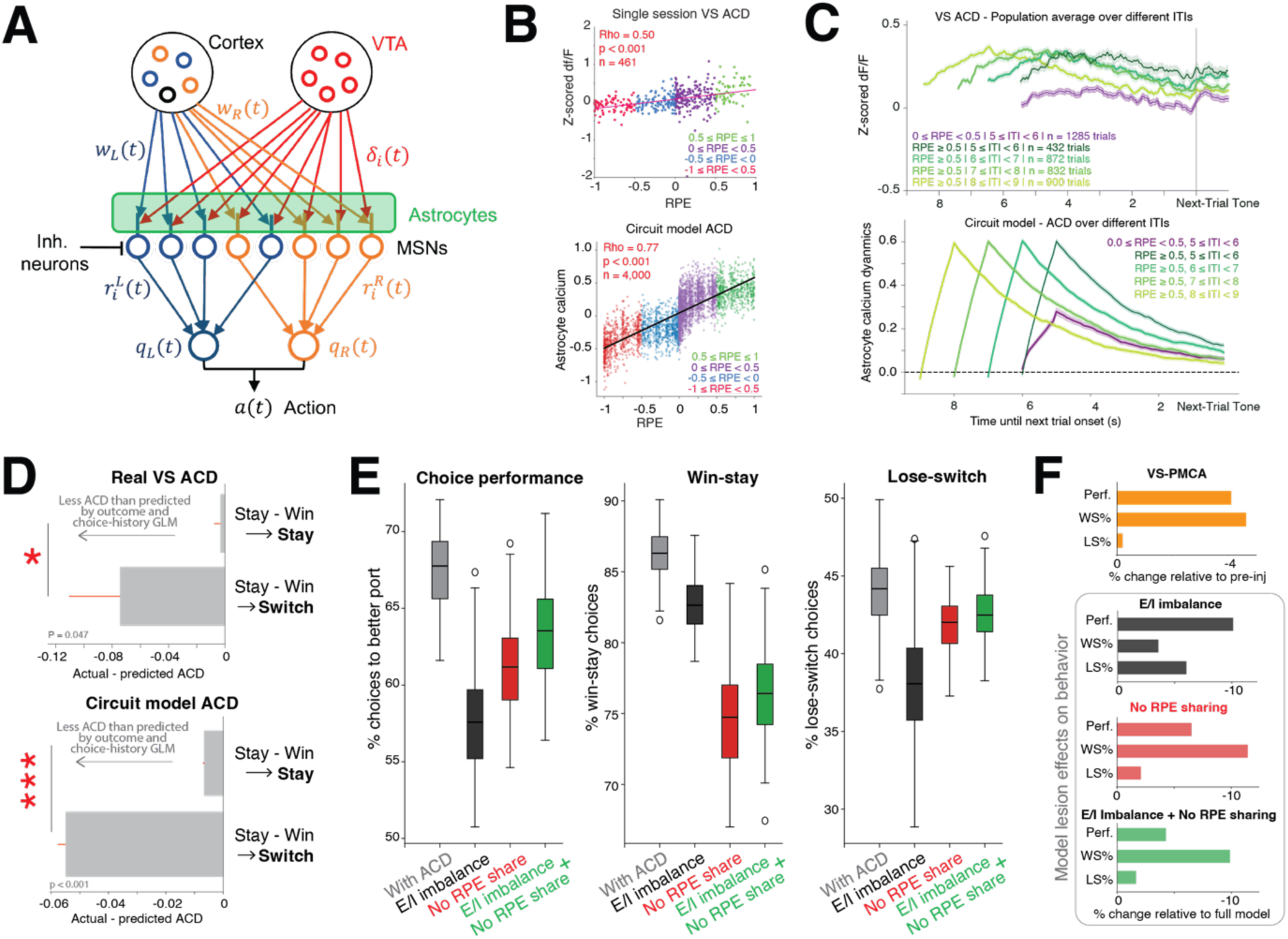
A circuit model recapitulates VS ACD trial-by-trial relationship to win-switch behavior. Disruption of astrocytic RPE-sharing *in silico* impairs decision performance and win-stay behavior. A. Schematic of the circuit model. B. Real VS ACD responses to outcomes with across different RPEs (top; same figure as 4G) and circuit model ACD responses to outcomes with different RPEs (bottom). By design, astrocytes in the circuit model correlate strongly with RPE. C. Real VS ACD responses (top) aligned to the start of the next trial. Green traces show post-outcome responses for highly positive RPEs across different ITIs; purple trace shows post-outcome response for a moderately positive RPE for the shortest ITI. For shorter ITIs and strong RPE responses, calcium transients could remain elevated into the baseline of following trials. Bottom shows equivalent circuit model ACD for trials of varying ITIs. D. Top: Model residuals of GLM fitted to real VS post-outcome ACD (Supplemental Figure 21A; calcium activity 1-3.5 seconds after outcome delivery) for stay-win-stay versus stay-win-switch trials. Trials where mice switched on the next trial had significantly lower ACD than predicted by choice and outcome history, compared to trials where mice stayed on the next trial. This effect was also significant at shorter time windows (p = 0.015 when GLM fit on calcium activity 1-2 seconds after outcome). Bottom: model residuals of GLM fitted to circuit model ACD (Supplemental Figure 21B, calcium activity 1-3.5 seconds after outcome). In alignment with real VS ACD, circuit model ACD was lower than expected for outcomes where the agent switched the next trial compared to trials where the agent stayed. N real ACD = 11,212 stay-win-stay trials, 177 stay-win-switch trials; N model = over 93 model fits (Supplemental Figure 20). E. Behavioral effects under different *in silico* ablation models of the circuit model. From left to right: ratio of poke to the better port, win-stay ratio, and lose-shift ratio under the full model (with ACD), excitatory-inhibitory drive imbalance, no RPE sharing, and both E/I balance + no RPE sharing. The error bars represent the S.E.M. over 10,000 simulated sessions. P < 10^-5^ for all pairwise comparisons. F. Summary of mean behavioral changes relative to baseline for real behavioral changes in VS-PMCA mice (top; relative to pre-injection behavior) and circuit model performance in various *in silico* ablation conditions (bottom, relative to ‘With ACD’ full model conditions). Perf. = choice performance, WS% = win-stay %, LS% = lose-switch percent. Changes in behavior in the no RPE share model ablation condition best matched the changes seen induced by VS PMCA.

Striatal circuit dynamics were modeled in the probabilistic decision-making task. Each trial time course has a decision period, a feedback period, and an ITI period (Methods). This formulation takes into account that choice and action do not happen instantaneously. Our model features two populations of MSNs, each representing the values of left or right actions in the task. On each trial *t*, the activity of the MSNs encoded the values of the left and right actions (*q_L_(t)* and *q_R_(t)*, respectively (Samejima et al., 2005)). These values are propagated to downstream circuits to generate the action a(t) (Sutton & Barto, 2018) (Supplemental Figure 19). We fit the model to match macroscopic parameters of choice performance, win-stay rate, and lose-shift rate that mice exhibited (Methods, Supplemental Figure 20).

MSNs receive several forms of input. Cortex sends excitatory input that varies across decision, feedback, and ITI periods; the strength of excitatory drive onto left- versus right-value MSNs is determined by projection weights *w_L_* and *w_R_*. Local interneurons provide inhibitory input that varies across task and ITI periods, without left/right selectivity (Banaie Boroujeni et al., 2020; Owen et al., 2018). Finally, VTA provides a noisy dopaminergic RPE signal during the feedback period, which is used to update *w_L_* and *w_R_.* Although dopamine has historically thought of as providing a bulk volume signal, more recent work has shown that dopamine transmission is more spatiotemporally specific and more variable across individual synapses than previously thought (Liu et al., 2021; Pereira et al., 2016; Yee et al., 2025).

Striatal astrocytes are integrated into our model with the following features. First, astrocytes can perform spatiotemporal integration of noisy dopamine signals. This function is supported by literature and our experiments showing that astrocytes respond to dopamine with elevated ACD across many seconds (Corkrum et al., 2020; Lazaridis et al., 2024) (Figure 1F, Supplemental Figure 16) and could share signals via extensive gap junction coupling (Cooper et al., 2026; Dermietzel et al., 1991). This could allow astrocytes to effectively mean-filter and add shared noise to RPEs across synapses, producing cleaner inputs for learning. This is conceptually similar to weight-sharing techniques in deep learning (Fukushima, 1980; Kipf & Welling, 2017; Krizhevsky et al., 2017; LeCun et al., 1989, p. 19) which are essential for training convolutional and graph neural networks. In our circuit model, ACD correlates with RPE (Figure 5B) and has prolonged ACD that can extend into the next trial (Figure 5C) – features which by design align with patterns seen in real VS ACD in our data.

Second, astrocytes can modulate the strength of excitatory and inhibitory drive onto MSNs. This function is supported by extensive literature indicating astrocytes shape synaptic efficacy and gate synaptic plasticity (Araque et al., 1999b; Corkrum et al., 2020; Haydon & Carmignoto, 2006; Henneberger et al., 2010; J. Kang et al., 1998). We also performed in-vitro slice electrophysiology experiments and found that PMCA-induced ACD attenuation in VS led to increased excitatory drive across both D1 and D2 MSNs, leading to a shift in excitatory-inhibitory balance (Supplemental Figures 17-18).

With this model in hand, we aimed to answer two key questions. One, does the model align with real data in describing how VS ACD relates to win-stay behavior on a trial-by-trial basis? Two, can we use this model to generate mechanistic hypotheses about how astrocyte function supports choice performance and win-stay behavior?

To address question one: we hypothesized that endogenous variations in RPE-related astrocyte Ca^2+^ signals could be related to variations in win-stay versus win-switch choice behavior. When ACD was broadly attenuated with PMCA, win-switch behavior increased (Figure 2A). We predicted that on a trial-by-trial basis, higher ACD after outcomes should predict more win-stay decisions, and lower ACD after outcomes should predict more win-switch decisions. In order to pinpoint the relationship between endogenous ACD fluctuations on future choices while controlling for choice and reward history, we regressed out the effect of past and current choices and outcomes on both real VS and circuit model ACD for each trial. Weights for the regression models (GLMs fit to ACD with weighted linear combinations of past and current choices and outcomes) are shown in Supplemental Figure 21. We further restricted our analysis to a subset of trials where the choice and outcome favored the animal making a stay decision on the next trial (stay-win trials). We hypothesized that variations in ACD outcome signaling may occasionally produce abnormally low signals and be linked to stay-win-switch decisions.

Indeed, we found that ACD preceding the more common stay-win-*stay* choices were aligned with values predicted by the GLM (Figure 5D). However, ACD preceding stay-win-*switch* choices were lower than predicted by the GLM, and significantly lower than stay-win-stay model residuals. This effect was significant in the analysis applied to both real VS ACD and animal behavior, and to the circuit model ACD and model behavior. Thus, when VS ACD was low, agents were more likely to make win-switch choices – even after controlling for previous choice and outcome history.

Finally, we turned to question two: can we use this model to generate mechanistic hypotheses about how astrocyte function supports choice performance and win-stay behavior? Our model allowed us to selectively disrupt either MSN excitatory-inhibitory balance or astrocyte RPE-sharing to assess the behavioral outputs of these *in silico* lesions.

First, we tested the effect of changing MSN activity. Assuming MSN firing follows a Poisson like process (N. Kim et al., 2019; Mahon et al., 2006), an increase in excitatory drive ought to lead to increased noise. Consistently, when we increased MSN baseline activity in the model, the choice performance and lose-switch percentage decreased, and there was a mild decrease in win-stay rate (Figure 5E-F, dark gray bars).

Second, we tested the effect of changing astrocyte responses to RPEs. When we removed the RPE-sharing from the model, both the choice performance and win–stay ratio showed a large decrease, while the lose–shift ratio remained nearly unchanged (Figure 5E-F, red bars). This pattern is consistent with experimental observations of VS-PMCA manipulation in our task (Figure 5F, orange bars; Figures 1–2). The same trends were also observed when both baseline adjustment and signal sharing were removed, which is likely what happens under PMCA injection (Figure 5E-F, green bars).

This work suggests how input sharing of dopaminergic learning signals across MSNs could be an additional novel mechanism through which astrocytes regulate decision variability, alongside other potential influences such as modulation of E/I balance. Crucially, using biological constraints, our model generated clear hypotheses that can be readily dissociable by future experiments as new techniques allow ACD excitation and suppression with spatial restriction and fast temporal precision. By formalizing astrocyte-neuron interactions in a model that can be used to study the details of animals’ and models’ task behavior, inspired by the biological data, we provide an early algorithmic description of how astrocytes could contribute to decision-making, highlighting the value of computational models for generating hypotheses that extend beyond current experimental reach.

In summary, we found that on average, VS ACD activity correlated with model-inferred outcome RPE during probabilistic decision-making. Even when accounting for the history of choices and outcomes, we found that lower-than-expected VS ACD levels were more likely to be followed by switch decisions on a trial-by-trial basis. Across our experiments, we find that lower ACD in VS is associated with an increase in post-rewarded outcome switching and increased decision noisiness or choice stochasticity. Finally, using a biologically constrained circuit model that captures key trial-by-trial relationships between ACD and win-switch behavior, we generate novel testable hypotheses about how astrocytes can mechanistically support value-guided decision-making by smoothing and sharing dopaminergic teaching signals across neural populations.

## Discussion

Astrocyte calcium dynamics are known to influence synaptic function and plasticity, shape neuronal activity, and affect behavior, but their specific role in decision-making algorithms to date has been unclear. Here, we provide evidence that astrocytes in ventral striatum play an important role in regulating decision noisiness. Attenuation of ACD in ventral striatum led to decreased choice performance, driven by decreased win-stay behavior and increased decision noisiness. We generated a novel model with neuronal astrocyte interactions that suggests a set of testable hypotheses for the circuit-level contribution of ACD, including input sharing – a prominent idea in deep learning – and regulation of MSN excitatory-inhibitory balance. Finally, we found ACD reflected reward prediction errors in the VS, and ACD trial-by-trial variability predicted future choices.

### Astrocyte function in striatum

We focused on the striatum because it is a key region for maintaining and updating value estimates for guiding behavior in neuro-RL frameworks (Averbeck & O’Doherty, 2022; Cox & Witten, 2019; Frank, 2025; Glimcher, 2011; Maia & Frank, 2011; Schultz, 1998). Since the respective role of different striatal regions in RL remains a subject of investigation, we attenuated astrocyte activity in different parts of the striatum and characterized the resulting changes on RL-dependent behavior. VS astrocyte calcium attenuation disrupted RL behavior, consistent with theories proposing a functional gradient along the ventral-dorsal axis in which ventral-medial regions index value and value-updating processes and dorsal-lateral regions support control of habits, skills, and motor functions (Alexander et al., 1986; Balleine & O’Doherty, 2010; Burton et al., 2015; Haber & Knutson, 2010; Ito & Doya, 2011; H. F. Kim & Hikosaka, 2013; Mohebi et al., 2024)

A growing body of evidence supports the notion that astrocytic function follows this dorsal-ventral functional gradient. In the dorsal striatum, astrocyte function has been linked to behavioral perseveration, behavioral flexibility (Boender et al., 2021; S. Kang et al., 2020; Montalban et al., 2025; Yu et al., 2018), and engagement (Lazaridis et al., 2024). Ventral striatal ACD seems to play a role in preference learning and drug-reward self-administration (Bull et al., 2014; Mongrédien et al., 2025; Serra et al., 2025). Together, these findings support the idea that astrocytes are functionally embedded within striatal computations in a regionally organized manner, operating alongside neuronal circuits while contributing on distinct spatial and temporal scales (Allen, 2014; Araque et al., 1999a; Khakh, 2025; Ma et al., 2016; Murphy-Royal et al., 2023).

Although astrocyte functions broadly align with the dorsal–ventral functional axis, they cannot be cleanly distinguished at narrow spatial scales (Charles et al., 1991; Cooper et al., 2025; Dermietzel et al., 1991; Fujii et al., 2017; Murphy-Royal et al., 2023; Oberheim et al., 2009; Serra et al., 2022). For instance, we cannot distinguish between the effects of PMCA-related attenuation in nucleus accumbens core versus shell, both of which are in VS. Viral spread in nearly all VS injections encompassed both subregions. While these subregions have distinct anatomical projections and are thought to contribute differentially to behavior (Castro & Bruchas, 2019; Di Chiara, 2002; Zahm, 1999), recent studies show astrocytes may have complex interactions with neural circuits across space that are not cleanly delimited by neuronally-defined subregion boundaries. (Cooper et al., 2026; Serra et al., 2022).

In our computational model, this broad spatial influence is a feature—not a bug—that underlies the input-sharing hypotheses we propose. Input-sharing maybe critical to consider when modelling how the brain implements many forms of RL (Dabney et al., 2020; Lowet et al., 2025) in part because the spatial scale of dopaminergic influence on the striatum maybe more sharp or precise than previously assumed (Yee et al., 2025).

### RPE sharing and functional stabilization

We found that VS ACD correlated with a model inferred RPE (Figure 4G, 4I) – a key teaching signal in RL (Glimcher, 2011; Schultz, 1998). This post-outcome ACD signal predicted future choices, with larger ACD outcome responses specifically predicting more win-stay behavior and weaker ACD outcome responses predicting more win-switch behavior – matching the results of PMCA ACD attenuation. Because VS ACD is highly sensitive to dopamine release (Corkrum et al., 2020), future studies must uncover the details of how dopamine modulates astrocyte-neuron interactions (Allen, 2014; Araque et al., 1999a; De Pittà et al., 2011; Favetta & Bubacco, 2025; Lefton et al., 2025) as well as create further computational models to formalize these interactions in the context of learning theory. We note that our theories are agnostic to whether astrocytes play an active role in computing RPEs or value in RL.

The BOLD signal in ventral striatum has been shown to encode reward prediction errors (O’Doherty et al 2003). Notably, astrocytes are well-known to mediate neurovascular coupling (Haydon & Carmignoto, 2006; Petzold & Murthy, 2011), and can drive some components of the fMRI BOLD responses even with limited neuronal activation (Institoris et al., 2022; Takata et al., 2018). The astrocytic RPE response we find likely underlies a component of BOLD RPE responses, although future studies will be needed to fully disambiguate neural versus astrocytic contributions.

Our circuit model suggests that even simple linear input sharing of RPEs across neighboring MSNs may be enough to impact RL (Figure 5E). This type of function is well suited to astrocytic morphologic and physical traits (i.e., responses to dopamine (Corkrum et al., 2020), spatial spread across domains (Bushong et al., 2002; Fujii et al., 2017; Halassa et al., 2007; Ogata & Kosaka, 2002), gap junction coupling (Dermietzel et al., 1991), and integration of inputs over slow calcium timescales (Benoit et al., 2025; Charles et al., 1991; Fujii et al., 2017; Panatier et al., 2011; Scemes & Giuame, 2006; Stobart et al., 2018). Astrocytes may help control the variability of underlying neurons within functional domains by averaging and sharing information across units representing the same cue or value. This proposal is conceptually like weight-sharing techniques in deep learning (LeCun et al., 1989), which are essential for training convolutional and graph neural networks (Kipf & Welling, 2017; Krizhevsky et al., 2012). A recent but growing body of work is examining how astrocytes may be involved in normalization of local neural circuit activity (Gong et al., 2024; Kozachkov et al., 2023; Miguel-Quesada et al., 2023).

This hypothesis could be extended to test how astrocytes stabilize perceptual and cognitive representations, beyond the striatum or RL (Doron et al., 2022; Miguel-Quesada et al., 2023). There is some evidence to suggest astrocytes could perform such normalization functions as in functional columns in sensory cortex (Philips et al., 2017; Schipke et al., 2008). Consistent with this hypothesis, indiscriminate activation of astrocytes can enhance behavior in ways that indiscriminate activation of neurons does not (Adamsky et al., 2018). Recent work suggests that within the VS, astrocytes form functional ensembles associated with specific cue-value associations (Serra et al., 2025). Emerging technologies will help investigators evaluate these processes in vivo.

Future studies are required to understand the relationship between this type of linear input-sharing hypotheses generated by our RL study and recently proposed ideas of how astrocytes can actively shape the types of computations performed by neural circuits in state-dependent, non-linear ways (Murphy-Royal et al., 2023). Astrocyte integration of inputs and active shaping of circuit computations need not be mutually exclusive (Mu et al., 2019). What is shared and what differs between these ideas – i.e., molecular mechanisms and computational, spatiotemporal scales – requires careful assessment.

### Consideration of other astrocyte functions

Our circuit model generates testable hypotheses about how astrocytes may help stabilize value-guided decision-making by reducing variability in population learning signals – likely phasic dopamine RPE signals – across neural populations. Astrocytes may also modulate dopaminergic tone in the striatum. *In-vivo* striatal astrocyte calcium responses modulate and reduce subsequent dopamine release (Lazaridis et al., 2024), and astrocytes are known to clear extracellular dopamine through DAT (Segura-Aguilar et al., 2022). Blockade of ACD could potentially attenuate astrocyte dopamine clearing functions and lead to increased tonic dopamine tone. Prior modeling and some experimental work point indeed to the notion that hyperdopaminergic tone could decrease inverse temperature, not learning rate (Beeler et al., 2010; Collins & Frank, 2014) (Figure 3H). Directly assessing whether and how tonic dopamine tone changes due to astrocyte activation or attenuation across different behavioral states is an important next step for understanding astrocytic function in the context of striatal circuitry. Importantly, a change in dopamine tone over long timescales is unlikely to account for all our results, because we observed variations in ACD linked to win-stay behavior unfolding on relatively short, trial-by-trial timescales.

In addition to dopamine (Corkrum et al., 2020; Lazaridis et al., 2024), striatal astrocytes also respond to influence the processing of nearly all signaling molecules present in striatum and elsewhere, such as glutamate (D’Ascenzo et al., 2007, p. 201; Martín et al., 2015; Serra et al., 2022), GABA (S. Kang et al., 2020; Nagai et al., 2021; Roberts et al., 2020; Yu et al., 2018), acetylcholine (Papouin et al., 2017; Stedehouder et al., 2024; Takata et al., 2011), opioids (Corkrum et al., 2019; Murlanova et al., 2023), endocannabinoids (Adermark et al., 2024; Martín et al., 2015; Navarrete & Araque, 2010), serotonin (Miyazaki & Asanuma, 2016), and of even energy metabolites (Molnár et al., 2011). As we gain deeper insight into how different modulators together cooperate to shape learning, their impact on astrocytes and astro-neuro circuit function needs further study.

Astrocytes could also participate in the control of excitatory and inhibitory synaptic drive. We found that PMCA attenuation of VS ACD altered shifted E/I balance towards increased excitation in both D1 and D2 pathways, likely through a presynaptic mechanism (Supplemental Figures 17-18). This is consistent with literature indicating that astrocytes can directly inhibit synaptic excitatory drive via an adenosinergic, A1R-dependent mechanism that suppresses presynaptic glutamate release (A. B. Chen et al., 2025; Corkrum et al., 2020; Lefton et al., 2025). However, because of the unique properties of D1 and D2 MSNs, it is still possible that the net impacts of ACD are distinct among the neural subtypes. These distinctions, as well as other potential mechanisms (e.g., changes in local versus exogenous presynaptic inputs, presynaptic inhibition) should be disambiguated in future work with experimental and modeling work that that will account for biophysical, electrophysiological and anatomical properties of these projection neurons (i.e. (Collins & Frank, 2014)).

We used a viral approach that broadly and indiscriminately attenuates astrocyte calcium activity via constitutive calcium pump (Yu et al., 2018) as a first-line strategy to investigate how astrocyte calcium transients, the main functional signal of these cells, impact value-guided behavior. It is important to note that while PMCA significantly attenuates calcium signaling, it does not ‘turn off’ astrocytes – PMCA-expressing astrocytes undergo a variety of gene and protein expression changes (Yu et al., 2018). Thus, it is possible that some of the behavioral effects we see are due to gene expression changes above and beyond downstream effects of calcium attenuation (Yu et al., 2018). Additionally, it is possible that the extrusion of astrocytic calcium may alter local neural activity by altering neural calcium signaling (Berridge, 1998), although this is unlikely, as astrocyte cytosolic calcium levels at rest (∼100 nM (Zheng et al., 2015)) are orders of magnitude below normal extracellular calcium levels at rest (∼1.2 mM (Jones & Smith, 2016)). All this said, it remains that specific targeting of astrocytic calcium signaling in specific brain regions without inducing glial or neural depletion (Meron Asher & Goshen, 2025; Yu et al., 2018) makes for a powerful and useful method to study astrocyte contributions to behavior in a circuit-specific manner.

### Astrocyte computations in RL

Our relatively simple task allowed us to start to assess, in a high-throughput manner, the relationship between astrocyte activity across striatum and the core RL computation of how trial-by-trial decision-making is guided by previous reward and choice history. VS astrocyte attenuation led to decreased inverse temperature as captured in a Q-learning model. This normative learning framework allowed us to capture and describe consistent changes in VS-PMCA animals’ behavior, but our utilization of these theories does not imply that the model is instantiated in a 1:1 mapping in the brain.

While inverse temperature in this task and parameter regime led to changes in win-stay rate, in other regimes the same parameter may affect other behavioral outputs, including lose-shift rate. While changes in inverse temperature manifest as decision noise, future studies with task variations beyond those explicitly tested here are required to dissociate this from other RL-related changes, as in directed vs undirected exploration.

These findings pave the way for future studies with more complex task designs that allow for further exploration of cognitive and learning processes as uncertainty and behavioral strategy vary (Akam et al., 2015; Costa et al., 2016; Doll et al., 2016; Grossman et al., 2022; Niv et al., 2015; Wilson et al., 2014). Some groups are already linking potential astrocyte function to RL in theoretical frameworks (Gong et al 2024, Chen and Yang, 2025). We believe it will be particularly important to explore astrocyte computations in non-human primates, where more complex RL schedules can be used (Costa et al., 2016; Muller et al., 2024; Oemisch et al., 2019; H. Tang et al., 2021) and emerging technologies make functional, cell-type specific manipulations possible (Heffernan et al., 2022; Tremblay et al., 2020).

### Summary

Our findings suggest that VS astrocytes influence decision variability. Their ACD covaries with RPE and predicts future win-stay behavior. Computational modelling suggested that their impact on behavior could occur through input sharing across neuronal populations and via modulation excitability over broader spatiotemporal scales than neurons. Our results generate novel theories for how astrocytes contribute to RL and algorithm level computation. These implications extend beyond biology. Understanding how astrocytes regulate noise, excitability, and information sharing provides a conceptual bridge between molecular mechanisms, circuit-level models of decision-making, and neuromorphic computing (the effort to design hardware to run AI based on the architectures and circuit motifs of the brain). Embedding astrocytic principles into artificial systems may yield architectures that are both more biologically realistic and computationally efficient — for example, by reducing training duration through hardware that enacts context-dependent input sharing, these principles may enable biologically plausible neural networks to learn faster while retaining the advantages of local learning rules over global gradient descent. Our study provides one entry point into this emerging frontier.

## Materials and methods

### Subjects

76 C57BL/6 mice (37 male, 39 female; Table 1) were obtained from Jackson Laboratory and from in-house breeding. Animals were housed individually and maintained on standard 12/12 light dark cycles. All procedures conformed to the Guide for the Care and Use of Laboratory Animals and were approved by the Institutional Animal Care and Use Committee at Washington University.

### Behavioral training

Mice were trained to perform tasks on a FED3 device (Matikainen-Ankney et al., 2021). These are small, open-source operant devices running off Arduino code with a left and right nose-poke port, pellet delivery system, a strip of Neopixel lights on the front of the device, and a piezo speaker. A FED3 was placed against the back wall of the home cage of each mouse. Custom cut stainless steel cage dividers magnetically attached to the front of the FED3s prevented the mice from accessing the backs of the devices. During behavioral training, mice were food restricted and received all food (Dustless Precision Pellets® Rodent, Grain-Based 20mg, Product Number, F0163, BioServ) from the FED3. Mice were weighed before behavioral training to establish their baseline weight and weighed every week during training to ensure their weight did not drop below 80%. None of the mice used had weights that fluctuated below 80% of their baseline weight.

Mice were first acclimated to the FED3s and to receiving pellets from the FEDs during an initial 1-2 day magazine training period. During magazine training, pellets were automatically delivered into the hopper, or magazine, without any action needed from the mouse. Mice that successfully learned how to pick up pellets from the hopper progressed to the next training stage. The few mice that were unsuccessful in learning how to pick up pellets were removed from the study.

The next training stage was a time-unrestricted version of the operant non-stationary probabilistic decision-making task, also known as a bandit task. In this task, the left and right nose-poke ports were associated with either 80% or 20% pellet delivery, such that P(left) = 1 – P(right). If the mouse made a rewarded poke, the FED3 delivered a 200ms 4kHz tone and a pellet was delivered into the hopper. If the mouse made an unrewarded poke, a 300Hz error tone was played. After either a rewarded or unrewarded ‘choice’ poke, the FED3 entered a 5 second time out period during which the Neopixel lights turned off and pokes could not trigger any outcome, although they were still logged. After 20-30 delivered pellets, the reward contingencies between the two nose-poke ports switched, uncued to the mouse.

During this training stage, mice learned that nose-poking was necessary for pellet dispensal, and they started to learn the structure of the task. In this intermediate training stage, the task was available to the mice between the hours of 1am and 9pm. When the task was active, the strip of Neopixel lights on the front of the FED3 turned blue, indicating that pokes could lead to reward- or no-reward outcomes. Outside of active task hours, the Neopixel lights turned off and pokes could not trigger any outcome, although they were still logged.

After 1-2 days of the relatively time-unrestricted bandit task training stage, mice started the final stage of the bandit task, during which the task was only available to the mice from 5pm to 9pm. Animals on average initiated around 200 trials in each 4-hour session (mean = 194.7, SD = 38.2). After 7-10 days from the start of this training state, the performance of each mouse over days was assessed. Mice that could not consistently achieve 60% correct choice rate were removed from the study because studying their learning or adaptation during block switches was not possible (e.g., low performance indicated that they did not learn or switch as the task contingencies changed).

Mice were further trained for 3-4 weeks on the bandit task (Figure 1) and then injected with control or PMCA virus into different regions of striatum. After virus injection, mice recovered in their home cage on ad lib normal chow with no FED3. After 2 weeks of recovery and virus expression time, mice were again food restricted, the FED3s programmed with the final time-restricted bandit task were placed back in their home cage, and behavioral data was collected for another 3-4 weeks. To capture block switch related learning changes rather than transient changes related to a post-surgery rest period, we let mice acclimate to the task again after surgery (see Behavioral data analysis section).

We ran an additional cohort (n = 7) of VS-PMCA injected mice on a version of the task where block switches occurred every 20-40 trials irrespective of rewards earned (reward non-contingent task), to test that these PMCA-induced behavioral effects were not specific to the reward contingency meta-structure of the first task (reward-contingent task). We found similar decreases in performance and win-stay behavior in this smaller cohort (Supplemental Figure 5). Data from both groups were combined and collectively referred to as VS-PMCA results.

To obtain more controls, we injected two cohorts in VS with a control virus (AAV5-GFAP104-mCherry or AAV5-gfaABC1D-mCherry), running one cohort on the reward-contingent version of the task (n = 13) and one cohort on the reward non-contingent version of the task (n = 5). We also injected a cohort of mice on the reward-contingent task with saline in the region of motor cortex overlying VS (n = 6). None of these cohorts exhibited significant shifts in correct choice rate, win-stay, or lose-switch behavior after injection compared to before. Data from these three cohorts were pooled and collectively referred to as ‘VS-control’ (Supplemental 5).

For the PR1 progressive ratio task, FED3s were programmed to deliver pellets in respond to increasing number of pokes to the left nose poke port. The right nose poke port was inactive but still logged pokes. The task was available to mice continuously over 5 days of testing. After each pellet was dispensed, the number of pokes required to dispense another pellet increased by one. This continued until the mouse did not make any pokes in a 30-minute window. At this point, the number of pokes the mouse had made for that last pellet was recorded as one breakpoint, and the threshold for dispensing the next pellet reset to one poke.

### Virus injection

Mice received intracranial injections of control (AAV5-GFAP104-mCherry, Addgene 58909-AAV5, or AAV5-gfaABC1D-tdTomato, Addgene 44332-AAV5) or PMCA (AAV5-pZac2.1-GfaABC1D-mCherry-hPMCA2w/b, Addgene 111568-AAV5), targeted at different regions of striatum. Surgeries and behavioral tests were interleaved between experimental cohorts. Anesthesia was induced with 3-4% isoflurane and maintained at 1-2%. Buprenorphine SR was injected subcutaneously at 1.0 mg/kg at the start of surgery for pain relief. After induction of anesthesia, mice’s heads were shaved, sterilized with alternating betadine and alcohol swabs, and an incision was made over the skull. Coordinates were marked bilaterally on the skull for ventral striatum (AP: +1.0-1.2, ML: +/- 1.2, DV: 4.5 relative to dura), dorsomedial striatum (AP: +0.7, ML: +/- 1.5, DV: 2.5 relative to dura), and dorsolateral striatum (AP: +0.7, ML: +/-2.5, DV: 3.0 relative to dura) relative to bregma. A 0.5mm burr hole was drilled over the marked coordinates (Fine Science Tools 19007-05, Foredom K.1070 Drill). Virus was loaded into a syringe (Hamilton Neuros 65457-02), slowly lowered into place, and delivered (VS: 0.25 µl/side at 0.05 µl/min; DLS and DMS, 0.45 µl/side at 0.1 µl/min). After virus delivery, the syringe was raised by 100-200 µm. An additional 5 minutes were allowed to pass before the syringe was slowly raised. After injections were made, the skin incision was sutured, and mice were returned to their home cage to recover.

Our study required bilateral expression in VS, DMS, or DLS. Animals without strong bilateral expression were excluded. Occasional virus expression in nearby regions adjacent to the injection needle track was noted; a common and difficult to avoid effect of intracranial virus injection. We report percentages of animals with off-target virus expression in nearby brain regions in Supplemental Figure 2A.

For fiber photometry, the same surgical procedure was followed. Mice received intracranial injections of AAV5-pZac2.1-gfaABC1D-Ick-GCaMP6f (Addgene 52924-AAV5) in VS (AP: +1.0-1.2, ML: +/- 1.2, DV: 4.5 relative to dura); a fiber optic (Doric; 200um ID, NA0.37, 5mm long) was implanted 200 µm above the virus injection site.

### Behavioral data analysis

All statistical tests of difference wherever p-values are reported were non-parametric and two-sided unless otherwise noted. Correlations were Pearson’s correlations unless otherwise noted. For pre-injection sessions, we took the last 20 sessions relative to the injection date. For post-injection sessions, we took the 20 sessions after 4 days of task re-acquisitions. Any sessions with fewer than 70 trials were excluded. For cohort analyses, data from pre-injection and post-injection session were averaged for each animal. Wilcoxon sign rank tests were then run on pre- and post-injection data for each cohort. For within-animal analyses, Wilcoxon rank sum tests were run for pre- and post-injection sessions for each animal.

For Bayesian inference of block switches, we followed methods developed and described in (Bartolo & Averbeck, 2020). In essence, the algorithm operates as follows. Around each true change point, where the reward probability shifts, we defined a window spanning half a block length before and after the change. Within this window, we evaluated the likelihood that each trial represented the change point, under the assumption that exactly one change point was present. Both choice and reward information were used to estimate how an ideal observer would classify a trial as occurring before or after the change. The trial with the highest likelihood was then identified as the inferred block switch point. If multiple trials within the window had nearly identical likelihoods (differing by less than 0.01), the corresponding block was excluded from further analysis. The same window and analysis were used to identify changepoints as the animal detected them, based off choice information alone.

The same algorithm was used to analyze block switches for both the reward contingent and reward non-contingent versions of the bandit task. Since the changepoint detection algorithm relies solely on the likelihood, not any priors, the algorithm takes the identical form in both cases.

A similar algorithm using just the choice behavior was used to infer when animals detected a change point for each block switch. We used this to infer the latency (in trials) of how long it took animals to infer block switches, relative to ideal observers (restricted to block switches where it was possible to infer both).

### Head Fixed Bandit Training - Habituation

After recovering from surgery mice were water restricted at 1ml per day. After 1 day of restriction mice began participating in 2-4 sessions of habituation handling daily. For the first 2-3 days, mice were placed in the palm of the experimenter’s hand and offered water via syringe. For 1-3 days after mice were held over the location where they would be head posted and offered water via syringe. Then for 1-2 days mice were placed on the wheel and held to the head post by the experimenter while being offered water via syringe. Finally, for 1-3 days the mice were fully head posted and were periodically rewarded with 4 ul of 15% sucrose and water via the lickers in the behavior rig. Rewards were given manually in 1-3 second intervals, alternating which side was rewarded every 5-10 rewards. Each habituation session lasted between 5-15 minutes or until the mice showed visible agitation, freezing, or refusal to receive reward. Steps were repeated as needed for mice who showed reluctance to participate.

### Head fixed Bandit Training - Task Training

After habituation training, mice began training for the probabilistic decision-making task. While most mice generally followed the listed progression, steps were repeated as needed for mice who showed reluctance to participate or tended to choose only one licker. At every stage, if the mouse made ∼150-200 consecutive choices on the same side, the task was paused and the position of the lickers adjusted to make the unchosen side closer than the other.

The task took place in a sound attenuated box, wherein mice were head fixed atop a stationary styrofoam wheel. Two lickers fixed approximately 4mm apart were placed directly in front of the mouse’s mouth such that each licker was within reach. All tasks described used a 0.3s go cue and 4 microliters of 15% sucrose and water for reward. Rewards were administered immediately after selection. Additionally, to signify the length of the response window and to provide visual feedback, a signal LED was placed in the mouse’s left periphery. This light turned on at the beginning of the response window and remained on until the window closed, or the mouse made a choice.

The first day of training was on a Pavlovian task. Mice were rewarded for licking from either licker within 10s after the tone. The task ran for 400-450 trials or until the mouse refused to participate. On day 2, mice began a task where only one licker was active at a time. Following the go tone the mice had 10s to lick the active licker to receive reward. Licking the inactive licker gave no feedback. The active licker was determined by alternating blocks 5-10 trails in length. Additionally, we varied the inter trial interval (ITI) between 2.5-5.5 seconds. On days 3-4 we used the same task structure but progressively tightened the window to respond (from 5s to 3s) while increasing the ITI (from 3.5-6.5s to 5-9s). On days 5 and 6 we activated both lickers. The mouse was still tasked with finding the licker that guaranteed reward, but a wrong choice would end the current trial. Both days used a 1.5s response window and 5-9s ITI. Day 5 we used a 15-25 trial block length but increased to 20-35 on day 6. However, only correct choices contributed towards the block length. Such that mice who made more errors had longer total block lengths. Day 7 and onwards used the full task. Probabilities were introduced following the same convention as used in the free moving task. Task parameters were held constant with a 1.5s response window, 5-9s ITI, and 20-35 trial block lengths. Additionally block length was no longer reward dependent

### Fiber photometry recordings

Fluorescent calcium signals were acquired with a FPS-S fiber photometry system from Doric Systems, through which we could excite and collect emission light from GCaMP from the same implanted optical fiber. Excitation wavelength was 488nm; we used a 405nm LED for isobestic excitation. Light was filtered and collected through a 4-channel fluorescent MiniCube (Doric) coupled via a low autofluorescence mono fiber optic patch cable (Doric).

Photometry signals were first preprocessed by fitting the isosbestic signal (IS) to the GCaMP transporter signal (TS). We subtracted and then divided the GCaMP signal by this fitted isobestic signal (fIS) (df/F = (TS -fIS) / fIS); this signal was then smoothed with a 200ms window. Since individual trials often started at different initial fluorescences, trial by trial analysis used baseline subtracted values. We defined a baseline for each trial as the mean fluorescence in the 4 secs before tone start. Individual trial fluorescence is reported as the difference between df/F and this baseline value. Additionally, due to the slow time course of astrocytic calcium, when analysis dictated a single measure, we defined the post outcome fluorescence as the mean fluorescence between 1 to 2 seconds after outcome.

### Head-fixed behavioral and photometry analysis

We included 58/89 sessions (24,609 trials; 22,249 trials responded) from 4 mice in our data. Days were excluded if mice did not reach either >60% participation or >60% correct choice rate. This criterion was applied to all analyses performed. RPE was estimated using a Q-learning RL model (see below section on RL modeling). Each trial was sorted into buckets based on its RPE: strong positive (0.5 <= RPE <= 1), low positive (0 <= RPE < 0.5), low negative (-0.5 <= RPE < 0), and strong negative (-1 <= RPE < 0.5). These buckets were used for all analyses that tested for RPE related effects.

To capture past trial effects on choice, we constructed a GLM to predict log odds of a left choice using a logistic link function for binomial data. For regressors we included past trial choice (Left: 1, Right:-1), outcome (Rewarded: 1, Unrewarded: -1), Choice x Outcome, and a spatial bias term (1). The model was fitted to 19,386 trials pooled from all 4 mice; trials were excluded if one or more of the regressors was missing a value. We included data from 1 to 3 trials prior.

We constructed another GLM to predict mean fluorescence 1-2s post outcome using a linear link function and Gaussian noise distribution. We used a similar construction as in the choice GLM; however, we defined choice in terms of whether it was the same or different than the current trial choice. We included regressors for current outcome (Rewarded: 1, Unrewarded: -1), Past (1-3 trials prior) Choice Same (same:1, else: 0) x PastOutcome, Past (1-3 trials prior) Choice Different (different: 1, else: 0) x PastOutcome, and side bias (1). We fitted the model to 19,386 trials pooled from all 4 mice; trials were excluded if one or more of the regressors was missing a value. We included data from 1 to 3 trials prior. We then looked at the residual fluorescence (actual - model predicted) in trials where mice made two consecutive choices on the same side and were rewarded and divided them based on whether the next choice was the same (“stay-win-stay”, n=11,212) or different (“stay-win-switch”, n=177).

### Perfusion

Animals were deeply anesthetized with a 100/10 mg/kg ketamine-xylazine cocktail injected intraperitoneally. Animals were then intracradially perfused, first with 15ml of 1X phosphate buffered solution (PBS), followed by 15ml of 4% paraformaldehyde (PFA). The brains were extracted and stored in 4% PFA overnight. They were then transferred to a 30% sucrose solution where they were left to fully saturate for at least 2 days for cryoprotection. Serial coronal sections were cut at 50 µm on a cryostat (Leica CM1860). Sections were stored in 1X PBS or 0.1% sodium azide in 1X PBS at 4deg C.

### Histology

Free floating sections were rinsed in 1X PBS three times for 5 minutes each. Sections were incubated in 10% normal goat serum (NGS) in 0.3% Triton-X in 1X PBS for 1 hour at room temperature, then incubated in primary antibodies (rabbit anti-mCherry, abcam ab167453; 1:800-1:1000; guinea pig anti-NeuN, Sigma Aldrich ABN90, 1:1000; guinea pig anti-S100B, Synaptic Systems 287-004, 1:1000) overnight at room temperature. Between primary and secondary incubation, sections were thrice rinsed in 1X PBS. Secondary antibodies (goat anti-rabbit AlexaFluor 555; goat anti-guinea pig AlexaFluor 488; goat anti-rabbit AlexaFluor 488) was applied at 1:300 for 4 hours at room temperature. Finally, sections were rinsed twice with PBS for 5 minutes, followed by a 5-minute incubation in 1:1500 DAPI before being mounted on glass slides and cover slipped with Prolong Diamond Anti-fade mounting medium. Mounted sections were imaged with a Leica DM6 B or a Nikon AXR confocal microscope.

All animals tested were imaged to ensure that virus injections were bilaterally expressed and localized to the target region. Animals with unilateral, weak, or overly extensive viral expression were excluded from the dataset.

### In vitro calcium imaging

#### Two-photon imaging

GCaMP6f recordings of calcium activity in the striatum astrocytes were performed in acute slices from C57BL/6J mice microinjected with AAV5-gfaABC1D::lck-GCaMP6f (Addgene 52924-AAV5) three to four weeks prior. Two-photon laser scanning microscopy (2-PLSM) recordings of GCaMP6f fluorescence over a field of view of 452 µm × 452 µm at 512 × 512 pixels resolution were captured using Bruker Ultima 2pPlus with Nikon 25X, 1.10NA objective, 1.6x optical zoom, laser power at 37.5 mW, recording rate at 1 Hz, recording depth at 30-40 μm. A 920 nm tunable laser was used to excite the fluorophores. Acute slices were incubated in a chamber with oxygenated artificial cerebrospinal fluid (aCSF) containing 1µM tetrodotoxin to block neuronal firing. After a stable baseline recording, stock norepinephrine (NE, norepinephrine bitartrate, Tocris 5169) or dopamine (DA, dopamine hydrochloride, Tocris 3548) solution is spiked into the aCSF to reach a working concentration of 20 µM for NE and 100 µM for DA and astrocytic calcium response is recorded.

#### Calcium imaging analysis

Astrocytic calcium responses in the striatum to NE and DA are performed across the field of view (FOV) for each recording for both the control and CalEx groups. For the control group, cell boundaries are delineated with the aid of static tdTomato fluorescence using Fiji for further cell-based analysis. Average fluorescence intensity across FOV or each cell is extracted from the recordings, and the traces are aligned so perfusion of NE and DA starts around 100 s into the recording. The traces are then processed using STARDUST (Wu et al., 2024) to detect active calcium signals using two standard deviations of the baseline as the threshold. The largest active signal within 100-300 s of the recording is taken as NE/DA response. For FOV analysis, in cases where no active signal is detected within the pre-determined temporal window, the largest fluorescence value between 100-300 s and its corresponding area under the curve are evaluated.

### Ex vivo slice electrophysiology

Whole-cell patch-clamp recordings were performed as previously described (Lucantonio et al., 2025). Briefly, mice were anesthetized with isoflurane and decapitated. Brains were rapidly extracted and placed in ice-cold, NMDG-based cutting solution (in mM: 92 NMDG, 20 HEPES, 25 glucose, 30 NaHCO₃, 2.5 KCl, 1.2 NaH₂PO₄, 5 sodium ascorbate, 3 sodium pyruvate, 2 thiourea; osmolarity: 303–306 mOsm), continuously bubbled with 95% O₂/5% CO₂. Coronal brain slices (300 µm thick) containing the nucleus accumbens (NAc) were prepared using a Leica VT1200 vibratome at a speed of 0.07 mm/s in the same ice-cold cutting solution. Slices were then transferred to artificial cerebrospinal fluid (aCSF; in mM: 92 NaCl, 20 HEPES, 25 glucose, 30 NaHCO₃, 2.5 KCl, 1.2 NaH₂PO₄, 5 sodium ascorbate, 3 sodium pyruvate, 2 thiourea; osmolarity: 303–306 mOsm) at room temperature, continuously bubbled with 95% O₂/5% CO₂, and incubated for at least 1 hour before recordings. Recordings were conducted at 32°C in a perfusion chamber continuously supplied with aCSF (in mM: 126 NaCl, 2.5 KCl, 1.4 NaH₂PO₄, 1.2 MgCl₂, 2.4 CaCl₂, 25 NaHCO₃, 11 glucose; osmolarity: 303–305 mOsm) at a rate of 1.5–2.0 ml/min using a World Precision Instruments pump. Spontaneous excitatory postsynaptic currents (sEPSCs) were recorded using borosilicate glass microelectrodes (resistance: 2–3 MΩ) filled with a cesium-based internal solution (in mM: 117 cesium methanesulfonate, 20 HEPES, 0.4 EGTA, 2.8 NaCl, 5 TEA-Cl, 4 Mg-ATP, 0.4 Na-GTP, 5 QX-314; osmolarity: 280–285 mOsm). To isolate sEPSCs, neurons were voltage-clamped at –70 mV; to isolate inhibitory postsynaptic currents (IPSCs), cells were clamped at +10 mV. Medium spiny neurons (MSNs) were identified via infrared differential interference contrast (IR-DIC) optics using an Olympus BX5iWI inverted microscope. TdTomato-positive cells were classified based on strong fluorescence signal. Recordings were made using a Multiclamp 700B amplifier (Molecular Devices), filtered at 1 kHz, and digitized at 20 kHz using a Digidata 1440A digitizer (Molecular Devices). Synaptic responses were analyzed with ClampFit (Molecular Devices) and Easy Electrophysiology (Easy Electrophysiology Ltd) software. Series resistance (10–20 MΩ) was monitored via a –5 mV voltage step, and cells exhibiting >20% change in series resistance were excluded from analysis.

### RL modeling

We fit a Q-learning model to the mice’s choices in the probabilistic decision-making task. To ensure near-perfect parameter recovery, for each mouse, we concatenated data from all sessions included in other behavioral analyses into pre- and post-injection datasets. The model estimated trial-by-trial action values for the left and right nose poke actions, Q_left_ and Q_right_, using a single learning rate αq. The model also maintained action history values Kleft and Kright, with an action kernel learning rate αq. At the first trial of each day, we initialized the Q-values and K-values to zero. In addition, we modeled potential side biases using an additional value-like term b to the estimated Q-values of the left side.

On each trial, a decision variable was calculated with the sum of the difference between the left and right action values (weighted by βq), the difference between right and left action history values (weighted by βk), and the side bias term.

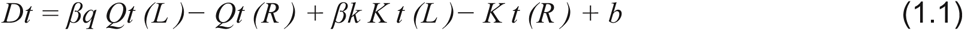

This decision variable was passed to a sigmoidal decision function of the form:

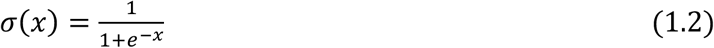

Hence resulting in:

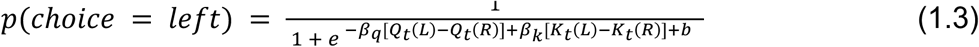

After carrying out the choice, the outcome was delivered (1 for reward, 0 for no reward). The reward prediction error (RPE(t)) was computed by subtracting the value of the chosen action from the outcome:

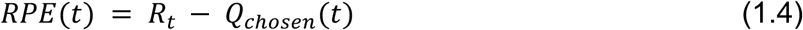

Finally, the value of the chosen action was updated for the next trial by adding RPE(t) weighted by a learning rate α:

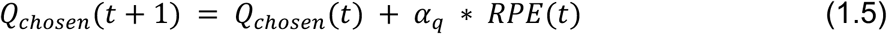

While the unchosen side’s Q values were not updated:

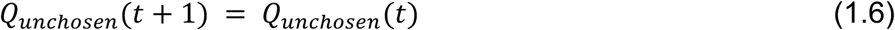

In addition, on each trial, the action kernel was updated with its own learning rate:

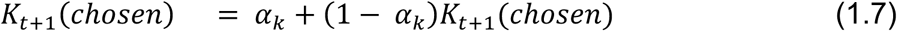

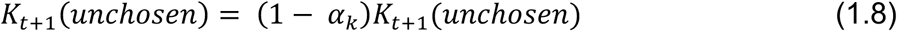

We conducted the standard maximum likelihood fitting using the *p_observed_*, the probabilities of the model matching mice’s action (i.e., *p_observed_* = *p*(*c*ℎ*oice* = *left*) if the observed choice of the animal is left, and *p_observed_* = 1 − *p*(*c*ℎ*oice* = *left*) otherwise). We optimized the log likelihood of the data, *L* = ∑_trials_ *log*(*p_observed_*), using a heuristic global optimization algorithm (Matlab GlobalSearch;(Ugray et al., 2007)). We used GlobalSearch with its default Matlab parameters (NumTrialPoints = 1000, NumStageOnePoints = 200, FunctionTolerance = 1e-6, XTolerance = 1e-6). We initialized the parameters for the initial local optimization process as:

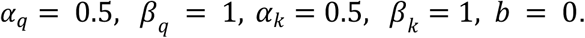

The initial values for subsequent optimization attempts were chosen automatically by the GlobalSearch algorithm. To prevent the optimization from finding unreasonable parameter values, we constrained the search for the parameters as:

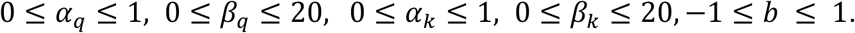

The coefficient of determination, *r*^2^ values, was calculated as:

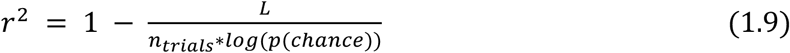

Where, *n_trials_* is the total number of trials in the concatenated data, and *p*(*c*ℎ*ance*) = 0.5. Note that this *r*^2^ value is an equivalent of the *r*^2^ of the linear regression in the case of maximum likelihood fitting and quantifies the model’s ability to predict observed behavior above and beyond chance (Daw, 2009).

#### Version of model with separate learning rates

In addition to this model, which had a single learning rate *α_q_* for all trials, we also fit the data with another model, which had two separate learning rates for win and loss trials: 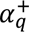 and 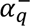, such that:

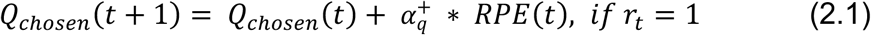

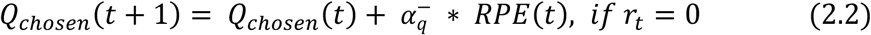

Where *r_t_* denotes the reward in trial t. This model was equivalent to single learning rate model otherwise.

#### Parameter recovery / uncertainty estimation

In order to assess goodness of parameter recovery, and quantify uncertainty in parameter estimations, we ran parameter recovery analysis as follows: For each animal, after fitting the model, we simulated it with the fitted parameter values for *n_sim_* = 100 times. Then, we refit the model to each simulated dataset. This approach provided *n_sim_* = 100 simulation and refit datasets for each animal. We then correlated the simulated parameters with the refitted parameters, which quantified the parameter recovery around the parameters found by the actual fits to the experimental data.

We then used these simulated and refitted datasets to quantify parameter estimation uncertainty. We resampled the inverse temperatures from these refitted parameters, which provided a bootstrap estimate for the uncertainty in parameter estimations.

### Circuit model

We constructed a simplified circuit model to investigate how MSNs integrate cortical task- and motor-related inputs with RPE signals from VTA. To simulate the behavioral paradigm, each trial of the bandit task was divided into three distinct epochs: a 1-second decision period, a 1-second feedback period, and a 5 to 9 second ITI. In the model, corticostriatal synaptic weights are continuously updated, driving action selection based on striatal population activity.

#### Circuit Dynamics and Action Selection

The network consists of 200 MSN neurons, evenly divided into two sub-populations (*N* = 100 each) corresponding to left-action (*L*) and right-action (*R*) choices. The firing rate of the *i*-th left-action neuron at time *t*, denoted as 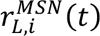, is governed by the following dynamics:

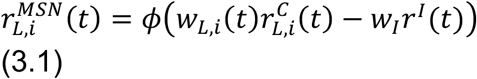

where *w_L,i_*(*t*) represents the average weight of the corticostriatal projection to the *i*-th left neuron, *w_I_* is the weight of local inhibition, and *φ*(*x*) = max(0, *x*) is a rectified linear (ReLU) nonlinearity that ensures non-negative firing rates. MSNs representing the right action, 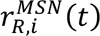, obey the symmetric dynamics:

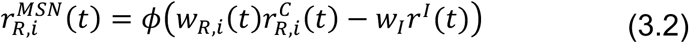

The cortical excitatory drive for MSNs is modeled as a Poisson process. To capture task-dependent modulations, the firing rate of left (right) action neurons is set to 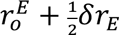 during the decision period, 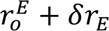 during the feedback period given left (right) action, and 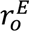 during the ITI period and right (left) action feedback period. Similarly, the local inhibitory input *r^I^*(*t*) follows a Poisson process with a rate of 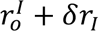 during both the decision and feedback periods, dropping to a baseline rate of 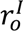 during the ITI.

Downstream structures accumulate this MSN population activity over time with leaky integrators:

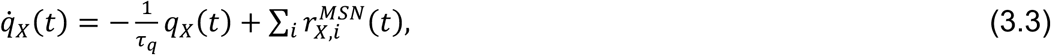

where *t_q_* = 0.2*s*. is the integration decay time constant and *X* = *L* or *R* for left and right integrators *q_L_* and *q_R_*, respectively. Actions are selected probabilistically at the end of the decision period (*t* = *t_d_* = 1*s*) using a softmax choice rule, where the probability of choosing the left action is defined as: *p*(*L*) = 1/(1 + exp[*β*(*q_L_*(*t_d_*) − *q_R_*(*t_d_*))]). Rewards *R*_w_ are delivered according to the bandit schedule, featuring an 80% or 20% reward probability for the preferred and non-preferred ports, respectively.

#### Synaptic Plasticity and Astrocytic Modulation

Corticostriatal synaptic plasticity at synapses ***w****_L_* and ***w****_R_* are formulated based on a learning rule combining a presynaptic eligibility trace of cortical inputs with neuromodulatory feedback. For the *i*-th action synapse, the eligibility trace *e_X,i_* filters the presynaptic activity:

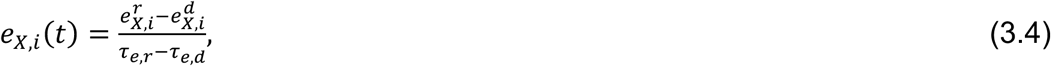

where *X* = *L* or *R* for left and right eligibility traces, respectively. The auxiliary variables 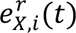 and 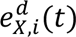 capture the rise and decay kinetics of the eligibility trace, respectively, according to

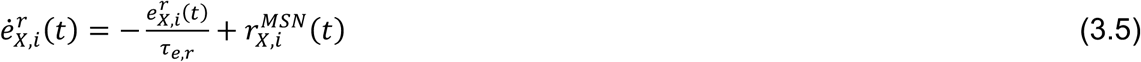

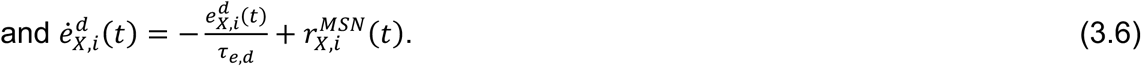

To align with empirical dopamine-dependent plasticity windows in striatum, the rise and decay time constants are set to *τ_e,r_* = 0.5s and *τ_e,d_* = 2.0s (Yagishita et al. 2014).

Dopaminergic inputs to each MSN, denoted as *d_X_*_,*i*_(*t*), deliver a noisy RPE signal during the feedback window:

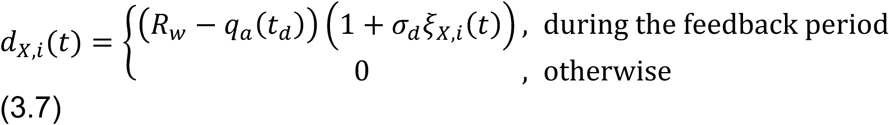

where *a* ∈ {*L*, *R*} is the selected action and *ξ_X_*_,*i*_(*t*) represents standard Gaussian white noise. The continuous synaptic weight update is determined by the product of the eligibility trace and the localized effective neuromodulatory factor *m_X_*_,*i*_(*t*):

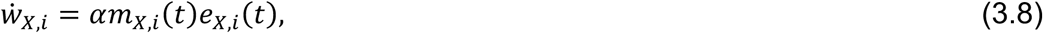

where *α* is the learning rate. In the model without ACD, neuromodulation is localized to direct dopaminergic transmissions, setting *m_X,i_*(*t*) = *d_X,i_*(*t*). By contrast, in the astrocyte-mediated model, we hypothesize the following the spatiotemporal integration of local neuromodulatory signals:

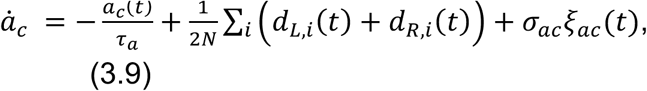

where *τ_a_* = 3s represents the slower astrocytic integration timescale, and the effective neuromodulatory factor is set to *m_X,i_*(*t*) = *a_c_*(*t*). The last term, *σ_ac_ξ_ac_*(*t*) is a noise term where *ξ_ac_*(*t*) is sampled from the standard Gaussian.

#### Excitatory/Inhibitory (E/I) Balance

In slice experiments, we observed that the suppression of astrocytic calcium activity alters the baseline excitatory and inhibitory input firing rates of MSNs. To incorporate this global effect into the model, the baseline firing rates under VS-PMCA are set to 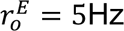 and 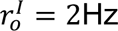. In contrast, the control model with normal astrocyte function uses baseline rates of 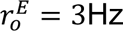 and 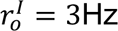.

#### Parameter Optimization and Ablation

While several parameters were constrained by experimental data, five free parameters (*δr^E^*, *δr^I^*, *w_I_*, *α*, and *β*) remained unconstrained. These parameters were optimized to replicate macroscopic animal behavioral statistics by minimizing the following objective function

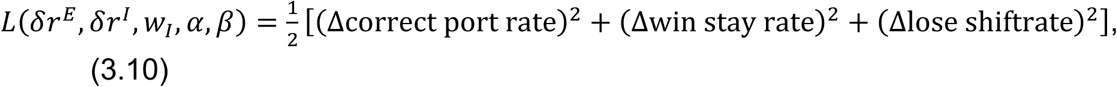

where each Δ term denotes the difference between the average behavioral metric observed in experimental data and that estimated via model simulations. Although the model is underspecified, meaning multiple parameter combinations can successfully replicate the macroscopic behavior, the impact of subsequent model ablations remained robust across optimizations initiated from different random seeds.

For the model ablation analysis (Figure 5E), the five free parameters were first optimized to fit the pre-injection behavioral statistics of VS-PMCA mice (correct port rate: 67%, win-stay rate: 87%, lose-shift rate: 44%). We then conducted three model ablations:

- ’No ac’ model: Eliminates astrocytic integration by enforcing direct neuromodulation (*m_L_*_,*i*_(*t*) = *d_L_*_,*i*_(*t*) and *m_R_*_,*i*_(*t*) = *d_R_*_,*i*_(*t*)) and incorporates the baseline E/I input imbalance, 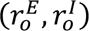 = (5.0, 2.0) Hz.
- ’No sharing’ model: Isolates the lack of spatiotemporal pooling by imposing direct neuromodulation (*m_L_*_,*i*_(*t*) = *d_L_*_,*i*_(*t*) and *m_R_*_,*i*_(*t*) = *d_R_*_,*i*_(*t*)) while maintaining normal baseline E/I inputs: 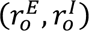 = (3.0, 3.0) Hz.
- ’Imbalance’ model: Isolates the global excitability change by introducing the baseline E/I imbalance, 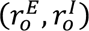 = (5.0, 2.0) Hz, while preserving normal astrocytic neuromodulatory integration (*m_L,i_*(*t*) = *m_R,i_*(*t*) = *a_c_*(*t*)).

#### Implementation Details

At the start of each session, network variables were initialized to 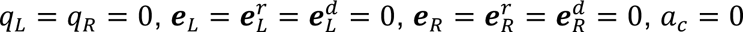, and ***w****_L_* = ***w****_R_* = 0.5. Dynamics were simulated with Euler’s method with timestep Δ*t* = 0.1s. The inter-trial interval was sampled uniformly from 5 to 9 seconds at each trial. The variables were continuously updated over a session spanning 500 trials. To minimize the transient influence of initial value configurations, the first 100 trials of each session were excluded when calculating the correct port rate, win-stay rate, and lose-shift rate.

The noise amplitudes for the dopaminergic and astrocytic dynamics were kept constant at *σ_d_* = 1.0 and *σ_ac_* = 0.5, respectively, throughout all simulations. To allow direct comparison with experimental results (Figures 4, 5), the RPE was post-hoc estimated by fitting a Q-learning model featuring a side bias and a lapse rate directly to the simulated behavioral choices. Finally, the ACD signal was quantified as the mean value of *a_c_*(*t*) during the 1 ≤ *t* ≤ 3.5s time window.

### Additional behavioral assays

#### Sensorimotor battery

The procedures used follow those previously described (J. Chen et al., 2021). For each test the experimenter manually recorded, using a stopwatch, the test time in seconds and hundredths of a second. Two trials were conducted for each test and the average of the two yielded a single time in sec and hundredths of a sec, which was used in the analyses. The order of tests was not counterbalanced between animals so that each animal experienced the test under the same conditions.

In brief, for the walk initiation test, mice were placed in the middle of a 24×24cm square boundary facing away from the experimenter; the time it took for the animals to reach the outside of the boundary was recorded. For the ledge test, animals were placed on a ledge apparatus for 60 seconds; the amount of time they were able to balance on the ledge was recorded. For the inverted screen, animals were placed on a 90 degree inclined screen and assessed for how long they could hold on (up to 60 seconds) For the platform test, animals were recorded for how long they could balance on a wooden platform (measuring 3.5cm thick, 3.0cm in diameter, elevated 25.5 cm above the base) for up to 60 seconds. For the pole test, animals were placed head up on top of the vertical, textured pole apparatus; the time it took for them to turn downward 180 degrees and climb to the bottom of the pole was recorded (up to 120 seconds). For the rotorod, animals were placed on a rotating rod and assessed for how long they could stay on it at either constant or increasing speed (up to 60 seconds).

#### Y-maze

Animals were placed in a Y-maze composed of 3 arms measuring 40cm x 10cm with 20cm high walls, placed at 120 degrees to each other. They were placed in the in the same arm facing away from the center, then allowed to freely explore the arena for 8 minutes. Animal movement was tracked using ANY-maze (Stoelting Co., Wood Dale, IL; http://www.anymaze.co.uk/index.htm). Number of entries and alternations between arms were recorded. Arenas were cleaned with Nolvasan between animals.

#### Novel object recognition

Animals were handled for at least 3 sessions before the NOR was performed. Objects and sides were counterbalanced across animals. Animals were tested in a 40cm x 40cm open field inside sound attenuating boxes. Animals were tested over 3 days for 10 min each. Day 1 consisted of habituation to the environment with no objects; day 2 consisted of habituation to 2 of the same sample objects; day 3 consisted of testing with one sample object and one novel object (2x 10 minute trials with 50 minute ITI). Time animals spent investigating the objects was quantified as time where the animal was oriented towards and within 20mm of an object. NOR index was quantified as the proportion of time spent investigating the novel object relative to all investigation time. Equipment was cleaned with 70% ethanol between animals.

#### Open field

Animals were evaluated in clear 47.6 x 25.4 x 20.6 cm enclosures with lids. They were monitored for 1 hour each using Kinder Scientific Motor Monitor, which tracked distance traveled, rearing behavior, and time spent in the center of the arena (middle 50%). Runs were counterbalanced across mice in different experimental groups. Arenas were cleaned with 70% ethanol between animals.

#### Self grooming

Mice were tested for 10 minutes in transparent enclosures (29.5 x 18.5 x 13cm). Behavior was tracked and quantified for self-grooming (licking, scratching, and nibbling) behavior using Ethovision (https://noldus.com/ethovision-xt). Mice were acclimated to the room for 1 hour prior to testing and enclosures were cleaned with 70% ethanol between animals.

#### Sucrose preference

Testing was performed on single housed mice using automated sucrose preference testing devices (Sippers, (Godynyuk et al., 2019). In brief, animals were single housed for 96 hours and presented with two liquid dispensers that provided either plain water or 1% sucrose. The consumption of sucrose vs plain water was tracked by the Sipper devices over time. The location of sucrose vs water was counterbalanced across animals and switched across days. Prior to testing, animals were presented with plain water in both dispensers.

## Acknowledgements

This work was seeded by the MURI DOD grant to Fabio Pasqualetti, Samet Oymak, Bruno Sinopoli, Thomas Papouin, ShiNung Ching, and Ilya Monosov. Subsequent funding was provided by support from Washington University School of Medicine (to IEM), Johns Hopkins University (to IEM), the National Institute of Mental Health (F31MH135646 to JP), the McDonnell Center for Systems Neuroscience, McDonnell Center for Cellular and Molecular Neuroscience, and the Taylor Family Institute at Washington University in St. Louis. We thank Drs. Bruno Averbeck and Ethan S. Bromberg-Martin for modelling advice, and Drs. Ethan S. Bromberg-Martin and Daeyeol Lee for detailed comments about the manuscript during its preparation. We thank Dr. Robert Gereau for providing necessary resources. The Animal Behavior Core at Washington University performed behavioral testing. We thank Rebecca Mellor for assistance with tissue processing and subsequent tissue imaging.

