## Supplemental Materials and Figures for "Ventral striatal astrocytes contribute to reinforcement learning"

Supplemental Figures.

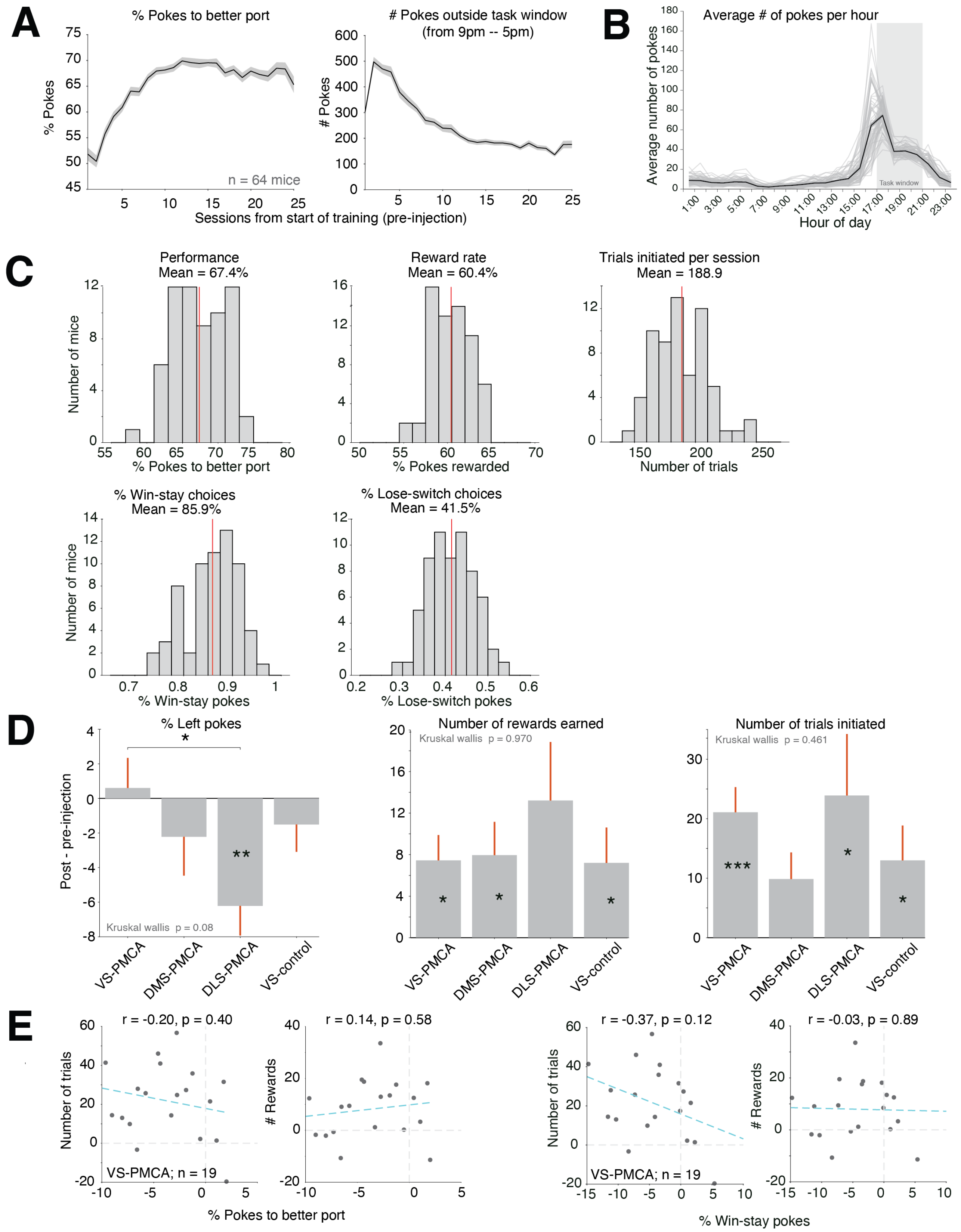

Supplemental Figure 1. Additional behavioral characterizations for free-moving probabilistic decision-making task.

A. Average task performance (% pokes to the better port; right) and number of pokes made outside of the 4 hour task window (left) over pre-injection training sessions for all mice combined. Error bars are S.E.M. across animals. Mice's behavior improve rapidly within the first 5 training sessions.
B. Average number of pokes across the day over pre-injection training sessions for all mice combined. Gray lines show average pokes per hour for individual mice; black line shows average across all mice. Mice show the highest rates of poking in the hour preceding the task period, indicating that most of the pokes made outside of the task period are due to hungry mice anticipatorily poking before the task is made available. VS-PMCA mice did not show a significant change in pokes made outside the task period compared to other groups ($p = 0.81$, Wilcoxon rank sum).
C. Distribution of pre-injection performance, reward rate, trial number, win-stay rate, and lose-switch rate across all mice ($n = 64$).
D. Change in behavioral metrics (% left pokes, number of rewards earned per session, and number of trials initiated per session) pre- vs post-injection for VS-PMCA, DMS-PMCA, DLS-PMCA, and VS-control mice. Similar to other groups tested, VS-PMCA group initiated more choices (trials) per session after the injection compared to pre-injection, suggesting that decreased choice performance in VS-PMCA mice is not simply due to decreased motivation to perform the task. DLS-PMCA injected mice show decreased left bias after injection. Since all injections were bilateral and we did not note systematic biases in hemispheric virus expression, subsequent behavioral experiments will be required to tease out the detailed mechanism for the alteration of the systematic side bias across animals following DLS PMCA injection. For example, future studies ought to explore the relationship of this change in persistent bias and increases in repetitive behavior following ACD attenuation in DLS that have been characterized by Khakh and his colleagues (Yu et al., 2018).
E. Correlation between change in number of trials and change in number of rewards earned against change in task performance (left) or change in win-stay rate (right) for VS-PMCA mice. There is no significant correlation between performance/win-stay vs trial/reward number change pre- vs post-injection (Pearson's r), indicating that mice's decision-making strategy or performance is not directly tied to change in motivation to initiate trials or reward rate.

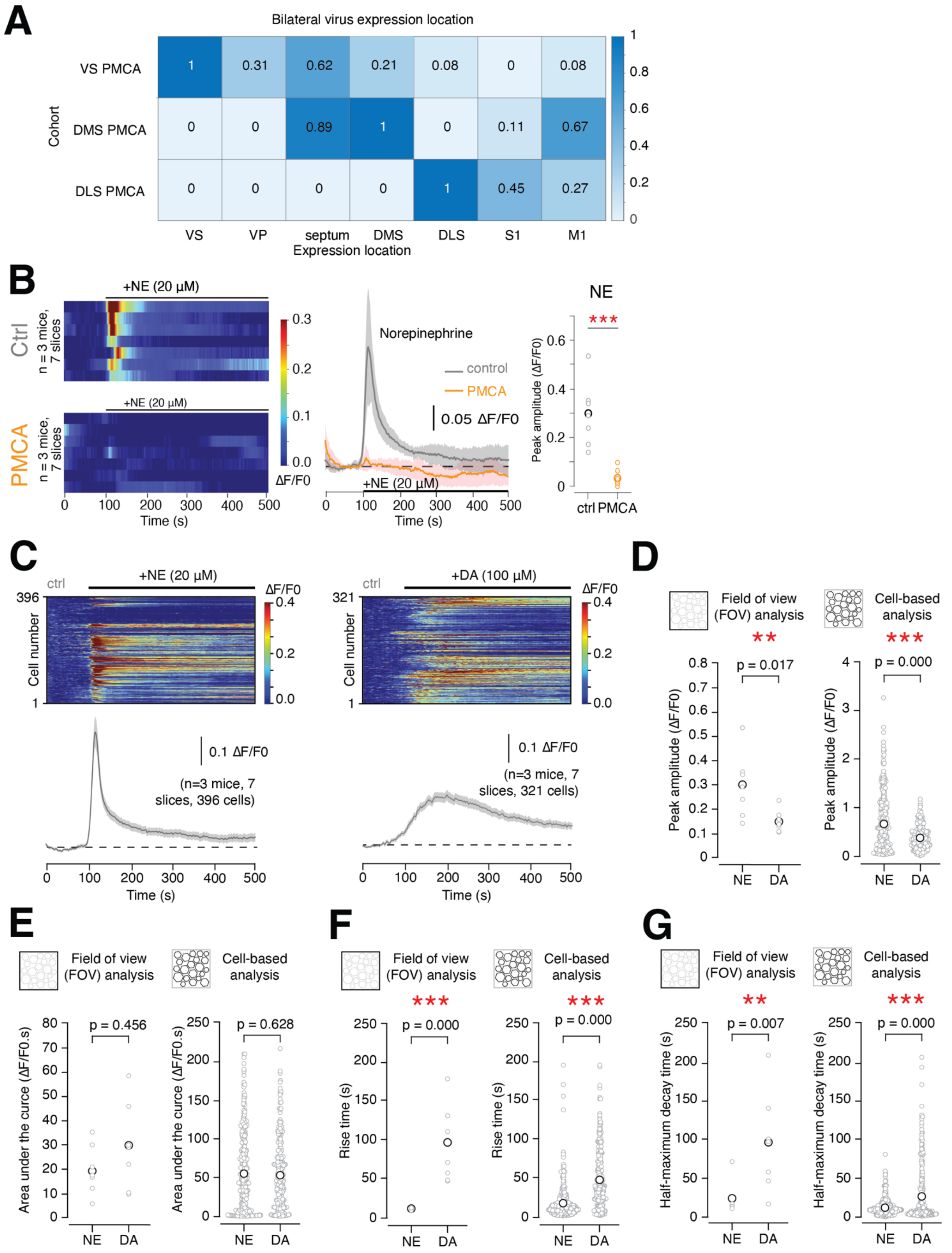
A
Bilateral virus expression location
Cohort
VS PMCA
DMS PMCA
DLS PMCA
VS
VP
septum
DMS
DLS
S1
M1
Expression location
B
+NE (20 µM)
Ctrl
n = 3 mice, 7 slices
PMCA
ΔF/F0
Time (s)
Norepinephrine
control
PMCA
0.05 ΔF/F0
NE
***
Peak amplitude (ΔF/F0)
ctrl PMCA
C
ctrl
+NE (20 µM)
+DA (100 µM)
Cell number
0.1 ΔF/F0
(n=3 mice, 7 slices, 396 cells)
(n=3 mice, 7 slices, 321 cells)
D
Field of view (FOV) analysis
Cell-based analysis
p = 0.017
p = 0.000
NE
DA
E
Area under the curce (ΔF/F0.s)
p = 0.456
p = 0.628
F
Rise time (s)
G
Half-maximum decay time (s)
p = 0.007

Supplemental Figure 2. PMCA attenuates astrocyte calcium responses to both dopamine and norepinephrine in striatum.

A. Quantification of virus spread in striatum and adjacent regions for each injection cohort (VS-PMCA, DMS-PMCA, and DLS-PMCA). Numbers in each square indicate the proportion of animals in that cohort that showed bilateral expression of PMCA in each region. Abbreviations: VS = ventral striatum; VP = ventral pallidum; DMS = dorsomedial striatum; DLS = dorsolateral striatum; S1 = primary sensory cortex; M1 = primary motor cortex.
B. Left: heatmaps of VS astrocyte calcium responses to bath application of norepinephrine (NE) in control (top heatmap) or PMCA (bottom heatmap) slices. NE strongly drives astrocyte calcium responses in several brain regions (Lefton et al., 2025; Paukert et al., 2014; Reitman et al., 2023), and locus coeruleus provides direct noradrenergic innervation of striatum, particularly ventral regions (Berridge et al., 1997; Su et al., n.d.). NE application occurs at 100s. Each row represents one slice's responses to one application; black lines on top indicate the time scale of NE application. Middle: average traces of control or PMCA ACD responses to NE application. Shaded area is 95% confidence interval. Right: comparison of the peak responses to NE (maximum response between 100-300 s) between control and PMCA slices.
C. Heatmap of astrocyte calcium responses to bath application of norepinephrine (left) or dopamine (right) in control slices. Each row represents one cell's responses to one application (see Methods on cell-based analysis; due to lack of astrocyte response in the PMCA condition cell-based segmentation and analysis was not available for PMCA slices). NE or DA application occurs at 100s. Black lines on top indicate the time scale of DA or NE application. Bottom: average traces of responses to NE or DA application. Shaded area is 95% confidence interval.
D. Comparison of the peak responses (maximum responses between 100-300 s) to NE or DA application for control slices, for both field of view (FOV) and cell-based analyses (Wilcoxon rank sum test).
E. Comparison of AUC (100-300 s) to NE or DA application; conventions same as D.
F. Comparison of rise time (time to maximum response between 100-300 s) to NE or DA application; conventions same as D.
G. Comparison of half maximum decay time (time between maximum response and half-max) for NE and DA application; conventions same as D.

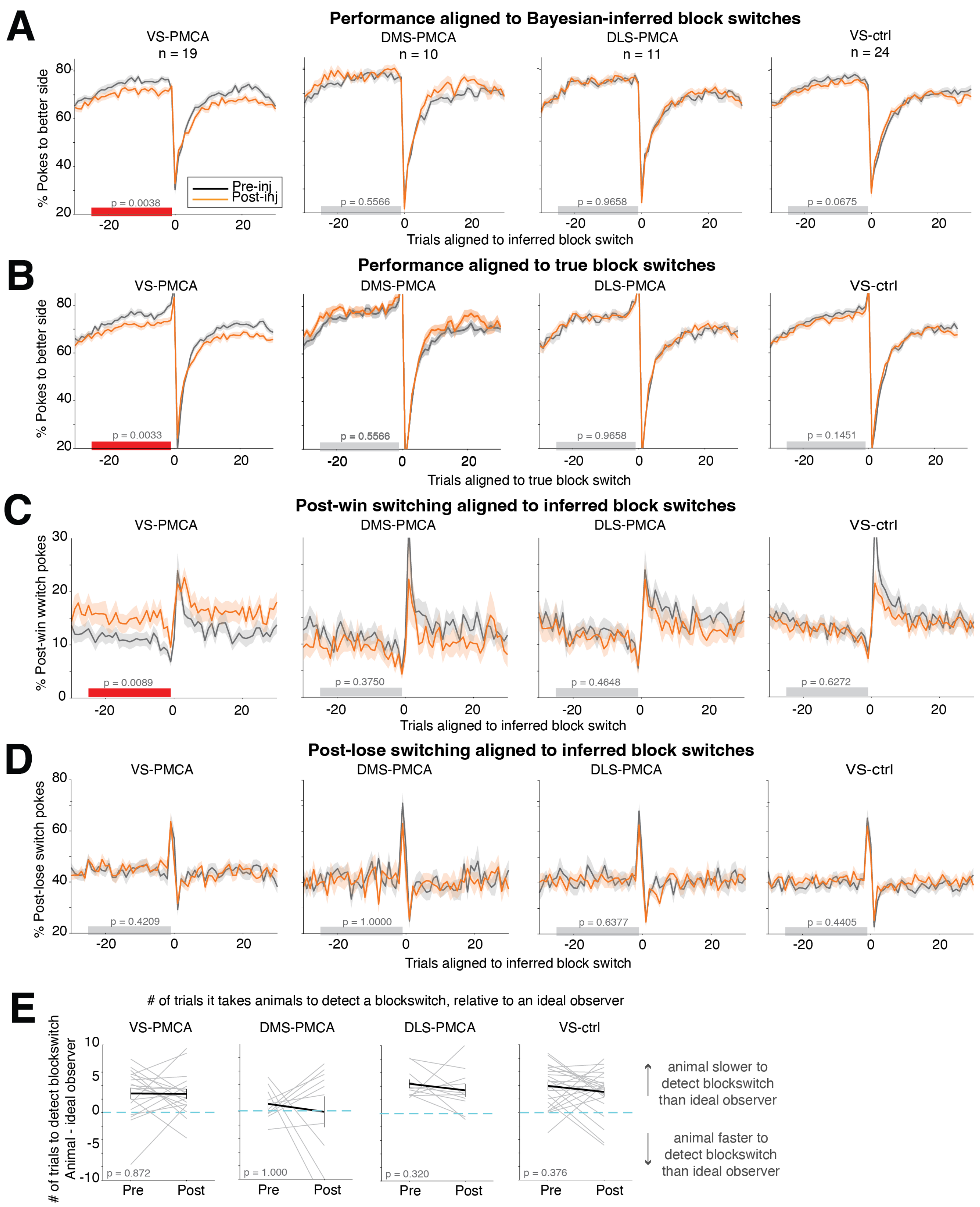
A
Performance aligned to Bayesian-inferred block switches
VS-PMCA n = 19
DMS-PMCA n = 10
DLS-PMCA n = 11
VS-ctrl n = 24
% Pokes to better side
Pre-inj
Post-inj
p = 0.0038
p = 0.5566
p = 0.9658
p = 0.0675
Trials aligned to inferred block switch
B
Performance aligned to true block switches
VS-PMCA
DMS-PMCA
DLS-PMCA
VS-ctrl
p = 0.0033
p = 0.5566
p = 0.9658
p = 0.1451
Trials aligned to true block switch
C
Post-win switching aligned to inferred block switches
% Post-win wwitch pokes
p = 0.0089
p = 0.3750
p = 0.4648
p = 0.6272
Trials aligned to inferred block switch
D
Post-lose switching aligned to inferred block switches
% Post-lose switch pokes
p = 0.4209
p = 1.0000
p = 0.6377
p = 0.4405
Trials aligned to inferred block switch
E
### of trials it takes animals to detect a blockswitch, relative to an ideal observer
### of trials to detect blockswitch Animal - ideal observer
p = 0.872
p = 1.000
p = 0.320
p = 0.376
Pre
Post
animal slower to detect blockswitch than ideal observer
animal faster to detect blockswitch than ideal observer

Supplemental Figure 3. Choice performance and switching aligned to block switches across all cohorts.

A. Average choice performance (% pokes to the better port) across animals aligned to block switches as a Bayesian ideal observer would detect them; for each cohort. Difference tests between pre- and post-injection were calculated in a 25-trial window preceding block switches (gray bar, Wilcoxon sign rank test). Error bars are S.E.M. across animals. All mice achieved a steady-state of choosing the higher valued option (VS-PMCA: 73.9%, DMS-PMCA: 76.4%, DLS-PMCA: 74.4%, VS-control: 74.5%; average % pokes to better port in 25 trial window preceding inferred block switch). None of the cohorts showed a significant difference in pre- vs post-injection performance in the trials immediately following an inferred block switch (5 trial window following block switch, suggesting that the performance deficit in VS-PMCA mice was not specific to the value-updating processes immediately after block switches.
B. Average performance across animals aligned to true block switches for each cohort; conventions follow A. Since most animals were run on a version of the task where blocks switch after 20-30 rewarded trials, note there is a sharp peak in performance prior to true block switches because trials preceding block switches are rewarded. Other than this artifact of task design, results qualitatively and quantitatively align with A.
C. Post-win switching aligned to inferred block switches for all cohorts. DMS-PMCA, DLS-PMCA, and VS-ctrl mice do not show differences between pre-injection and post-injection behavior, compared to VS-PMCA mice (same as Figure 2C). Conventions follow A.
D. Post-lose switching aligned to inferred block switches for all cohorts (VS PMCA results are same as Figure 2C). Conventions follow A.
E. Comparison of how long it took animals to detect block switches, relative to ideal observers. Across all cohorts, it took animals on average 3.0 more trials to detect block switches compared to ideal observers (i.e., if an ideal observer detects a change point 1 trial after the true block switch, on average a mouse will detect a change point 4 trials after the true block switch). This did not differ across cohorts or change as a function of injection (Kruskal-Wallis $p = 0.654$), suggesting that the ability to detect un-cued block switches (which can vary with some parameters of reinforcement learning) was not affected by subregion-specific ACD attenuation.

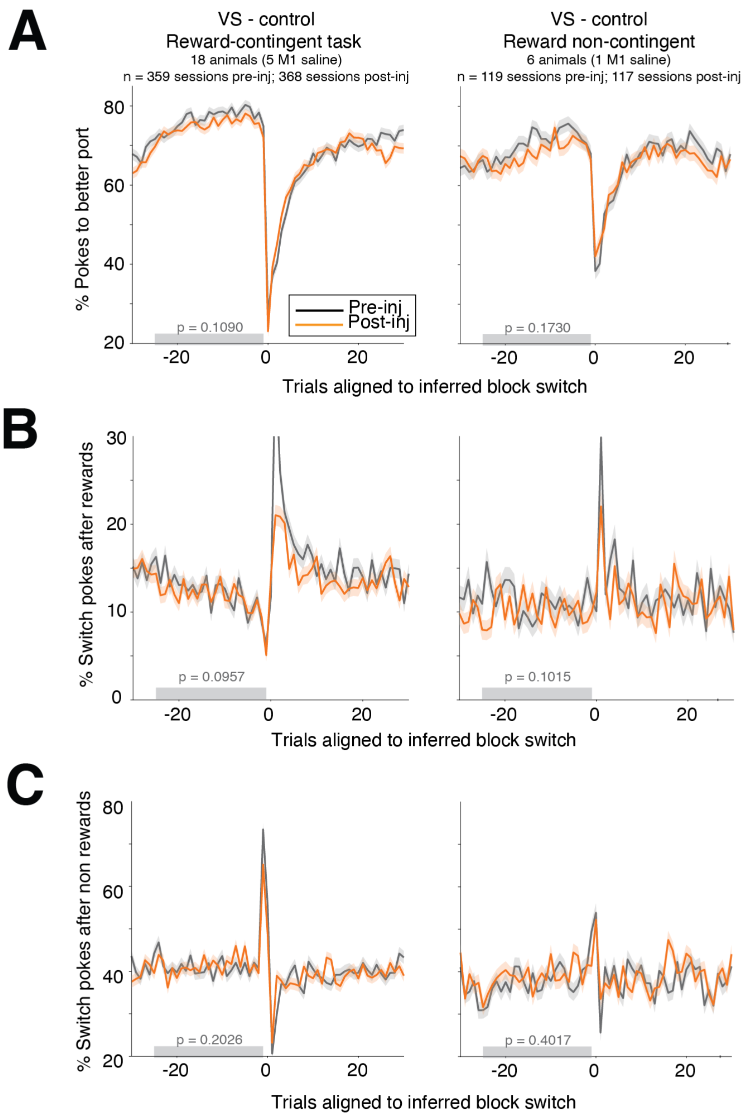


Supplemental Figure 4. Performance and switching aligned to block switches for VS-ctrl mice, split out by reward-contingent and reward-non-contingent block switch versions of the task.

A. Performance aligned to inferred block switches in mice injected with VS-PMCA, split out by animals run on the reward-contingent (blocks switch after 20-30 rewards) task, and animals run on the reward non-contingent (blocks switch after 20-40 trials) task. A subset of animals grouped in VS-ctrl were mice injected with saline in M1, the cortical region in the injection needle trajectory above VS. Because splitting out animals by sub-cohort reduced our statistical power to detect effects, to test if there were differences in behavior pre- vs post-injection we tested across all pre- and post-injection sessions for each sub-cohort. None of these manipulations induced significant changes in correct choice rate, win-stay, or lose-switch behavior. These results suggest that the performance deficit we saw in the VS-PMCA cohort was not due to the injection surgery itself. Lines and error bars are mean and S.E.M. across trials; gray bars indicate 25 trial period pre-block switch where statistical tests (Wilcoxon rank sum test) were run.
B. Switching aligned to inferred block switches, for trials following rewarded trials. Conventions follow A.
C. Switching aligned to inferred block switches, for trials following non-rewarded trials. Conventions follow A.

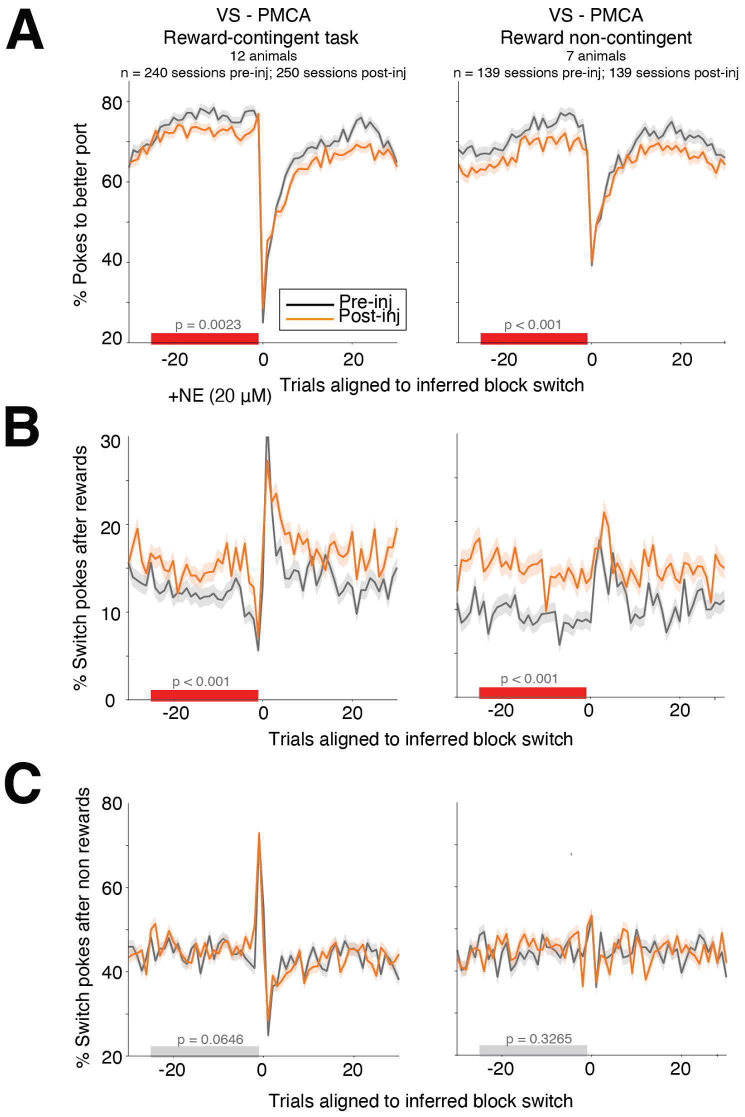


Supplemental Figure 5. Performance and switching aligned to block switches for VS-PMCA mice, split out by reward-contingent and reward-non-contingent block switch versions of the task.

A. Performance aligned to inferred block switches in mice injected with VS-PMCA, split out by those run on the reward-contingent vs reward non-contingent versions of the task. These sub-cohorts were collectively analyzed as ‘VS-PMCA’ mice. Because splitting out animals by sub-cohort reduced our power to detect effects, to test if there were differences in behavior pre- vs post-injection we tested across all pre- and post-injection sessions for each sub-cohort. Attenuating astrocyte calcium in VS reduced performance in mice run on both tasks with different reward contingency meta-structure. Lines and error bars are mean and S.E.M. across trials; gray bars indicate 25 trial period pre-block switch where statistical tests (Wilcoxon rank sum test) were run.
B. Switching aligned to inferred block switches, for trials following rewarded trials. Attenuating astrocyte calcium in VS affects post-reward switching in mice run on both versions of the task. Conventions follow A.
C. Switching aligned to inferred block switches, for trials following non-rewarded trials. There was no significant difference between pre- and post-injection switching for cohorts of mice run on both versions of the task. Conventions follow A.

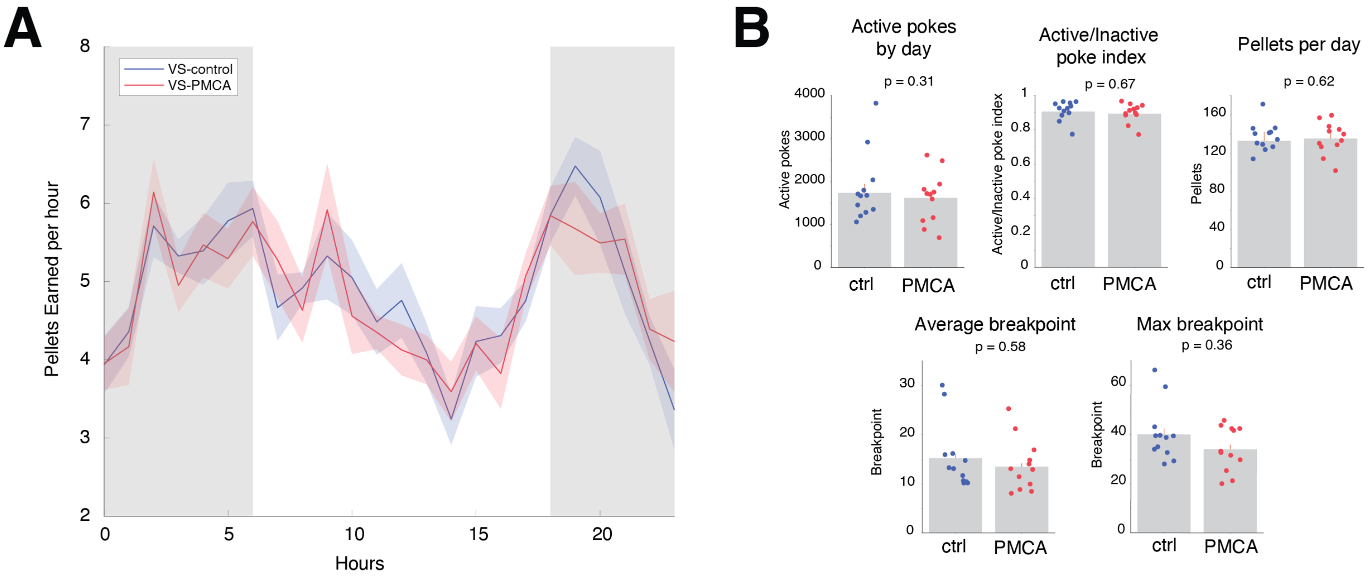


Supplemental Figure 6. VS-PMCA and VS-control mice do not exhibit differences in motivation in a progressive ratio task.

A. Pellets earned per hour for VS-PMCA (n = 12) and VS-ctrl (n = 12) mice in a progressive ratio (PR1) task that was run continuously in the home cages (Methods). We ran the PR1 task continuously to assess not only if motivated behavior differed between groups, but also if circadian patterns in motivation/feeding differed. Unsurprisingly, mice were the most active at engaging with the task and earn the most pellets from 6pm-6am during lights off (gray shaded areas), with two peaks in activity right after lights off and before lights on. Overall, there was no difference in pellets earned or circadian pattern of activity between VS-PMCA and VS-control mice.
B. Commonly used measures of motivation assayed in progressive ratio tasks, including active pokes and pellets earned by day, active/inactive poke index (ratio of pokes to the active vs inactive port), and breakpoint (number of pokes mice are willing to make for one pellet). There was no significant difference between VS-PMCA and VS-control mice across any of these measures.

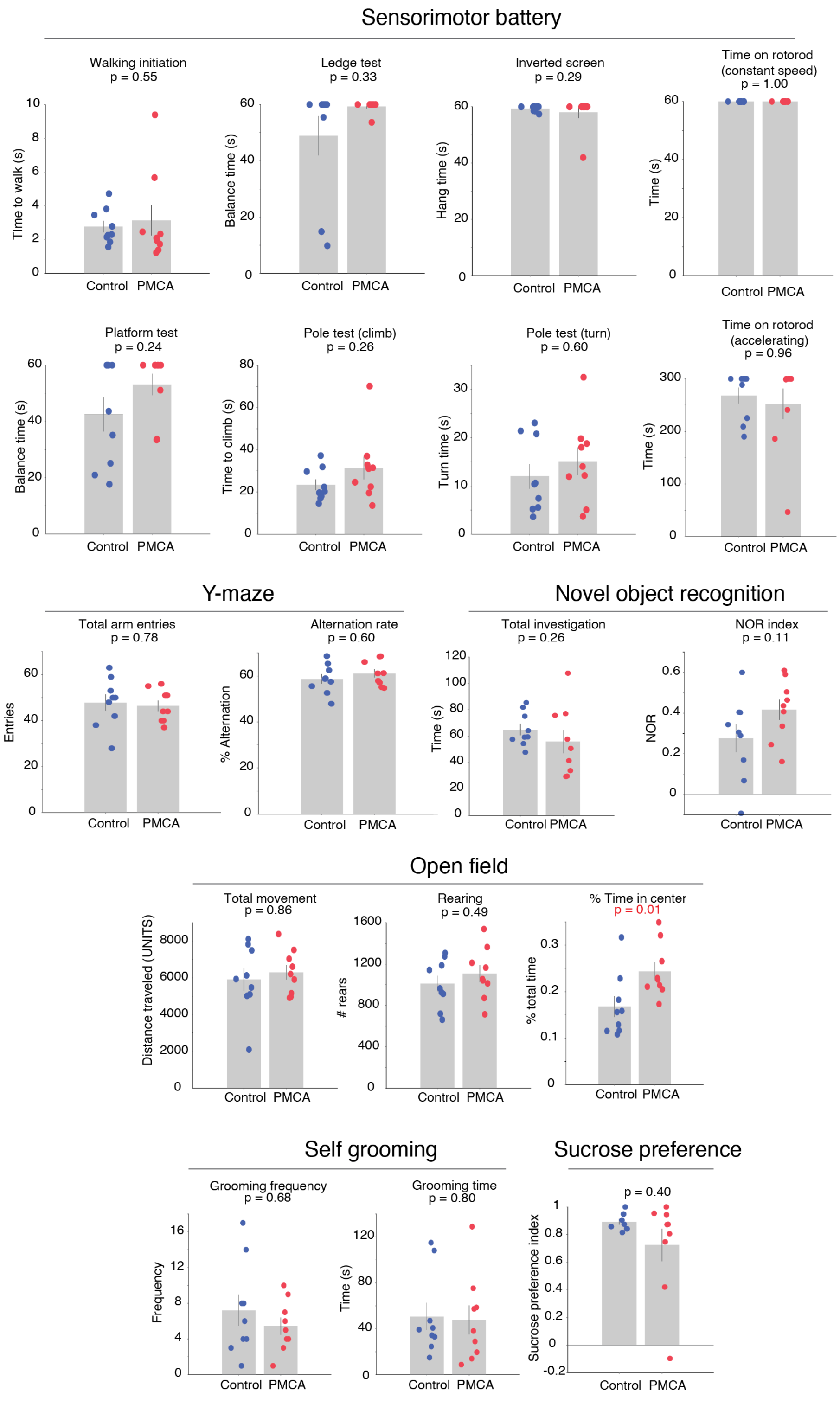


Supplemental Figure 7. VS-PMCA mice do not show deficits in assays of sensorimotor, cognitive, or motivational function compared to VS-control mice.

Naive cohorts of mice were injected bilaterally in VS with either PMCA (n = 9) or control (n = 9) virus and run on a battery of standard behavioral tests commonly used to assay proxies of sensorimotor control, memory, anxiety, cognition, and motivation (see Methods and (J. Chen et al., 2021)).
The only test that revealed a significant difference between cohorts was for the open field test, where VS-PMCA mice spent more time in the center of the arena compared to VS-control mice (uncorrected p value = 0.01; note that with multiple comparison correction considering all assays this effect is not significant). Other measures in this same assay, namely rearing and total movement measures, were not significantly altered. Time in the center of the arena is commonly interpreted as a measure related to less anxiety-like pro-exploratory phenotype (Seibenhener & Wooten, 2015). However, this phenotype may arise due to many sources (e.g., including changes in valuation and expectation that also give rise to win-stay deficits) and therefore must be interpreted with caution. In VS PMCA cohort, there were no significant differences in other assay-measures of sensorimotor coordination and strength, memory (Y-maze and novel object recognition), repetitive behavior (self-grooming), or hedonia (sucrose preference). Altogether, these data suggest that VS-PMCA mice do not show obvious decreases in motivation, memory, or sensorimotor ability. P-values are uncorrected for multiple comparisons.

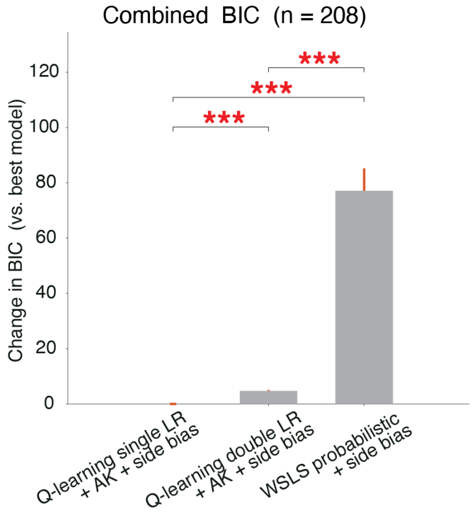


Supplemental Figure 8. Animal behavior is better fit by a Q-learning model than win-stay lose-shift models.

A. Bayesian information criterion (BIC) values for a subset of models (Q-learning model with single learning rate + action kernel + side bias; Q-learning model with two learning rates for positive and negative outcomes + action kernel + side bias term; probabilistic win-stay lose-shift model with side bias term) tested during model comparison (complete list of models in Table 3). The Q-learning model with a single learning rate, action kernel, and side bias term provided the most parsimonious fit to behavior accounting for model fit and number of free parameters.

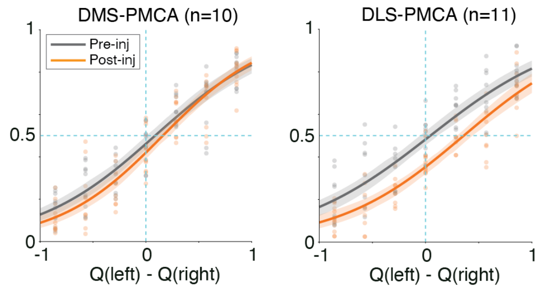


Supplemental Figure 9. Psychometric function shifts for all cohorts pre- and post-injection. Psychometric functions of DMS-PMCA and DLS-PMCA mice pre- and post-injection. Dots are individual animals' left choice probabilities across different inferred Q-value differences; lines are average fitted sigmoid curves for pre-injection (black) and post-injection (orange) sessions across animals. DLS-PMCA animals show a rightward shift in their psychometric functions, consistent with increased rightward side bias seen in their raw behavior (Supplemental Figure 1D) and their model fits (Supplemental Figure 11). Since all injections were bilateral and we did not note systematic biases in hemispheric virus expression, subsequent behavioral experiments will be required to tease out the detailed mechanism for the alteration of the systematic side bias across animals following DLS PMCA injection. Error bars = S.E.M. across animals.

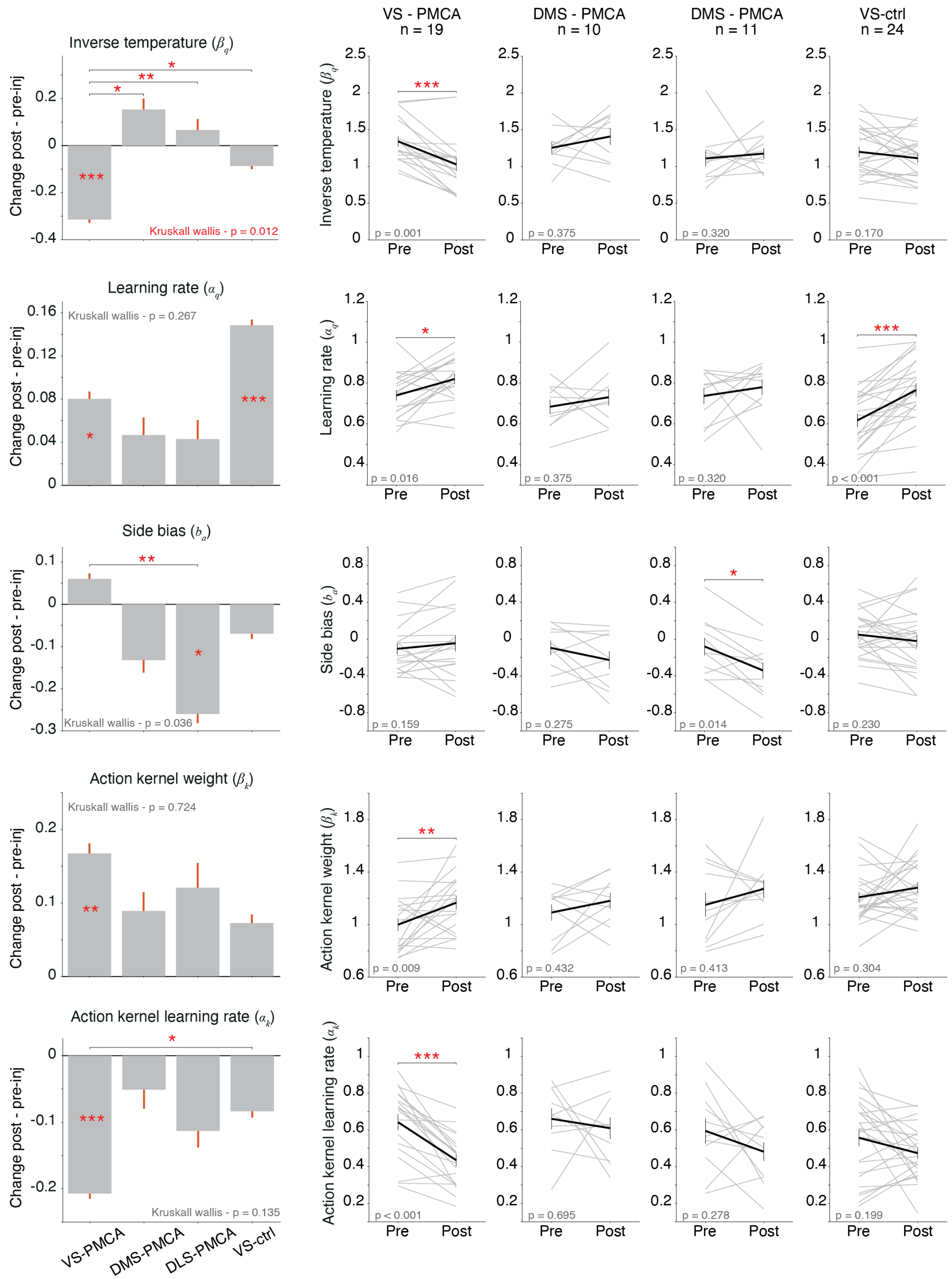


Supplemental Figure 10. Q-learning model parameter shifts for all cohorts pre- and post-injection.

Q-learning model parameter fits for all cohorts. Left subplots: Average change in model parameters pre-vs post-injection for each cohort; error bars are S.E.M. across animals. Comparisons across groups are Wilcoxon rank sum. Within group tests are Wilcoxon signed rank (same as right subplots). Right subplots: Change in model parameters. Inverse temperature values for VS-PMCA and VS-ctrl. Gray lines are individual animals' inverse temperature parameter fits; black lines are average across animals. Error bars are S.E.M. across animals. Within-animal comparisons for each cohort are Wilcoxon signed-rank. P-values are uncorrected for multiple comparisons.
DLS-PMCA mice showed changes in model side bias parameter, consistent with shifts in their overall side bias poking (Supplemental Figure 1D).
VS-PMCA mice showed a significant decrease in action kernel learning rate ($\alpha_k$), indicating slower trial-to-trial updating of K-values. This is consistent with a decrease in choice persistency. VS-PMCA mice also show an increase in action kernel weight ($\beta_k$), although this result was barely significant not statistically significant across groups.

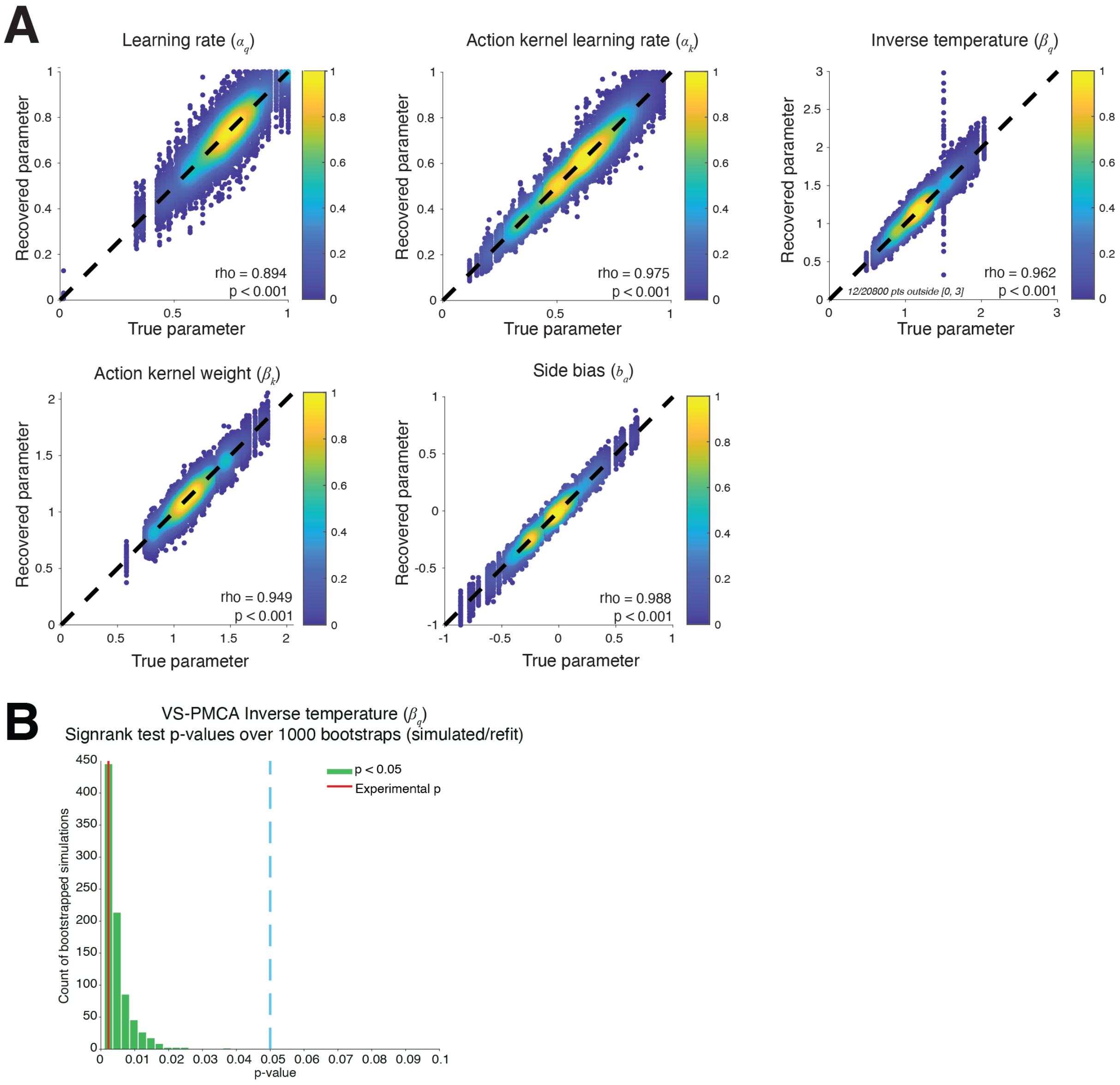


Supplemental Figure 11. Q-learning parameter uncertainty estimation shows strong convergence to fitted parameters.

A. Parameter recovery analysis for Q-learning model parameters. For all animals and pre- and post-injection model fits, we simulated models with the fitted parameters 100 times and refit the model to these simulated data. We then plot the recovered parameters against the initial fit parameters to gain an estimate of the parameter fitting uncertainty. The very high correlation between true and recovered parameter values indicate the reliability of the estimated parameters around the actual fitted values. *Dashed line:* identity. Colors indicate a rank-based density percentile of scatter points at each x and y value, estimated by using a 2-D Gaussian kernel density (Matlab mvkdensity), then converted to a density percentile, hence 0 is sparsest and 1 is the densest region (analogous to Python seaborn.kdeplot)
B. Sign rank p-values run for VS-PMCA inverse temperature pre- vs post-injection fits using recovered model fits over 1,000 bootstraps. 100% of p-values from these tests were significant ($p<0.05$),

indicating the reliability of the result that VS-PMCA mice show significantly decreased inverse temperature pre- vs post-injection (Figure 3**I**, Supplemental Figure 10).

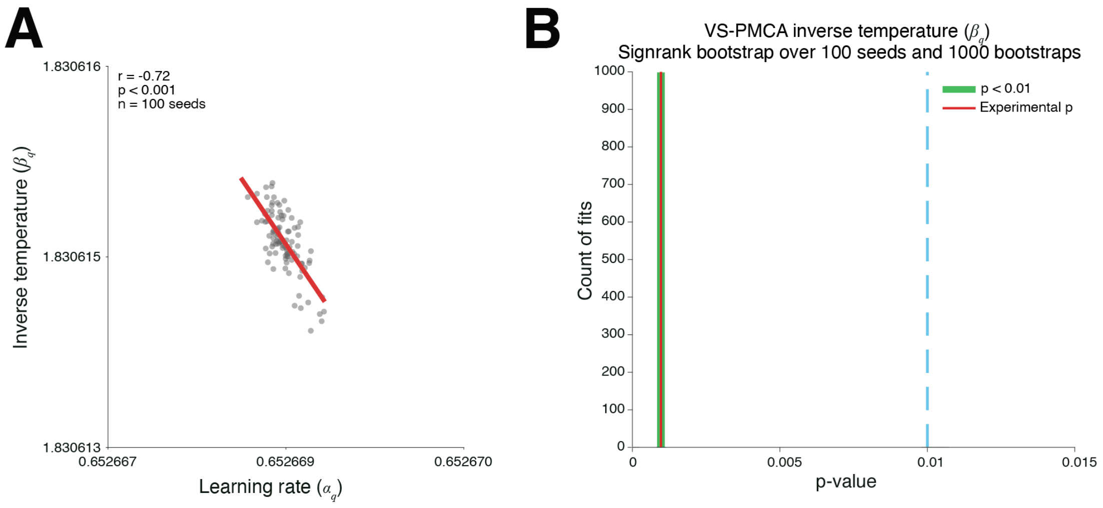


Supplemental Figure 12. Learning rate and inverse temperature show parameter reliance, but within a quantitatively constrained range.

A. Example of covariance between learning rate and inverse temperature for an animal. To quantify the correlation between the learning rate and inverse temperature, we refit the models to the data n = 100 times, each time using a different random seed and a randomly drawn initial value for each parameter. Note that the range in variation across both parameters is exceedingly small; this is because the global optimization method used during model fitting makes the fitting robust to different initializations of the model (see Methods).

B. Histogram of sign rank p values for VS-PMCA pre and post injection, bootstrapped across n =100 fits. Note that the histogram is almost entirely concentrated at the experimental value; similar to the narrow range shown in A, this is due to the fact that the global optimization-based fitting finds very similar fitted parameters across different random seeds and initializations.

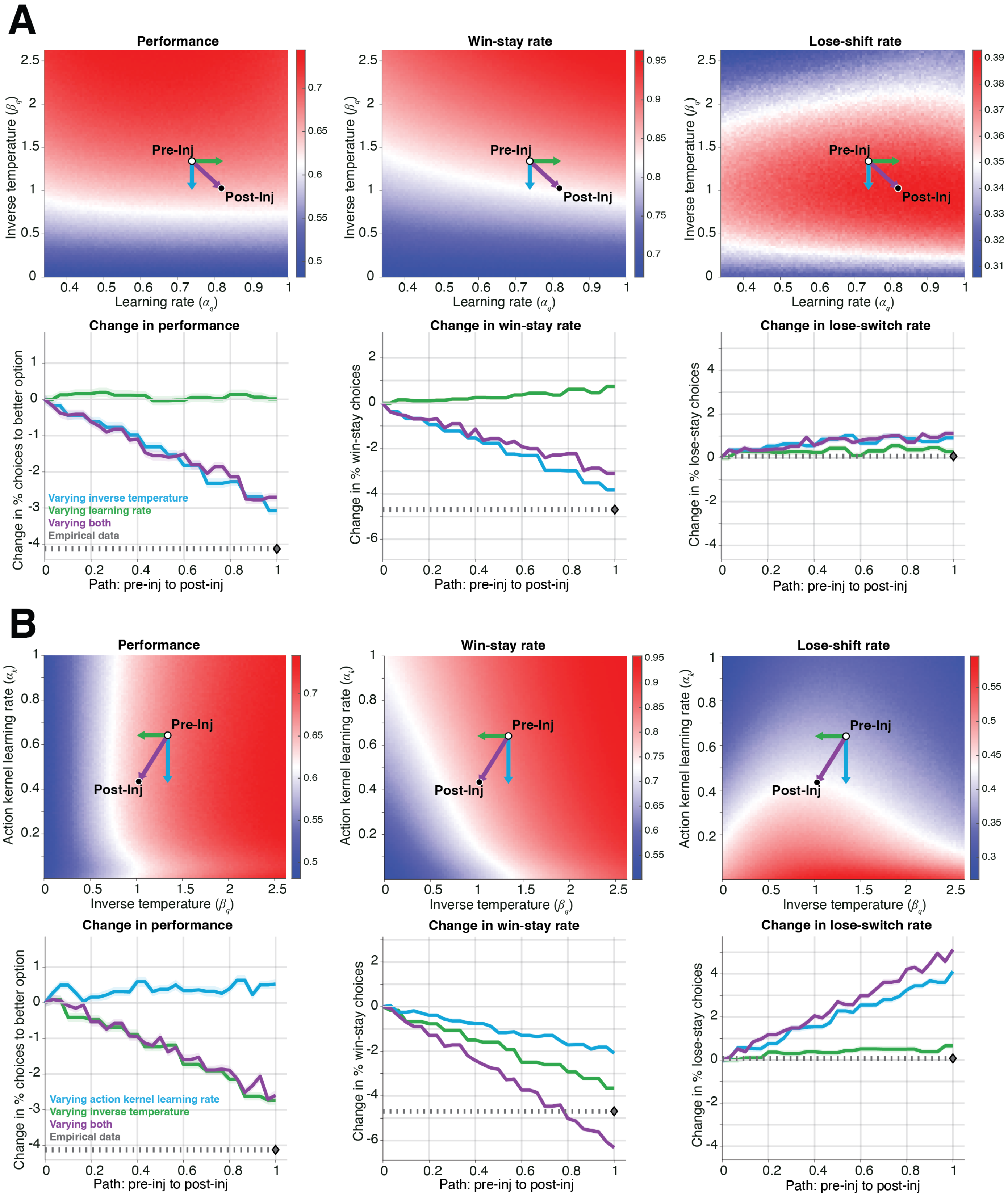
A
Performance
Win-stay rate
Lose-shift rate
Inverse temperature ($\beta_q$)
Learning rate ($\alpha_q$)
Pre-Inj
Post-Inj
Change in performance
Change in win-stay rate
Change in lose-switch rate
Change in % choices to better option
Change in % win-stay choices
Change in % lose-stay choices
Path: pre-inj to post-inj
Varying inverse temperature
Varying learning rate
Varying both
Empirical data
B
Performance
Win-stay rate
Lose-shift rate
Action kernel learning rate ($\alpha_k$)
Inverse temperature ($\beta_q$)
Pre-Inj
Post-Inj
Change in performance
Change in win-stay rate
Change in lose-switch rate
Change in % choices to better option
Change in % win-stay choices
Change in % lose-stay choices
Path: pre-inj to post-inj
Varying action kernel learning rate
Varying inverse temperature
Varying both
Empirical data

Supplemental Figure 13. Changes in inverse temperature best describe changes in VS-PMCA decision-making within the parameter regime fitted to animals' behavior.

A. Changes of model's performance, win-stay rate, and lose-switch rate as inverse temperature ($\beta_q$) and learning rate ($\alpha_q$) are changed. To understand which parameter changes can recapitulate the behavioral changes we observed, we simulated the RL model, drawing parameter values around the fitted values of pre and post injection. For each pair of 5 model parameters, we made 2D grid, by getting the maximum and minimum values observed in actual fittings, extended it by 50% in both upper and lower bounds, and clipped it to the given parameters' valid bounds. We then sampled 100 evenly- spaced values from that range, hence resulting in a 100x100 grid for each parameter pair around the actual fitted values. In each simulation grid, we kept the other 3 parameters of the model fixed at the population mean of the actual fitted values of VS-PMCA. We ran each simulation 200 times to lower stochasticity in model behavior, and obtained the mean performance, win-stay and lose shift values across these simulations for each parameter value in each grid.
*Top:* Heatmaps, showing performance, win-stay and lose-shift of the model simulated across the grid. White dot shows the population mean of VS-PMCA group pre-injection; and black dot shows the population mean of the VS-PMCA group post-injection. Green arrow shows the change in the parameter in x axis (here ($\alpha_q$ )) from pre to post injection values while the parameter in y axis is kept the same (here ($\beta_q$)); blue arrow shows the change in the parameter in y axis (here ($\beta_q$)) from pre to post injection values while the parameter in y axis is kept the same (here ($\alpha_q$ )). The purple arrow shows the composite change of the two parameters from pre to post injection.
*Bottom:* Change in performance, win-stay and lose-shift of the model, as the 2-D grid in the heatmap in top is sliced through the directions of green, yellow and purple arrows. Green line: change in the direction of green arrow, blue line: change in the direction of blue arrow, purple line: change in the direction of purple arrow. Gray dashed line: change in the experimental data.

B. Changes of model's performance, win-stay rate, and lose-switch rate as the inverse temperature ($\beta_q$) and action kernel learning rate ($\alpha_k$) are changed. Conventions follow A.

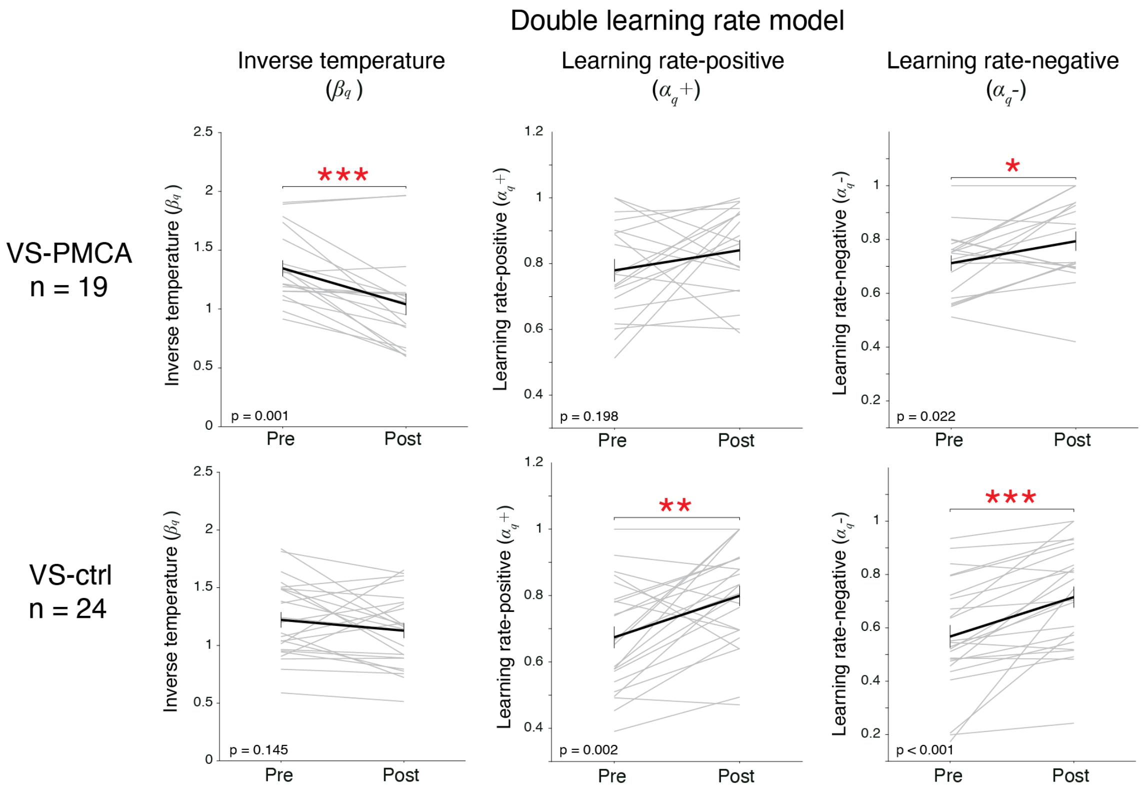


Supplemental Figure 14. Inverse temperature significantly decreases in VS-PMCA mice when fitted with a Q-learning model with separate learning rates.

Inverse temperature and learning rate parameter fits for VS-PMCA and VS-ctrl mice from a variation of the Q-learning model with separate learning rates for rewarded and non-rewarded trials (Methods). Gray lines are individual mice, black line is average over mice. Statistical tests within cohorts are Wilcoxon signed rank. Similarly to the version of the model with one learning rate, VS-PMCA showed significantly decreased inverse temperature, and VS-ctrl mice (and other cohorts; data not shown) did not. The difference in effect between VS-PMCA and VS-ctrl mice (Wilcoxon rank sum) $p = 0.051$.

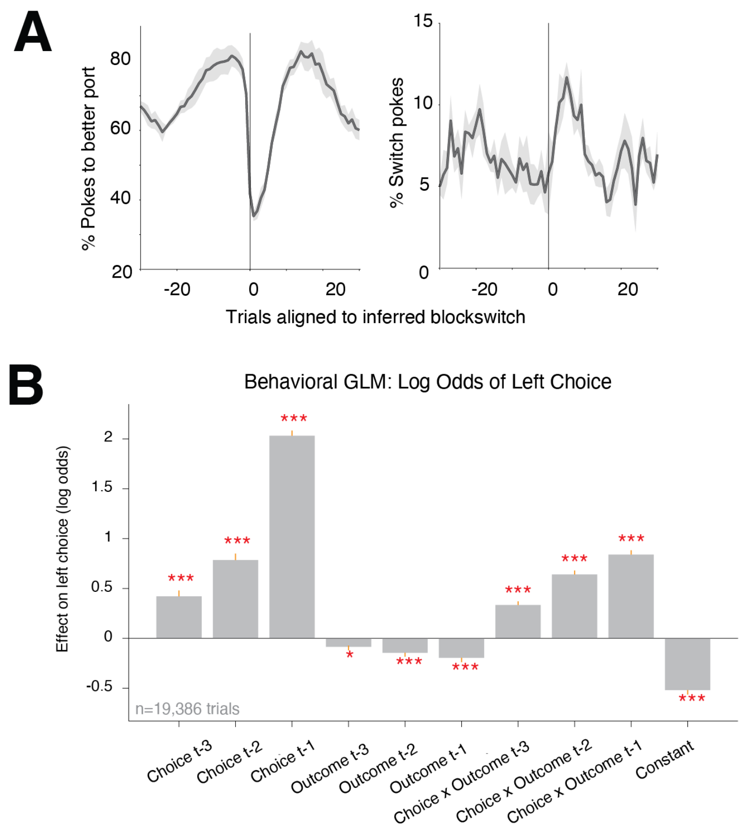


Supplemental Figure 15. Performance and GLM models for behavior in a head-fixed probabilistic decision-making task.

A. Choice performance (left) and switching (right) behavior for mice in the head-fixed decision-making task, aligned to Bayesian-inferred block switches. Thick line is averaged behavior over animals (averaged over all blocks for each animal first); error bar is S.E.M. over animals.
B. Mean fitted GLM beta weights of regressors representing past trial choice (Left: 1, Right: -1), outcome (Rewarded: 1, Unrewarded: -1), Choice x Outcome, and side bias (constant) on the log odds of choosing left in the current trial. The model was fitted to trials pooled from all included behavioral sessions (Methods) using a logistic link function. We included separate regressors representing those variables 1, 2, and 3 trials prior. In this task Past Choice weights were high because mice often made the same choice as in previous trials, indicating a choice persistency effect that is strongest for the most recent choice history. Crucially, the interaction between Past Choice x Outcome had additional predictive value beyond just Past Choice, with larger effects of more recent outcomes. This is a key signature of reinforcement learning, indicating that animals learned by adjusting their choices based on their consequences. Finally, individual mice had some degree of side bias in their behavior. On average, mice modestly preferred the right option indicated by a negative weight of Constant, and this appears to be stronger following a reward, indicated by negative weights of Past Outcome. Error bars are 1 SE; asterisks represent significance of each weight.

Calcium Aligned to Next Trial Tone

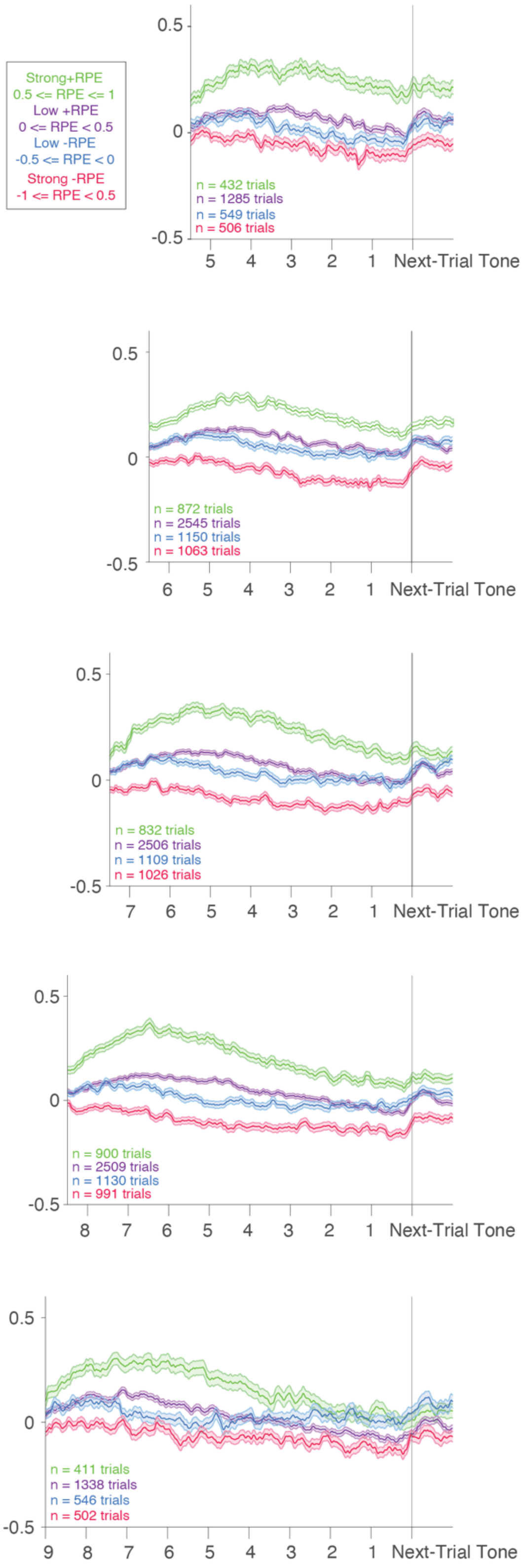

#### Supplemental Figure 16. Ventral striatal astrocyte responses to outcome prediction errors over long inter-trial intervals.

Astrocyte calcium responses split out by RPE (conventions following Figure 4F) aligned to the start of the next trial. From top to bottom, showing responses to trials where the ITI ranged from short (5 seconds) to long (9 seconds). Calcium transients, particularly to strong positive RPEs, remained elevated for 5-7 seconds after the outcome, which led to an elevated baseline calcium in trials with shorter ITIs.

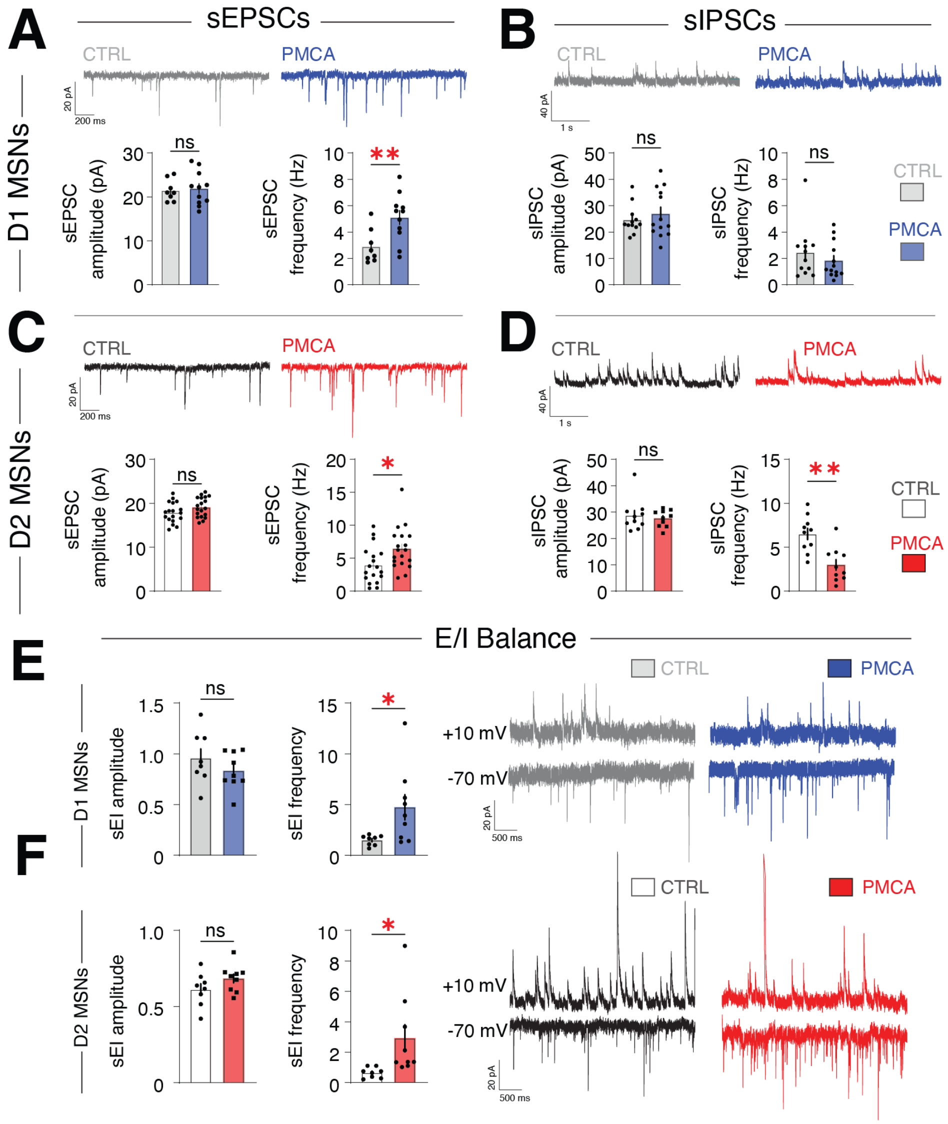


Supplemental Figure 17. Attenuation of astrocyte calcium disrupts striatal excitatory-inhibitory balance.

A-B. Bottom, histograms of the means obtained from amplitudes (left, a measure of post-synaptic efficacy) or frequencies (right, a measure of pre-synatic neurotransmitter release) of sEPSCs (A) and sIPSCs (B) recorded from VS D1-MSNs brain slices from PMCA (sEPSC: n = 11 cells, 5 mice; sIPSC: n = 13 cells, 5 mice) or control mice (sEPSC: n = 8 cells, 4 mice; sIPSC: n = 12 cells, 4 mice). Top, example traces. D1 MSNs exhibited significantly increased measures of pre-synaptic excitatory drive in PMCA slices compared to control. Scale bar, 200 ms, 20 pA.

C-D. Bottom, histograms of the means obtained from amplitudes (left) or frequencies (right) of sEPSCs (C) and sIPSCs (D) recorded from VS D2-MSNs in brain slices from PMCA (sEPSC: n = 19 cells, 6 mice; sIPSC: n = 10 cells, 4 mice) or control mice (sEPSC: n = 18 cells, 6 mice; sIPSC: n = 10 cells, 4 mice). Top, example traces. D2 MSNs exhibited significantly increased measures of pre-synaptic excitatory drive and significantly decreased measures of pre-synaptic inhibitory drive in PMCA slices compared to control. Scale bar, 1 s, 40 pA.

E-F. Comparison of E/I ratio between PMCA (D1: n = 9 cells, 5 mice; D2: n = 9 cells; 4 mice) and control mice (D1: n = 8 cells, 4 mice; D2: n = 8 cells; 4 mice) obtained from NAc D1-MSNs (**E**) or D2-MSNs (**F**). Right, representative traces of sEPSCs at −70 mV and sIPSCs at +10 mV recorded from the

same neurons. Both D1- and D2-MSNs show significantly increased relative excitatory drive in PMCA compared to control slices. Scale bar, 500 ms, 20 pA. Data are mean ± S.E.M.
ns, not significant.

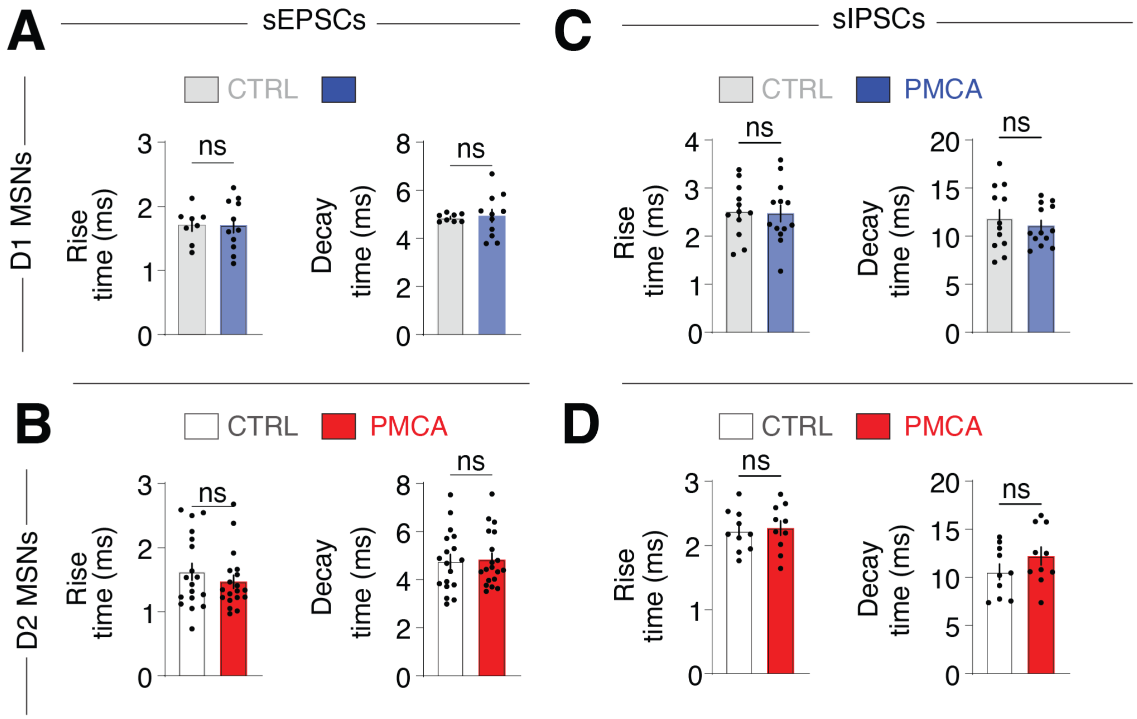


Supplemental Figure 18. Ventral striatum medium spiny neurons do not show differences in post-synaptic temporal dynamics after ACD attenuation.

A-B. Histograms of the means obtained from rise (left) or decay (right) times of sEPSCs recorded from NAc D1-MSNs (A) or D2-MSNs (B) in brain slices from PMCA (D1: n = 11 cells, 5 mice; D2: n = 19 cells; 6 mice) or control mice (D1: n = 8 cells, 4 mice; D2: n = 18 cells; 6 mice).

C-D. Histograms of the means obtained from rise (left) or decay (right) times of sIPSCs recorded from NAc D1-MSNs (C) or D2-MSNs (D) in brain slices from PMCA (D1: n = 13 cells, 5 mice; D2: n = 10 cells; 4 mice) or control mice (D1: n = 12 cells, 4 mice; D2: n = 10 cells; 4 mice). Data are mean ± S.E.M. ns, not significant.

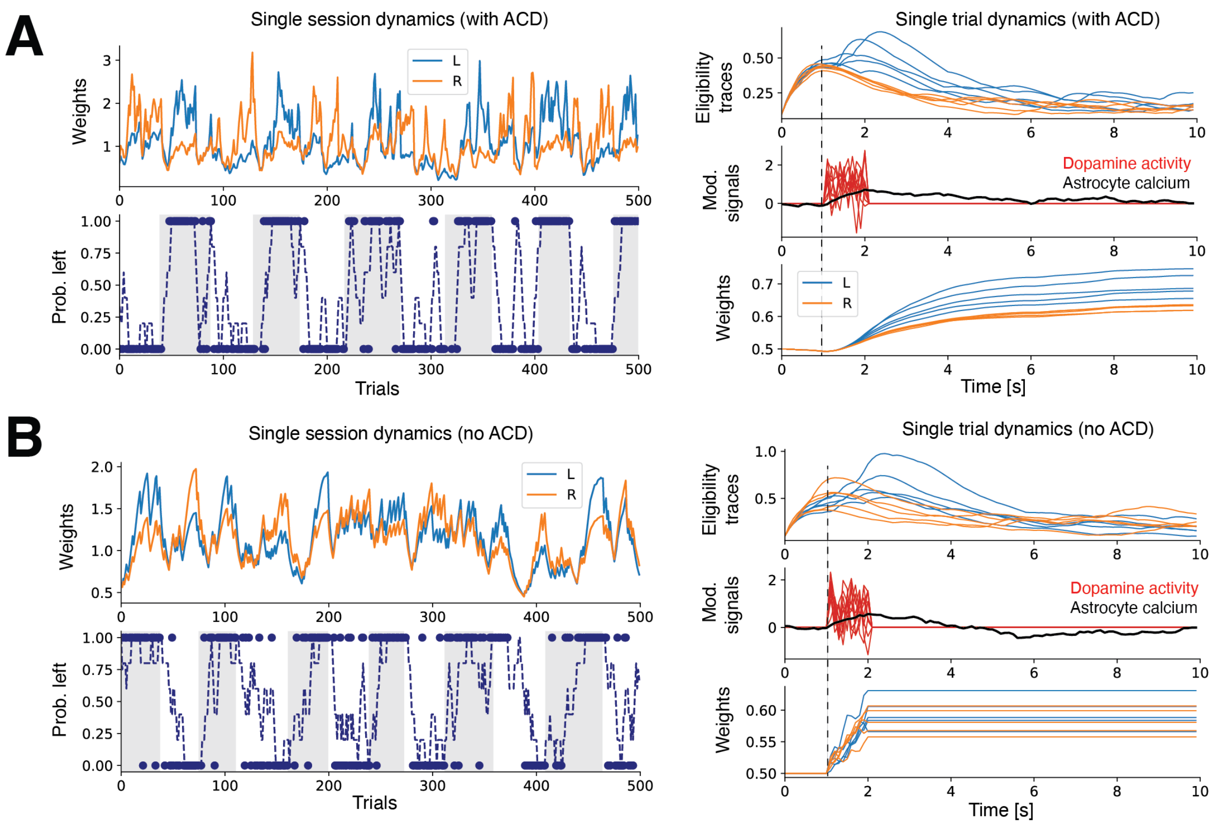


Supplemental Figure 19. Circuit model dynamics under normal and astrocyte ablation conditions.

A. Top left: Model weights for left and right over trials in the normal (no ablation) condition. Bottom left: model behavior in an example session, with left shaded areas indicating periods where left reward probability was higher, dots indicating left (top) or right (bottom) choice, and dashed line indicating rolling average of choice. Right subplots: for a single trial -- eligibility traces for left and right values, dopamine and astrocyte activity, and updated weights, relative to the time of feedback between $1 < t < 2$s.

B. Circuit model dynamics in the astrocyte ablation (E/I imbalance + no RPE share) condition. Conventions follow A. Note that in the no ACD condition, modulation of weights is localized to direct dopaminergic transmissions, without the spatiotemporal smoothing provided by astrocytes.

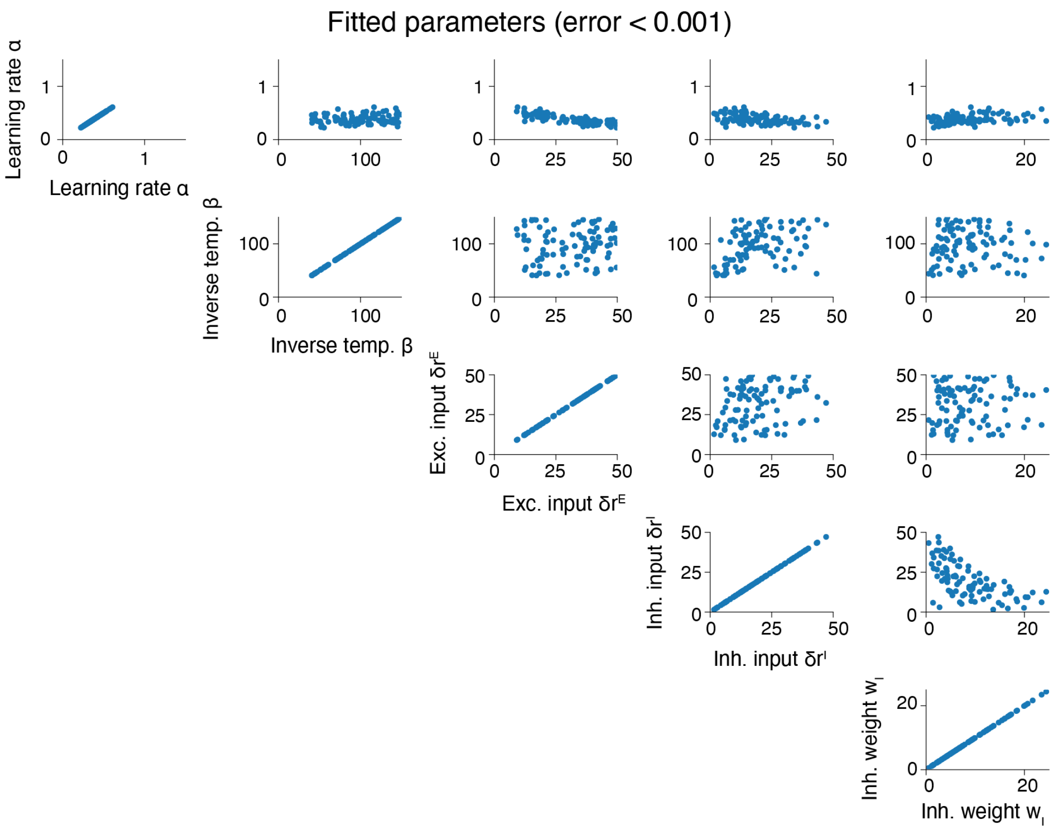


Supplemental Figure 20. Circuit model parameter assessment.

A set of parameters that achieves loss < $10^{-3}$ during model fitting. Note that we optimized five parameters using three macroscopic observations (Eq 3.10). As a result, the model is inevitably underspecified, and the spread of the parameter values reflects this remaining degree of freedom. N = 93 models spanning this set of parameters.

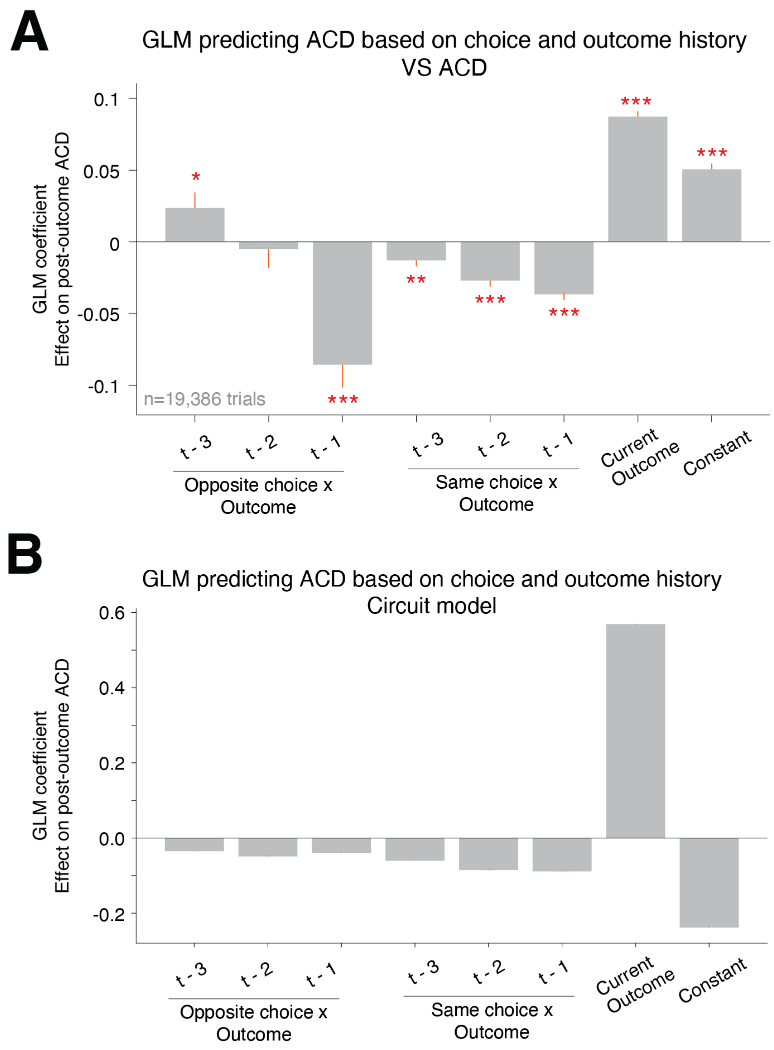


Supplemental Figure 21. GLM model of photometry-acquired ACD and circuit model ACD.

A. Mean fitted GLM beta weights of model fitted to VS astrocyte calcium fluorescence 1-3.5 seconds after outcome delivery. The model was fitted to trials pooled from all included behavioral sessions (Methods), including data from 1-3 trials prior, using a linear link function. The model included beta weights of regressors representing the effects of the current outcome, past outcomes from choices of the same side as the current trial, past outcomes from choices of the opposite side, and a constant term. Crucially, we observed a key signature of RPE coding. Because RPE is roughly expressed as "Actual reward - predicted reward," RPE coding activity in most RL tasks (where animals incrementally build reward predictions about each option based on its past outcomes) should be fit by a positive weight of the current outcome ("Actual reward") and by negative weights of past outcomes from the same choice, with larger negative weights for more recent outcomes ("Predicted reward") (Bayer & Glimcher, 2005). Indeed, this is exactly what we observed in the model's significant positive weight for current outcome and significant negative weights for same choice x outcome ($p<0.001$). The model also had a significant negative effect of the most recent past trial's outcome if the animal had chosen the opposite side on that trial. This likely occurs because this effect is based on a rare subset of trials where animals had previously built up an unusually high reward prediction attracting them to the currently chosen side (as indicated by the fact that they chose the current side even in spite of having just gotten a reward from choosing the opposite side on the previous trial), thus resulting in an unusually negative RPE signal in response to the current outcome.

B. The same GLM, fitted to astrocyte calcium dynamics simulated by the circuit model. Conventions follow A.

Tables.

Table 1. Number of mice in each task cohort.

| Cohort | Total | Female | Male |
|---|---|---|---|
| **VS-PMCA** | **19** | **13** | **6** |
| VS-PMCA<br>Reward contingent task | 12 | 7 | 5 |
| VS-PMCA<br>Reward non-contingent task | 7 | 6 | 1 |
| **DMS-PMCA** | **10** | **3** | **7** |
| **DLS-PMCA** | **11** | **3** | **8** |
| **VS-control** | **24** | **11** | **13** |
| VS-control<br>Reward contingent task | 13 | 6 | 7 |
| VS-control<br>Reward non-contingent task | 5 | 5 | 0 |
| VS-control – M1 saline<br>Reward contingent task | 6 | 0 | 6 |

Table 2. Quantification of key behavioral metrics for each mouse.

Performance = proportion of choices to the better port. Reward rate = proportion of rewarded choices. WinStay = proportion of post-win choices where mouse repeated its previous choice. LoseSwitch = proportion of post-unrewarded choices where mouse switched relative to its previous choice.

| AnimalID | Group | Sex | Performance Pre-inj | Performance Post-inj | Reward Rate Pre-inj | Reward Rate Post-inj | Win-Stay Pre-inj | Win-Stay Post-inj | Lose-Switch Pre-inj | Lose-Switch Post-inj |
|---|---|---|---|---|---|---|---|---|---|---|
| B2_M4 | VS-PMCA | M | 0.72 | 0.647 | 0.621 | 0.589 | 0.941 | 0.887 | 0.42 | 0.396 |
| B2_M6 | VS-PMCA | F | 0.624 | 0.591 | 0.582 | 0.545 | 0.744 | 0.702 | 0.483 | 0.513 |
| B2_M7 | VS-PMCA | M | 0.694 | 0.628 | 0.619 | 0.573 | 0.85 | 0.767 | 0.408 | 0.48 |
| B2_M34 | VS-PMCA | F | 0.675 | 0.662 | 0.613 | 0.596 | 0.863 | 0.828 | 0.489 | 0.488 |
| B2_M35 | VS-PMCA | F | 0.672 | 0.689 | 0.614 | 0.61 | 0.888 | 0.885 | 0.425 | 0.381 |
| B2_M66 | VS-PMCA | M | 0.647 | 0.667 | 0.594 | 0.599 | 0.805 | 0.859 | 0.467 | 0.389 |
| B2_M68 | VS-PMCA | M | 0.677 | 0.619 | 0.603 | 0.568 | 0.859 | 0.786 | 0.442 | 0.468 |
| B2_M70 | VS-PMCA | F | 0.697 | 0.606 | 0.613 | 0.564 | 0.849 | 0.733 | 0.396 | 0.456 |
| B2_M73 | VS-PMCA | M | 0.688 | 0.607 | 0.609 | 0.561 | 0.895 | 0.792 | 0.382 | 0.406 |
| B2_M120 | VS-PMCA | F | 0.69 | 0.662 | 0.611 | 0.607 | 0.918 | 0.872 | 0.427 | 0.411 |
| B2_M124 | VS-PMCA | F | 0.684 | 0.64 | 0.611 | 0.577 | 0.892 | 0.858 | 0.368 | 0.4 |
| B2_M127 | VS-PMCA | F | 0.671 | 0.676 | 0.604 | 0.602 | 0.836 | 0.852 | 0.441 | 0.472 |
| B2_M144 | VS-PMCA | F | 0.668 | 0.65 | 0.609 | 0.589 | 0.833 | 0.837 | 0.476 | 0.457 |
| B2_M145 | VS-PMCA | M | 0.727 | 0.721 | 0.644 | 0.633 | 0.95 | 0.96 | 0.355 | 0.415 |
| B2_M147 | VS-PMCA | F | 0.714 | 0.725 | 0.629 | 0.64 | 0.916 | 0.939 | 0.431 | 0.472 |
| B2_M149 | VS-PMCA | F | 0.717 | 0.652 | 0.636 | 0.592 | 0.916 | 0.804 | 0.489 | 0.469 |
| B2_M150 | VS-PMCA | F | 0.675 | 0.635 | 0.604 | 0.59 | 0.873 | 0.872 | 0.468 | 0.419 |
| B2_M151 | VS-PMCA | F | 0.66 | 0.614 | 0.607 | 0.567 | 0.856 | 0.784 | 0.428 | 0.435 |
| B2_M152 | VS-PMCA | F | 0.688 | 0.592 | 0.617 | 0.552 | 0.872 | 0.725 | 0.527 | 0.493 |
| B2_M13 | DMS-PMCA | M | 0.718 | 0.677 | 0.632 | 0.598 | 0.924 | 0.901 | 0.466 | 0.463 |
| B2_M14 | DMS-PMCA | M | 0.714 | 0.75 | 0.63 | 0.649 | 0.916 | 0.962 | 0.38 | 0.291 |
| B2_M16 | DMS-PMCA | M | 0.734 | 0.697 | 0.634 | 0.622 | 0.909 | 0.87 | 0.427 | 0.413 |
| RP_M2 | DMS-PMCA | M | 0.654 | 0.71 | 0.591 | 0.626 | 0.888 | 0.933 | 0.339 | 0.372 |
| B2_M24 | DMS-PMCA | M | 0.691 | 0.647 | 0.61 | 0.59 | 0.858 | 0.802 | 0.435 | 0.458 |
| B2_M28 | DMS-PMCA | M | 0.644 | 0.677 | 0.59 | 0.611 | 0.776 | 0.911 | 0.481 | 0.34 |
| B2_M61 | DMS-PMCA | F | 0.651 | 0.742 | 0.596 | 0.636 | 0.873 | 0.895 | 0.415 | 0.439 |
| B2_M62 | DMS-PMCA | F | 0.704 | 0.711 | 0.63 | 0.616 | 0.91 | 0.921 | 0.404 | 0.443 |
| B2_M63 | DMS-PMCA | F | 0.65 | 0.634 | 0.582 | 0.58 | 0.805 | 0.781 | 0.471 | 0.423 |
| B2_M65 | DMS-PMCA | M | 0.723 | 0.688 | 0.633 | 0.618 | 0.916 | 0.93 | 0.354 | 0.382 |
| B2_M17 | DLS-PMCA | M | 0.642 | 0.637 | 0.584 | 0.572 | 0.826 | 0.852 | 0.43 | 0.449 |

| | | | | | | | | | |
|---|---|---|---|---|---|---|---|---|---|
| B2_M20 | DLS-PMCA | M | 0.639 | 0.652 | 0.589 | 0.588 | 0.867 | 0.883 | 0.375 | 0.367 |
| B2_M21 | DLS-PMCA | M | 0.65 | 0.635 | 0.585 | 0.586 | 0.832 | 0.874 | 0.442 | 0.408 |
| B2_M23 | DLS-PMCA | F | 0.647 | 0.612 | 0.583 | 0.567 | 0.778 | 0.797 | 0.496 | 0.374 |
| B2_M37 | DLS-PMCA | M | 0.67 | 0.666 | 0.605 | 0.604 | 0.824 | 0.826 | 0.409 | 0.422 |
| B2_M42 | DLS-PMCA | M | 0.72 | 0.65 | 0.625 | 0.581 | 0.923 | 0.871 | 0.404 | 0.376 |
| B2_M51 | DLS-PMCA | M | 0.66 | 0.602 | 0.598 | 0.562 | 0.843 | 0.745 | 0.417 | 0.477 |
| B2_M52 | DLS-PMCA | M | 0.661 | 0.709 | 0.605 | 0.629 | 0.81 | 0.926 | 0.482 | 0.382 |
| B2_M55 | DLS-PMCA | F | 0.642 | 0.706 | 0.586 | 0.612 | 0.858 | 0.904 | 0.397 | 0.473 |
| B2_M69 | DLS-PMCA | F | 0.631 | 0.65 | 0.582 | 0.583 | 0.885 | 0.894 | 0.319 | 0.359 |
| B2_M72 | DLS-PMCA | M | 0.659 | 0.687 | 0.596 | 0.609 | 0.906 | 0.929 | 0.329 | 0.36 |
| B2_M100 | VS-ctrl | M | 0.631 | 0.548 | 0.581 | 0.532 | 0.804 | 0.693 | 0.38 | 0.472 |
| B2_M101 | VS-ctrl | M | 0.657 | 0.715 | 0.59 | 0.62 | 0.804 | 0.905 | 0.519 | 0.48 |
| B2_M102 | VS-ctrl | M | 0.621 | 0.637 | 0.57 | 0.579 | 0.751 | 0.804 | 0.466 | 0.453 |
| B2_M103 | VS-ctrl | M | 0.726 | 0.694 | 0.641 | 0.625 | 0.925 | 0.884 | 0.302 | 0.39 |
| B2_M104 | VS-ctrl | M | 0.743 | 0.637 | 0.64 | 0.586 | 0.922 | 0.824 | 0.357 | 0.389 |
| B2_M105 | VS-ctrl | F | 0.694 | 0.66 | 0.623 | 0.603 | 0.874 | 0.936 | 0.392 | 0.262 |
| B2_M107 | VS-ctrl | F | 0.619 | 0.638 | 0.559 | 0.581 | 0.775 | 0.88 | 0.37 | 0.305 |
| B2_M108 | VS-ctrl | F | 0.713 | 0.717 | 0.628 | 0.643 | 0.928 | 0.88 | 0.343 | 0.438 |
| B2_M113 | VS-ctrl | M | 0.608 | 0.662 | 0.564 | 0.596 | 0.773 | 0.883 | 0.41 | 0.395 |
| B2_M118 | VS-ctrl | M | 0.729 | 0.734 | 0.632 | 0.649 | 0.917 | 0.939 | 0.381 | 0.408 |
| B2_M123 | VS-ctrl | F | 0.663 | 0.63 | 0.594 | 0.584 | 0.852 | 0.819 | 0.41 | 0.426 |
| B2_M126 | VS-ctrl | F | 0.662 | 0.637 | 0.601 | 0.578 | 0.852 | 0.811 | 0.341 | 0.401 |
| B2_M129 | VS-ctrl | F | 0.69 | 0.67 | 0.615 | 0.609 | 0.85 | 0.893 | 0.383 | 0.348 |
| B2_M139 | VS-ctrl | F | 0.667 | 0.659 | 0.594 | 0.595 | 0.883 | 0.897 | 0.446 | 0.383 |
| B2_M140 | VS-ctrl | F | 0.713 | 0.678 | 0.636 | 0.622 | 0.926 | 0.892 | 0.36 | 0.422 |
| B2_M141 | VS-ctrl | F | 0.663 | 0.642 | 0.585 | 0.582 | 0.88 | 0.876 | 0.387 | 0.387 |
| B2_M142 | VS-ctrl | F | 0.654 | 0.671 | 0.579 | 0.599 | 0.837 | 0.916 | 0.371 | 0.392 |
| B2_M143 | VS-ctrl | F | 0.71 | 0.675 | 0.632 | 0.615 | 0.911 | 0.881 | 0.408 | 0.409 |
| B2_M130 | VS-ctrl | M | 0.694 | 0.688 | 0.605 | 0.611 | 0.926 | 0.926 | 0.277 | 0.342 |
| B2_M131 | VS-ctrl | M | 0.638 | 0.658 | 0.585 | 0.6 | 0.795 | 0.823 | 0.419 | 0.421 |
| B2_M132 | VS-ctrl | M | 0.728 | 0.668 | 0.639 | 0.605 | 0.905 | 0.843 | 0.357 | 0.444 |
| B2_M134 | VS-ctrl | M | 0.636 | 0.646 | 0.582 | 0.586 | 0.764 | 0.826 | 0.513 | 0.491 |
| B2_M135 | VS-ctrl | M | 0.634 | 0.665 | 0.579 | 0.591 | 0.775 | 0.809 | 0.418 | 0.5 |
| B2_M137 | VS-ctrl | M | 0.587 | 0.597 | 0.554 | 0.56 | 0.744 | 0.764 | 0.458 | 0.439 |

Table 3. BIC values for all tested models.

| # | Family | Side bias | Lapse | $k$ | AIC (mean) | BIC (mean) |
|---|---|---|---|---|---|---|
| 1 | Static | ✓ | – | 1 | 5208.7 | 5214.9 |
| 2 | Static | – | ✓ | 1 | 5288.7 | 5295.0 |
| 3 | Static | ✓ | ✓ | 2 | 5210.7 | 5223.1 |
| 4 | WSLS deterministic | – | – | 0 | 1731797.7 | 1731797.7 |
| 5 | WSLS deterministic | ✓ | – | 1 | 1709749.5 | 1709755.8 |
| 6 | WSLS deterministic | – | ✓ | 1 | 4769.8 | 4776.0 |
| 7 | WSLS deterministic | ✓ | ✓ | 2 | 4769.1 | 4781.5 |
| 8 | WSLS probabilistic | – | – | 2 | 3809.0 | 3821.5 |
| 9 | WSLS probabilistic | ✓ | – | 3 | 3764.8 | 3783.5 |
| 10 | Action kernel only | – | – | 2 | 3984.6 | 3997.0 |
| 11 | Action kernel only | ✓ | – | 3 | 3975.9 | 3994.6 |
| 12 | Action kernel only | – | ✓ | 3 | 3986.1 | 4004.8 |
| 13 | Action kernel only | ✓ | ✓ | 4 | 3977.2 | 4002.2 |
| 14 | Single LR | – | – | 2 | 4256.3 | 4268.8 |
| 15 | Single LR | ✓ | – | 3 | 4108.8 | 4127.6 |
| 16 | Single LR | – | ✓ | 3 | 4256.3 | 4275.0 |
| 17 | Single LR | ✓ | ✓ | 4 | 4109.1 | 4134.1 |
| 18 | Double LR | – | – | 3 | 4246.6 | 4265.3 |
| 19 | Double LR | ✓ | – | 4 | 4096.5 | 4121.5 |
| 20 | Double LR | – | ✓ | 4 | 4245.7 | 4270.6 |
| 21 | Double LR | ✓ | ✓ | 5 | 4096.0 | 4127.2 |
| 22 | Single LR + AK | – | – | 4 | 3716.2 | 3741.1 |
| 23 | Single LR + AK | ✓ | – | 5 | 3675.1 | 3706.3 |
| 24 | Single LR + AK | – | ✓ | 5 | 3718.1 | 3749.3 |
| 25 | Single LR + AK | ✓ | ✓ | 6 | 3677.0 | 3714.4 |
| 26 | Double LR + AK | – | – | 5 | 3715.2 | 3746.4 |
| 27 | Double LR + AK | ✓ | – | 6 | 3673.6 | 3711.0 |
| 28 | Double LR + AK | – | ✓ | 6 | 3717.2 | 3754.6 |
| 29 | Double LR + AK | ✓ | ✓ | 7 | 3675.5 | 3719.1 |
